# Oligomerization processes limit photoactivation and recovery of the Orange Carotenoid Protein

**DOI:** 10.1101/2022.02.04.479168

**Authors:** Elena A. Andreeva, Stanislaw Nizinski, Adjélé Wilson, Matteo Levantino, Elke De Zitter, Rory Munro, Fernando Muzzopappa, Aurélien Thureau, Ninon Zala, Gotard Burdzinski, Michel Sliwa, Diana Kirilovsky, Giorgio Schirò, Jacques-Philippe Colletier

## Abstract

The Orange Carotenoid Protein (OCP) is a photoactive protein involved in cyanobacterial photoprotection, by quenching of the excess of light harvested energy. The photoactivation mechanism remains elusive, in part due to absence of data pertaining to the timescales over which protein structural changes take place. It also remains unclear whether or not oligomerization of the dark-adapted and light-adapted OCP could play a role in the regulation of its energy quenching activity. Here, we probed photo-induced structural changes in OCP by a combination of static and time-resolved X-ray scattering and steady-state and transient optical spectroscopy in the visible range. Our results suggest that oligomerization partakes in regulation of the OCP photocycle, with different oligomers slowing down the overall thermal recovery of the dark-adapted state of OCP. They furthermore reveal that upon non-photoproductive excitation, a numbed-state forms, which remains in a non-photoexcitable structural state for at least ∼0.5 µs after absorption of a first photon.

**Significance Statement:** The orange carotenoid protein (OCP) is a photoactivatable protein involved in cyanobacterial photoprotection. Upon photoactivation, OCP becomes able to quench the excess of energy uptaken by the light-harvesting antennae, thereby evading damage to the cells. It remains unclear, however, what is the exact OCP photoactivation mechanism, and whether or not oligomerization partakes in the regulation of the OCP function. Here, we investigated these issues by combining static and time-resolved (TR) X-ray scattering and optical spectroscopy. Our results show that OCP oligomerizes in both the dark-adapted inactive and light-adapted active states, suggesting a functional role for oligomerization. TR scattering data furthermore reveal that the first large-scale conformational changes associated with OCP photoactivation take place on the µs time scale.

## Introduction

Photosynthetic organisms have evolved to make use of up to 99 % of light harvested energy (1). In most cyanobacteria, the main light harvesting antenna is the phycobilisome (PBS), a soluble complex capable of directly funneling its harvested photon energy into thylakoid-membrane-embedded reaction-centers. In the event of an energy overflow into photosystem II reaction center, charge recombination can occur that will lead to the production of ^1^O_2_, which in turn may damage the photosynthetic apparatus (2, 3). The main function of the soluble two-domain photoactive orange carotenoid protein (OCP) is to quench the excessive energy absorbed by the PBS, enabling the dissipation of this energy into heat. This is accompanied by a decrease in the PBS fluorescence. For its energy-quenching function to be elicited, OCP needs to be photoactivated by the absorption of a blue-green photon, which triggers the transition from the dark inactive orange state (OCP^O^) to the active red state (OCP^R^) (4–11) capable of quenching up to 80 % of the PBS fluorescence (12). As the photoactivation yield of OCP is extremely low (0.2 %) (13, 14), the interaction between PBS and OCP^R^ only occur when irradiance threatens cell survival. Additionally, OCP can also quench ^1^O_2_, and is thus one of the central players in cyanobacterial photoprotection (15, 16). Phylogenetic investigations have allowed classification of OCP sequences into three clades, viz. OCP1, OCP2 and OCPX (17).

Most research endeavors on OCP (including the present work) have been conducted on the *Synechocystis* PCC 6803 OCP1 variant featuring echinenone (ECN) as the functionalizing carotenoid (18, 19). Nonetheless, the first crystallographic structure of a dark-adapted OCP^O^ was from the cyanobacteria *Limnospira (Arthrospira) maxima*, and featured 3’-hydroxy-ECN as the natural pigment (15). The structure revealed that OCP crystallizes as a dimer (Fig. 1*A, B*) wherein each monomer folds into two domains separated by a ∼20 residue linker (Fig. 1*C*). The N-terminal domain (NTD), comprised of residues 1–165, is the effector domain binding to PBS, while the C-terminal domain (CTD) is comprised of residues 190–317 and serves as the regulator of OCP energy-quenching activity (20). The keto-carotenoid pigment binds in a ∼35 Å long hydrophobic tunnel that spans the two domains. The NTD is fully α-helical (αA-αJ), featuring a fold unique to cyanobacteria, whereas the CTD is a seven-stranded β-sheet (β1-β7) sandwiched between two sets of α-helices, viz. αK, αL and αM, on one side (referred to as the F-side) (21), and the terminal αA (residues 1-19; also referred to as the N-terminal extension or NTE) and αN (residues 304-317; also referred to as the the C-terminal tail or CTT), on the opposite side (referred to as the A-side) (21). The αA and αN helices have been shown to play important roles in the regulation of OCP photoactivation and recovery. Stabilization of the OCP^O^ state is achieved at two main interfaces, viz. the NTD/CTD interface, burying 677 Å^2^ of surface area and featuring two strictly-conserved H-bonds (*i.e.,* R155 to E244, N104 to W277; Fig. 1*C*)), and the αA/CTD interface, burying 790 Å^2^ and featuring six H-bonds (buried surface areas calculated from the *Synechocystis* OCP structure) of which two to the C-terminal αN (L16(O)-A308(N) and A18(N)-L307(O)). Thus, the αA helix is one of the main secondary structure elements supporting the stabilization of the OCP^O^ structure. Additionally, the carotenoid buries ∼990 Å^2^ of surface area across the two domains (∼ 545 and 445 Å^2^ in the NTD and the CTD, respectively), establishing two H-bonds with Y201(OH) and W288(Nε1) in the CTD (Fig. 1*C*). The pigment is therefore also essential in stabilizing OCP^O^.

**Figure 1.**
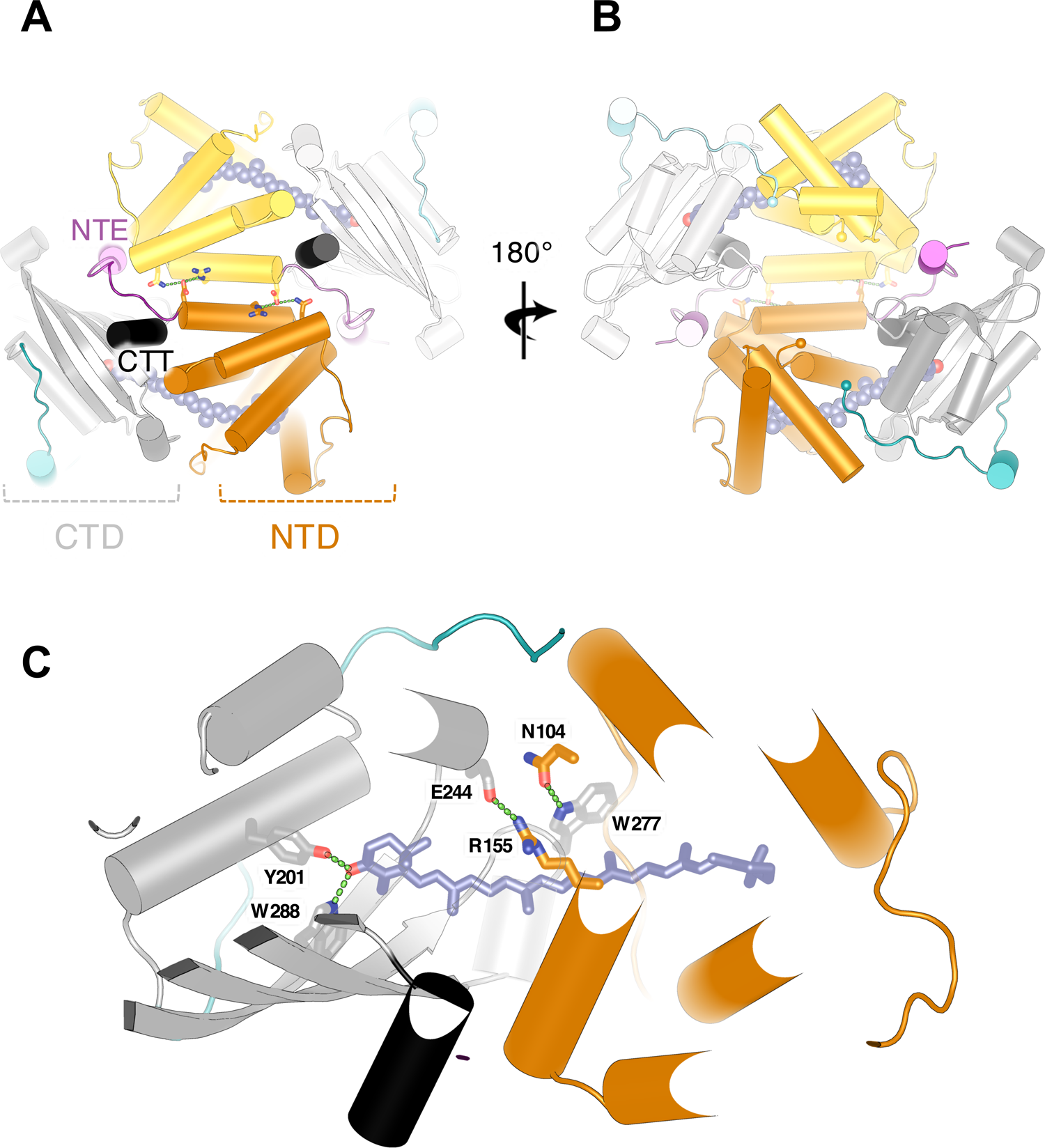
OCP crystallizes as a dimer. (*A, B*) Structure of dark-adapted OCP (OCP^O^) dimer from the cyanobacteria *Synechocystis* PCC 6703 (PDB id: 3MG1). Within the dimer, each OCP monomer features two domains: (i) an N-terminal domain (NTD; coloured in orange) comprising residues 1– 165; (ii) a C-terminal domain (CTD; coloured in grey) comprising residues 193–317. The N-terminal extension (NTE) and the C-terminal tail (CTT) are shown in pink and black, respectively. The two domains are attached by a ∼ 20 residue linker (coloured in cyan). The keto-carotenoid pigment, e.g., echinenone (ECN; coloured in slate and shown as spheres), binds in a ∼ 35 Å long hydrophobic tunnel spanning the two domains and H-bonds to Y201 and W288 in the CTD. The dimerization interface is supported by conserved H-bonds between D19, R27, N134 (shown as sticks). (*C*) Close-up view of dark-adapted OCP (OCP^O^) monomer in the dimer (A, B). The interaction between the NTD and the CTD within the OCP^O^ monomer is supported by two strictly-conserved interactions, *i.e.,* a salt-bridge between R155 and E244, and H-bonds between N104 and W277 (shown as sticks). The two H-bonds of the carotenoid to Y201and W288 (shown as sticks) are also shown (green dashes).

All available crystal structures of OCP^O^ feature a dimer, wherein monomers are associated through contacts between facing αA, αB and αH helices, burying ∼1090 Å^2^ at the dimerization interface (Fig. 1*A, B*). Due to the high concentration of OCP^O^ within crystals (20-30 mM, depending on the crystal structure considered), the prevalence of OCP^O^ in its dimeric form was, until recently, regarded as a crystallization artifact and therefore overlooked (18). Recent results obtained using native mass spectrometry have since shown that OCP^O^ dimers can form in solution at concentrations as low as 3 µM (0.1 mg/ml) (22). The presence of OCP^O^ dimers was also observed by size exclusion chromatography (SEC) and in small-angle X-ray/neutron scattering (SAXS/SANS) experiments (23, 24), which furthermore identified the crystallographic dimer as that which naturally occurs. Also supporting the hypothesis that OCP^O^ dimers could form *in vivo* is the report that the mutation of a single (R27) of the three most-conserved residues (D19, R27, S134) among the 22 making up the dimerization interface (18) results in a constitutively monomeric mutant (10). Further adding to this complexity is the question as to whether or not the light-adapted OCP^R^ can also oligomerize, which has been supported by recent SAXS/SANS data (21, 22) but not by native mass spectrometry (10, 22, 25, 26). Altogether these results raise cogent questions as to if and how oligomerization participates in the regulation of OCP function.

The exact photoactivation mechanism of OCP – meaning, accumulation of OCP^R^, not the mere appearance of a red spectrum achieved already 50 ps post-excitation – remains elusive. Indeed, despite the use of different experimental approaches to probe the OCP^O^ to OCP^R^ transition with a high resolution in either time (27–29) or space (19, 30–32), uncertainties remain regarding the exact sequence of events leading to photoactivation, in part because data are lacking that pertain to the (long) timescales over which (large-scale) structural changes take place. All investigators agree on the fact that upon photon absorption, the carotenoid transitions to an excited S_2_ state, which decays within ∼0.1 ps into multiple ps-lived excited states (S_1_, ICT, S*). Only one of these presumably leads to the minute-lived OCP^R^ state (S* in (27), S_1_ in (14)) after the formation of at least four different intermediates (photoproducts P_1_, P_2_, P_2_’, P_3_) over the ps-µs timescale (27). Thus, the formation of OCP^R^ is mainly limited by picosecond (ps) timescale excited state dynamics, with ∼ 99 % of carotenoids relaxing back to the ground state along non-productive pathways. It was suggested that photoproduct P_1_ is characterized by rupture of the H-bonds between the carotenoid and the protein; P_2_ by a repositioning of the β1-ring in the CTD; and P_3_ by the translocation (12 Å) of the carotenoid into the NTD. The dissociation of helices αA and αN from the A-side of the CTD was suggested to await the millisecond (ms) time scale (25), and to be followed by dissociation of the two domains, yielding the photoactive OCP^R^. This hypothesis has since been supported by SEC and SAXS (30). By specific labeling of accessible carboxylate groups, mass spectrometry further pointed to a signal transduction mechanism whereby disorder is propagated across the CTD β-sheet, upon photoactivation and rupture of H-bonds between the carotenoid and the CTD, resulting in a detachment of helix αA and subsequent dissociation of the dimer through destabilization of helix αB. Direct structural evidence for the existence of these steps is yet lacking. Most recently, experiments carried out on an OCP mutant wherein stabilization of the carotenoid to the NTD is modified (mutation into phenylalanine of W101, W110 and W277) supported the existence of two additional intermediate states between P_3_ and OCP^R^, namely P_M_ and P_X_, proposed to be characterized by dissociation of the αA and αN helices, respectively (29). It yet remains unclear whether these states also exist in the wild-type (wt) protein.

In the present work, we used a combination of mutagenesis, transient spectroscopy in the visible range, and static and time-resolved X-ray solution scattering to clarify (i) the oligomerization states of OCP^O^ and OCP^R^ in solution, (ii) the timescale over which large-scale structural changes associated to photoactivation take place, and (iii) whether or not pulsed-illumination permits the formation of OCP^R^. By conducting static SAXS experiments on the wild-type OCP and on a stable monomeric R27L mutant (10), wherein the conserved R27-D19 salt-bridge at the dimerization interface is suppressed, we obtained confirmation that both OCP^O^ and OCP^R^ can oligomerize in solution, suggesting that oligomerization could play a role in the regulation of OCP activity. Using TR-SAXS, we obtained evidence that the ‘red’-shifted state generated by pulsed illumination, which forms and decays within ∼10 µs and ∼10-200 ms (depending on the his-tag location), respectively, differs from the OCP^R^ state accumulated under stationary illumination conditions, which forms and decays within ∼1 ms and ∼1-30 min (depending on concentration), respectively. Our data furthermore reveal that upon a non-productive pulsed photoexcitation – *i.e.* in the 99.7 % of the cases where OCP^R^ does not form – OCP remains in a non-photoexcitable structural state for at least ∼ 0.5 µs, despite the carotenoid returning back to the electronic ground state within tens of ps and the protein featuring a spectrum characteristic of the OCP^O^ state (27).

## Materials and Methods

### Protein expression and sample preparation

OCP expression and extraction were carried out as described in Gwizdala et al. (8) and Bourcier de Carbon et al. (33). Briefly, expression in the holo form of the N-terminally his-tagged wild-type OCP (OCP_wt-Ntag_) and the N-terminally his-tagged monomeric mutant (OCP_R27L-Ntag_) was achieved by respective transformation of the *pCDF-NtagOCPSyn* or *pCDF-NtagOCPSynR27L* plasmids into ECN-producing BL21 (DE3) *E. coli* cells – *i.e.* cells previously transformed with the pAC-BETA and pBAD-CrtO plasmids described in Bourcier de Carbon et al. (31). Expression in *Synechocystis* cells of the holo form of C-terminally his-tagged wild-type OCP (OCP_wt-Ctag_) was achieved using the protocol described in Gwizdala et al. (8). The proteins were purified in three steps including affinity chromatography on a Ni-NTA column (Ni-Probond resin, Invitrogen), hydrophobic chromatography on a phenyl-sepharose column (HiTrap Phenyl HP, GE Healthcare) and size exclusion chromatography on an analytical HiLoad 16/60 Superdex 75 (HiLoad 16/600 Superdex 75 pg, Sigma Aldrich). Proteins were eluted from the latter at a flow rate of 1 ml/min using a 50 mM Tris-HCl pH 7.4, 150 mM NaCl buffer. To assert the molecular weight of eluted species, the column was calibrated beforehand using the following standard proteins: γ-globulin (158 kDa), ovalbumin (44 kDa), myoglobin (17 kDa) and vitamin B12 (1.35 kDa). The purity of both OCP variants were assessed by 12% SDS-PAGE electrophoresis (Fig. S11). OCP concentrations were estimated from the absorbance signal at 490 nm, using the extinction coefficient of 63000 M^-1^ cm^-1^ proposed by Sluchanko et al. (34). We note that an uncertainty remains as to the exact extinction coefficient that should be used to derive OCP concentration from absorbance measurement. Protein concentration is nowadays generally estimated using the absorbance signal at 280 nm and a calculated extinction coefficient (e.g. obtained from https://web.expasy.org/cgi-bin/protparam/protparam (35, 36)), which in the case of *Synechocystis* OCP amounts to 34 659 M^-1^.cm^-1^. This value agrees well with the extinction coefficient at 490 nm proposed by Sluchanko et al., as the ratio between the 490 and 280 nm peaks is ∼2 for the pure protein, but it neglects the potential absorption of the ketocarotenoid pigment at 280 nm. Thus far, most of the papers published on OCP have accordingly used an extinction coefficient of 110 000 M^-1^ cm^-1^ at 490 nm, based on the debatable assumption that the ketocarotenoid pigment has similar absorption spectrum when bound to OCP or dissolved in organic solvents. Indeed, if there is no absorbance of the carotenoid at 280 nm, then one expects a ratio of ∼3 between the absorbances peak at 490 nm and 280 nm. But if we consider absorption of the carotenoid at 280 nm, and assume the same extinction coefficient when bound to the protein to the protein scaffold or dissolved in organic solvents – *i.e.* roughly 15 % of the absorption peak at 490 nm (measured by us as well as others) – then the measured ratio of ∼2 between the absorbance at 490 and 280 nm can be rationalized since A_280 nm_ = (0.33+0.15) *A_490 nm_ = 0.48* A_490 nm_. At present, we cannot favor either hypothesis. However, as our current aim is to report on oligomerization processes, we believe it is preferable that we underestimate – rather than overestimate – dissociation constants. Hence, the concentrations reported in the present paper were estimated using the conservative extinction coefficient of 63 000 M^-1^ cm^-1^ (34). The actual concentrations and dissociation constants could yet be 1.7 times lower if the ketocarotenoid absorbs at 280 nm and the real extinction coefficient of OCP at 490 nm is 110 000 M^-1^.cm^-1^.

### Static SAXS measurements

Static X-ray scattering data were collected at 21°C on an EigerX-4M detector at the SWING beamline of the Soleil Synchrotron (Saint-Aubin, France). X-ray solution scattering signals in the SAXS region (q = 0.02 - 0.5 Å^-1^) were collected for OCP_wt-Ctag_ and OCP_R27L-Ntag_ at various concentrations using a monochromatic (double Si (111) monochromator) X-ray beam centered at 12 keV and an EigerX-4M (Dectris) detector. Two sets of measurements were carried out, before and after 30 min irradiation of OCP solutions with a 430 nm LED source (∼ 500 mW/cm²), allowing collection of X-ray scattering signals in the dark-adapted (OCP^O^) and light adapted (OCP^R^) states, respectively. For OCP_wt-Ctag_, data were acquired at 0.7, 3.5, 10.5 and 23 mg/ml for the OCP^O^ state and at 0.7, 3.5 and 10.5 mg/ml for the OCP^R^ state, while for OCP_R27L-Ntag_, data were recorded at 0.7, 3.5, 10.5 and 14 mg/ml for the OCP^O^ state and at 0.7, 3.5 and 10.5 mg/ml for the OCP^R^ state. For each concentration, 30 X-ray scattering signals (each registered after 990 ms of X-ray exposure) were collected with 30 analogous signals of the buffer before and after each protein measurement. Each 2D signal was converted to a 1D scattering profile by azimuthal integration using the Foxtrot-3.5.2 software available at the SWING beamline. Corresponding scattering profiles were averaged and protein signals were obtained by subtraction of buffer scattering. The distance distribution functions were computed using GNOM (37) and the data collection parameters reported in Table S1 were determined from the reduced data using relevant programs from the ATSAS (37, 38) suite. Data were also examined using RAW (39), which enabled extraction of Porod (V_p_) and correlation volumes (V_c_), and an independent estimate of the molecular weight of scattering (Tables S2 and S3). Low-resolution molecular envelopes were computed using the ATSAS reconstitution tool DAMMIF (37) (Fig. S1 and Table S4). To construct models for OCP^R^ dimer and higher-order oligomers, we used the published structures of paralogs of the isolated N-terminal and C-terminal OCP domains, viz. the crystal structures of *Anabaena Nostoc* PCC 7120 and *Fremyella diplosiphon* HCP (pdb ids: 6MCJ and 5FCX, respectively) (40, 41) and of *Anabaena Nostoc* PCC 7120 CTDH (pdb id: 5FEJ) (21), respectively. Our modeling strategy is presented in the Results section. The presence of oligomers larger than dimers was inferred from the superimposition of OCP^R^ models to the DAMMIF envelopes (Fig. S1) and validated by SREFLEX (42) normal mode analysis (Fig. S2 and Table S5). The final data and models for each OCP variant were deposited in the SASBDB (43). The accession IDs are available in Table S5.

### Spectroscopic monitoring of the OCP^R^ to OCP^O^ thermal recovery after accumulation of OCP^R^ by a prolonged continuous illumination

Thermal recovery was investigated as a function of concentration on the three OCP variants, using a JASCO V-630 UV/Vis spectrophotometer (Easton, USA). Purified proteins in OCP_R27L-Ntag_ and OCP_wt-Ctag_ were assayed at 0.1, 1.7, 3 and 16 mg/ml, and OCP_wt-Ntag_ at 0.1, 3, 10 and 16 mg/ml. Specifically, the absorption at 467 nm was monitored at room-temperature (21 °C) in the dark following 30 minutes of continuous blue light irradiation with a 430 nm LED light collimated to 1 cm (∼ 500 mW/cm²) to maximize accumulation of OCP^R^ and reproduce the conditions used in the static SAXS experiments. Measurements at 0.1 mg/ml were carried out in a PMMA cuvette with 1 cm pathlength; those at 1.7, 3 and 10 mg/ml in a quartz cuvette with 0.1 cm pathlength; and that at 16 mg/ml in an Infrasil cuvette with 0.005 cm pathlength. Stability of the proteins over the course of experiments was verified by comparison of their steady state absorbance spectra before illumination and after illumination and recovery, respectively (Fig. S6).

To facilitate comparison of data collected at various concentrations, the absorption difference at 467 nm was normalized using A_Norm_ = [A(*t*) - A(*t_0_*)]/[A(*t_max_*) - A(*t_0_*)], where A(*t*), A(*t_0_*) and A(*t_max_*) are the absorption values measured at a generic time *t* after switching off the 430 nm light; at time t_0_ (*i.e.,* just after illumination at 430 nm was switched off, when the concentration of OCP^O^ is at its minimum); and at the time *t_max_* when the starting OCP^O^ state has recovered, respectively. The non-linear least-squares optimization and parameter extraction were performed using LMFIT (44). After individual fits of each curve data pointed to the necessity to use at least three exponential components to fit the recovery data, we opted for a per-OCP variant global-fitting of the normalized absorption data. For each sample, the recovery data measured at four concentrations were jointly fitted using either a triple or a quadruple exponential function,

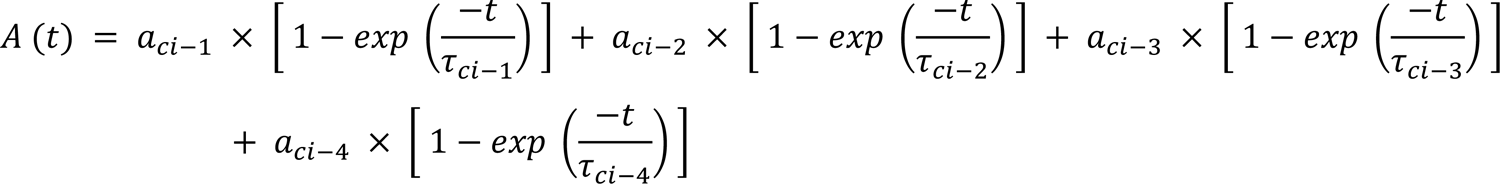

where a**_ci_**_-1_, a**_ci_**_-2_, a**_ci_**_-3_ and a**_ci_**_-4_ are concentration-dependent pre-exponential factors, τ**_ci-_**_1_, τ**_ci_**_-2_, τ**_ci_**_-3_ and τ**_ci_**_-4_ are process-dependent lifetimes, and a**_ci_**_-4_ is fixed to zero in the case of a triple exponential fit. A better fit was obtained with four exponential components for the OCP_wt-Ctag_ recovery data, but improvements were marginal for the data collected on OCP_wt-Ntag_ and OCP_R27L-Ntag_, with amplitudes for the fourth component consistently inferior to 0.1 (see Table S2 and Table S3). Hence, we retained results from the four-exponential fit for OCP_wt-Ctag_, but from the three-exponential fit, for the two N-tagged variants. We nonetheless present results for both types of fits in Fig. 4, Supplementary Fig. S7 and S8, and Tables S6 and S7.

### Spectroscopic monitoring of the spectral evolution after ns-pulsed excitation at 470 nm

Measurements were performed using the custom apparatus described previously (45). OCP^O^ photo-excitation was achieved using a 470 nm nanosecond laser delivering 5 mJ energy per 8 ns pulse at a repetition rate of 0.05 Hz. The probing beam was filtered using a bandpass filter, so that only 440 nm (FWHM: 9 nm) or 565 nm (FWHM: 12 nm) light passed through the sample, avoiding putative probe-induced effects from the white spectrum of the xenon lamp. Scattered pump light was reduced by a notch filter set installed upstream of the monochromator which precedes the photomultiplier tube. All measurements were carried out at a sample absorbance of ∼ 0.7 at 470 nm. For the lowest concentration, a 10 x 10 mm cuvette was used, which was set with a right-angle configuration between the pump and the probe beams. For higher concentrations, 2 mm and 500 µm flat cuvettes were used, affording to increase the concentration whilst keeping the absorbance in the optimal range. Use of these cuvettes required a quasi-collinear (≈ 5° angle) configuration between the pump and probe beams. Measurements were performed at different temperatures ranging from 8 to 36° C. Stirring was not applied to avoid sample displacement from the probed volume at longer time delays. All experiments were repeated 100 times on each of 4 partially-overlapping time-windows, altogether covering the 50 ns − 1 s time-range. Recorded data were then merged and projected on a logarithmic grid. Stability of the protein was checked by its steady-state absorbance after each experiment. The quantum yield formation of photo-products was determined using ruthenium as actinometer (34, 46) and molar absorption coefficient of OCP^O^ at 490 nm = 63000 cm^-1^.M^-1^ (34).

### Time-resolved X-ray scattering

Time-resolved SAXS/WAXS experiments were performed at the ID09 beamline of the European Synchrotron Radiation Facility (ESRF, Grenoble, France) (47). X-ray solution scattering signals in the WAXS region (q = 0.03 – 2.5 Å^-1^) were collected at 23 mg/ml (620 µM) for OCP_wt-Ntag_ and OCP_wt-Ctag_, using a pink polychromatic X-ray beam centered at 15 keV (∼0.3 keV bandwidth, achieved using a multilayer monochromator) and a Rayonix MX170-HS detector placed 350 mm from the sample. The protein samples, OCP_wt-Ntag_ and OCP_wt-Ctag_, were photoactivated with laser pulses from two different laser systems: either an EKSPLA NT342B laser or a COHERENT Evolution laser. While the first laser is a tunable laser generating 5 ns duration pulses (full width half maximum, FWHM) at a maximum repetition rate of 10 Hz, the second laser runs at 1 kHz, has a fixed wavelength (527 nm) and has a pulse duration that can be modulated by changing the current of its diode pump. The use of the COHERENT Evolution laser allowed us to perform a set of experiments with pulse durations longer than those of the EKSPLA laser (in particular, experiments with 300 or 450 ns long pulses were performed); moreover, it allowed us to excite the protein sample with either a single laser pulse or a burst of several laser pulses (up to 30) at 1 kHz. For each experiment, the required number of photoexcitation laser pulses was extracted from the train of pulses generated by the Evolution laser by means of a pair of synchronized fast mechanical shutters (Lambda SC, Sutter). By changing the relative delay of the first shutter opening and of the last shutter closing, it was possible to cleanly select either a single laser pulse or several. In all experiments, the laser beam was focused with cylindrical lenses to an elliptical spot approximately 2.5 × 0.25 mm^2^ (FWHM) and the energy was regulated so as to have an energy density of ∼ 3 mJ/mm^2^ at the sample position, corresponding to ∼ 75 absorbed photons per chromophore.

To maximize the overlap between the pump- and probe-illuminated volumes, an orthogonal pump– probe geometry was employed, with the X-ray beam (0.06 × 0.1 mm^2^, FWHM) hitting the sample capillary 0.3 mm from its edge. In order to both minimize X-ray radiation damage and allow the recovery of the OCP^O^ state, the protein solution (∼ 5 ml) was circulated with a peristaltic pump (Gilson Minipuls 3) through a 2 mm X-ray quartz capillary in a closed loop. During the flow, most of the sample was contained in a small reservoir kept in the experimental hutch, thermalized at 22 °C. The flow speed was set according to both the optical-laser pump – X-ray probe time delay and the repetition rate, allowing the sample to be kept in the pump–probe intersection area during a pump–probe sequence while refreshing it between two consecutive pump pulses. Single X-ray pulses (5, 10 or 30 µs depending on the time delay) were selected from the synchrotron pulse train by means of a high-speed chopper and a millisecond shutter. It was verified that the scattering signal obtained with 5-30

µs X-ray pulses is qualitatively similar. Laser-off (dark) signals were also acquired with the X-ray pulse arriving 50 ms before the laser pulse and used as a reference to compute the difference profiles after normalization to the water peak (2 - 2.2 Å^−1^) (47). Signals were azimuthally integrated and the peak of the undulator spectrum (∼ 0.83 Å^−1^) was used as the reference to convert the scattering angle to the momentum transfer q. Up to 30 scattering profiles per time delay were acquired and averaged to improve the signal-to-noise ratio. Independent TR-WAXS measurements on a solution of 4-amino-1,1′-azobenzene-3,4′-disulfonic acid monosodium salt with an optical density comparable to the OCP sample were used to correct the water heating scattering signal of the collected OCP data (48). TR-WAXS was also used to follow the OCP^R^ to OCP^O^ thermal recovery with a direct structurally sensitive technique. In particular, X-ray scattering profiles were collected over 30 minutes from a sample of OCP_wt-Ctag_ that had been exposed for 30 minutes to continuous illumination by a 430 nm LED light.

The TR-X-ray scattering dataset was analyzed by a singular value decomposition (SVD) (49) using a custom-written Python script (Fig. S12). The time-dependent difference profiles form a j × k matrix, with j the number of q values and k the number of time delays. The SVD algorithm calculates the matrices U and V and the vector S, so that A = U × S × V^T^. The columns of matrix U are called left singular vectors, or basis patterns, the rows of V^T^ are called right singular vectors, or amplitude vectors, and the elements of S are called singular values. The basis patterns are ordered following the high-to-low sorting of singular values. The SVD analysis of kinetic data provides a subset of time-independent patterns containing the relevant information out of the random noise. The dataset can be then reproduced as a linear combination of such time-independent patterns. Inspection of the shape of basis patterns, amplitude vectors and singular values together with the autocorrelation analysis of both basis patterns and amplitude vectors (50) indicates that only one component contains significant information. Hence, a two-step model (with growing and decaying exponentials) was used to fit the evolution of the integrated time-resolved X-ray scattering profile.

## Results

### Soluble OCP^O^ can form dimers in solution

We used small-angle X-ray scattering to investigate the oligomerization state of dark-adapted OCP (OCP^O^) in solution. For this purpose, we worked on two proteins: (i) a wild-type OCP, functionalized by ECN and featuring a six-histidine tag (6Xhis-tag) at the C-terminus (OCP_wt-Ctag_); and (ii) a R27L monomeric mutant, functionalized by ECN and featuring a 6Xhis-tag at the N-terminus (OCP_R27L-Ntag_) (10).

Data were collected on dark-adapted OCP_wt-Ctag_ and OCP_R27L-Ntag_ at increasing protein concentrations, informing on the OCP^O^ state. The buffer-subtracted scattering profiles of dark-adapted OCP_wt-Ctag_ and OCP_R27L-Ntag_ are shown in Fig. 2*A* and 2*B*. For OCP_R27L-Ntag_ at all concentrations, the I(0)/c and the radii of gyration (Rg) derived from the Guinier region are similar (Fig. 3*A* and 3*B*), and the derived pairwise distance distribution functions P(r) overlay (Fig. 2*C*). Accordingly, the low-resolution molecular envelopes calculated from the dark-adapted OCP_R27L-Ntag_ data collected at 0.7, 3.5 and 10.5 mg/ml (Fig. S1) are comparable, both matching the crystallographic OCP^O^ monomer. Hence the OCP^O^_R27L-Ntag_ sample, wherein the conserved D19-R27 salt bridge is suppressed, remains in the same monomeric state at all tested concentrations and therefore offers a good control for monomeric OCP^O^. Contrastingly, examination of the P(r) for dark-adapted OCP_wt-Ctag_ samples (Fig. 2*C*) reveals an increase in pairwise distances when the concentration exceeds 3.5 mg/ml. Accordingly, an increase is observed in the I(0)/c (Fig. 3*A*) and the radii of gyration derived from the Guinier region (Fig. 3*B*). These three observations indicate that the average size of dark-adapted OCP_wt-Ctag_ increases with concentration. The low-resolution molecular envelopes calculated by DAMMIF (37) from the dark-adapted OCP_wt-Ctag_ data are consistent with the presence of a predominant OCP^O^ dimer (Fig. S1) at concentrations ≥ 3.5 mg/ml, and a mixture of monomers and dimers at 0.7 mg/ml. It is of important note that the above assignments of molecular envelopes to monomers and dimers agree with the molecular weight estimates from I(0) (*i.e*., the intensity extrapolated at q=0) and Porod (Vp) and correlation (Vc) volumes (see Table S2).

**Figure 2.**
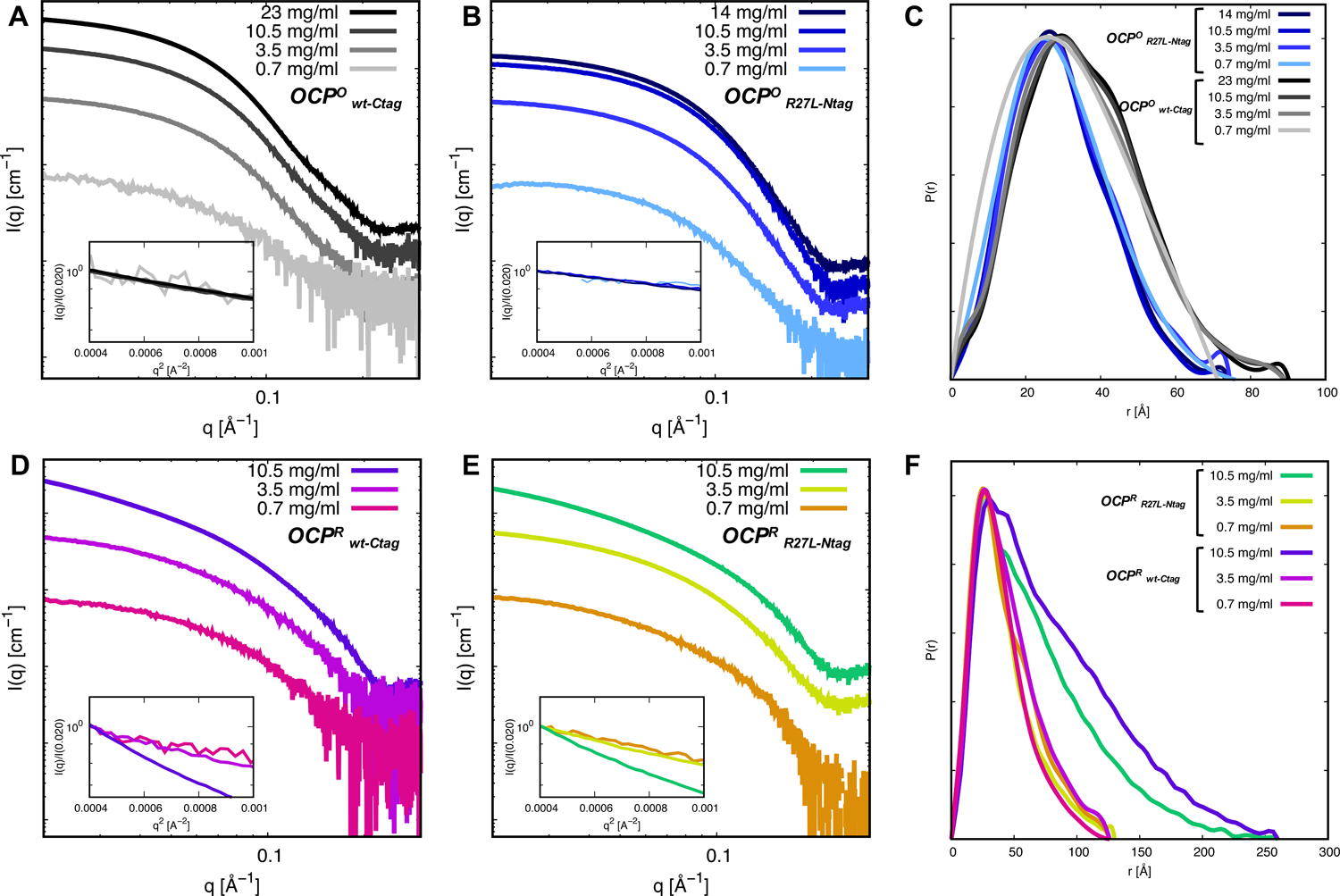
Static X-ray scattering reveals changes in the OCP structure and oligomerization state upon illumination and increase of concentration. (*A, B)* X-ray scattering profiles of dark-adapted OCP^O^_wt-Ctag_ (*A*) and OCP^O^_R27L-Ntag_ (*B*) and their corresponding overlays in the Guinier region (insets) at various concentrations. *(C)* Normalized pair distance distribution function P(r) of dark-adapted OCP^O^_wt-Ctag_ and OCP^O^_R27L-Ntag_ at various concentrations. (*D, E*) X-ray scattering profiles of light-adapted OCP^R^_wt-Ctag_ (*D*) and OCP^R^_R27L-Ntag_ (*E*) at various concentrations. (*F*) Normalized pair distance distribution function P(r) of light-adapted OCP^R^_wt-Ctag_ and OCP^R^_R27L-Ntag_ at various concentrations.

**Figure 3.**
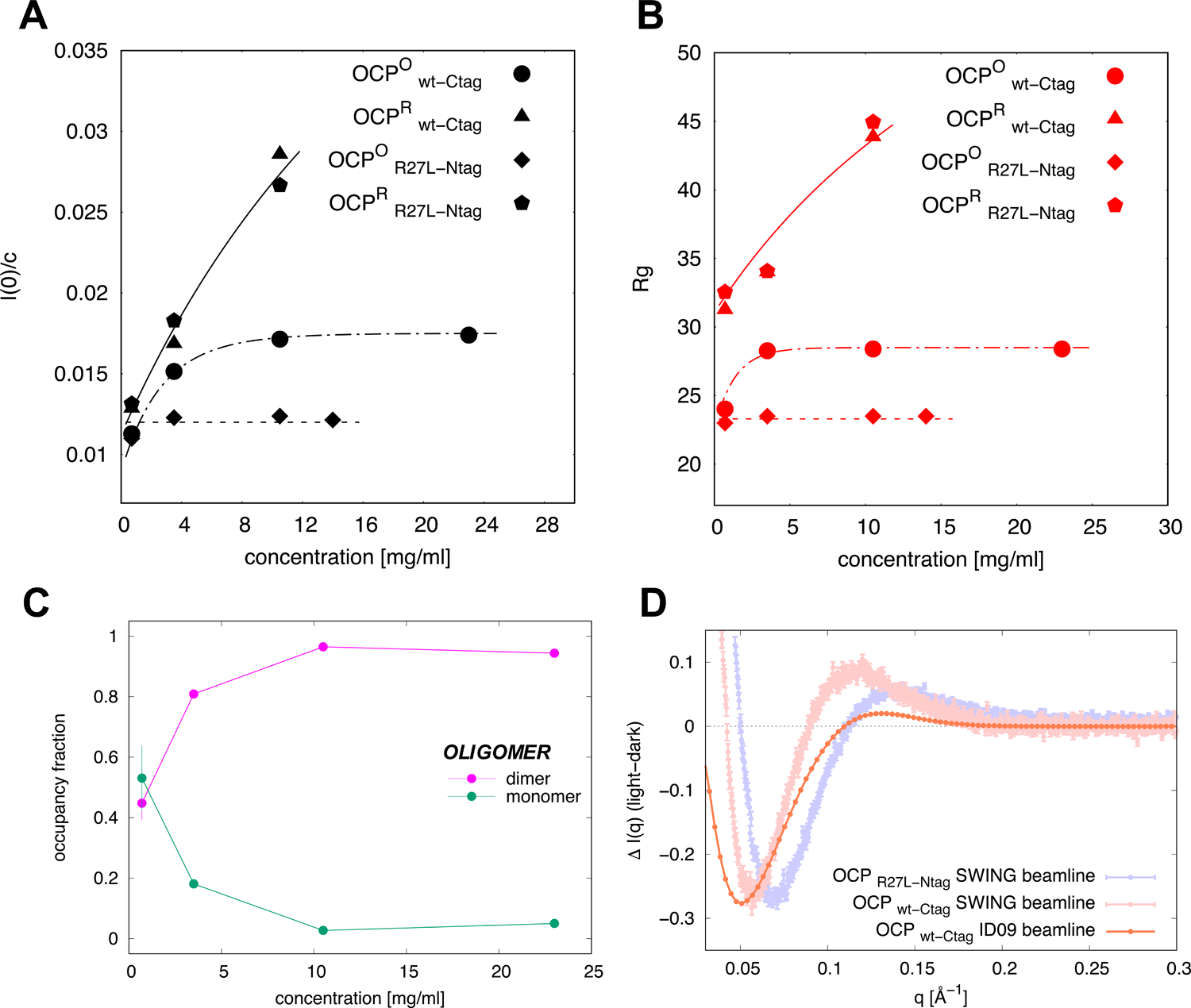
Probing OCP oligomerization in the dark and light-adapted states. (*A*) Evolution of I(0)/c for dark-adapted and light-adapted OCP_wt-Ctag_ and OCP_R27L-Ntag_ as a function of concentration (lines provide a guide to the eye). (*B*) Evolution of the radii of gyration (Rg) for dark-adapted and light-adapted OCP_wt-Ctag_ and OCP_R27L-Ntag_ as a function of concentration (lines provide a guide to the eye). (*C*) Estimation of the relative populations of OCP^O^_wt-Ctag_ monomer and dimer as a function of protein concentration using OLIGOMER. Error bars are reported when larger than the symbol size. (*D*) Light minus dark SAXS difference profiles Δ*I*(q) for OCP_R27L-Ntag_ and OCP_wt-Ctag_ at 10.5 mg/ml (signal recorded on the bio-SAXS SWING beamline, Synchrotron SOLEIL) and for OCP_wt-Ctag_ at 23 mg/ml (signal recorder at ID09 beamline, ESRF). Error bars are reported when larger than the symbol size. Shifts on the position of the positive and negative peaks are likely due to the different illumination protocols and concentrations of the protein. The larger beamstop and shorter sample to detector distance at ID09, likely also influence the signal at low q values. Nonetheless, the figure shows that depletion of the OCP^O^_wt-Ctag_ state can be achieved in the circulating conditions imposed at ID09, although the smaller positive peak suggests presence of less OCP^R^.

**Figure 4.**
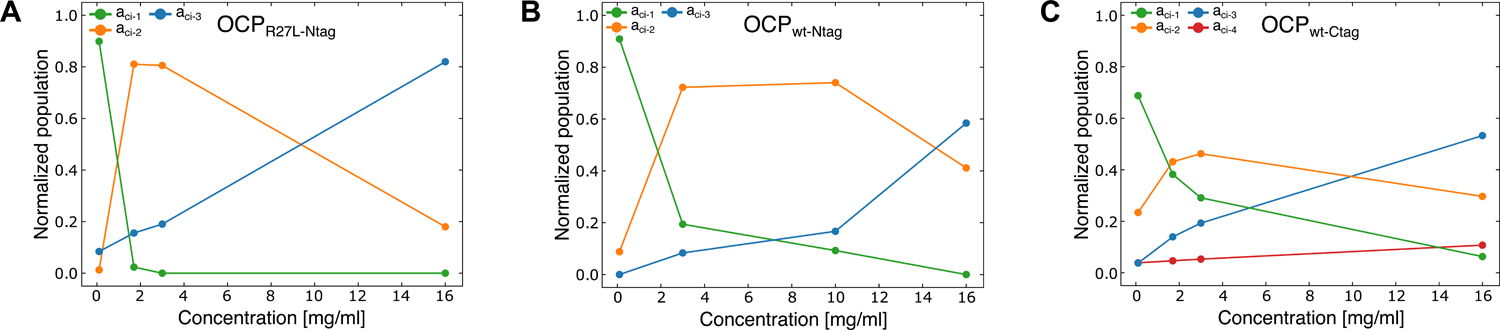
Kinetic analysis of OCP thermal recovery data at various concentrations suggests that oligomerized OCP^R^ reverts more slowly to the dark-adapted state. Plots of the pre-exponential factors retrieved from fits of thermal recovery kinetics monitored by following changes in absorbance at 467 nm (see Fig. S7 and S8, Table S6 and Table S7). (*A*) OCP_R27L-Ntag_, (*B*) OCP_wt-Ntag_ and (*C*) OCP_wt-Ctag_. The apparent lifetime of OCP^R^ state is increased in dimers and higher-order oligomers, suggesting that oligomerization stabilizes the light-adapted state.

We used the program OLIGOMER (38) to estimate the relative abundance of monomers and dimers in the dark-adapted OCP_wt-Ctag_ samples prepared at different concentrations (Fig. 3*C*). Calculated X-ray scattering curves for dimeric and monomeric OCP^O^ were generated from the crystal structure of *Synechocystis* PCC 6803 OCP (pdb id: 3MG1) (18) (Fig. 1) and used for deconvolution of the experimental scattering profiles measured at increasing concentrations (see Materials and Methods). The fit suggests that at 0.7 mg/ml (∼ 20 µM), dark-adapted OCP_wt-Ctag_ is present as a 1:1 mixture of monomers and dimers. The latter attain dominance at ∼3.5 mg/ml (∼ 100 µM) with a dissociation constant of c.a. 0.5 mg/ml (∼ 14 µM). Of course, the accuracy of this value is limited, given the low resolution of method used and the coarseness of the titration data.

### Illumination induces a dissociation of the dark OCP^O^ dimer and the formation of OCP^R^ dimers and higher-order oligomers

To provide details on the large-scale conformational changes accompanying the OCP^O^ to OCP^R^ transition, including possible change(s) in oligomeric state, we also collected X-ray scattering data on light-adapted OCP_wt-Ctag_ and OCP_R27LNtag_ at 0.7, 3.5 and 10.5 mg/ml (Fig. 2*D* and 2*E*). To assess the time required for the accumulation of the OCP^R^wt_-Ctag_ and OCP^R^_R27L-Ntag_ states, we carried out preliminary pre-illumination experiments at ESRF-ID09. A highly-concentrated protein solution (30 mg/ml) of OCP_wt-Ctag_ was used, which was presented to the X-ray beam in a 2 mm thick glass capillary. Firstly, a reference profile was collected on the dark-adapted protein, after which the protein was illuminated by exposure to a 430 nm LED. The distance between the light source and the sample was such that the diameter of the illumination spot was ∼ 1 cm at sample position. Scattering profiles were collected every 15 s, each with 750 ms X-ray exposure, and two illumination powers were consecutively tested, viz. 250 and 500 mW. Only after tens of minutes of illumination at 500 mW was accumulation of OCP^R^ found to reach a plateau, in our experimental conditions (Fig. S4). Therefore, all subsequent experiments aimed at characterizing the active OCP^R^ state involved ∼ 30 minutes illumination by the 500 mW continuous LED source emitting at 430 nm prior to data collection.

A concentration series was recorded for OCP_R27LNtag_ and OCP_wt-Ctag_ in their light adapted states, at the SWING bio-SAXS beamline (Synchrotron SOLEIL) (Table S1). Absolute X-ray scattering profiles of light-adapted OCP^R^_wt-Ctag_ and OCP^R^_R27L-Ntag_ (Fig. 2*D* and 2*E*, respectively) strongly differ from those obtained at the same concentrations for the corresponding proteins in their dark state (Fig. 2*A* and 2*B,* respectively). The profiles exhibit a dependence on protein concentration, with the low-resolution signal increasing as the latter augments. Accordingly, the Rg of light-adapted OCP^R^_R27L-Ntag_ and OCP^R^_wt-Ctag_ increase with concentration, starting from 32.5 and 29 Å at 0.7 mg/ml, respectively, and reaching 44.9 and 43.8 Å at 10.5 mg/ml, respectively (Fig. 3*B* and Table S3). These values are far larger than those observed for the dark-adapted counterparts, but notably similar for the two proteins at each of the three tested concentrations (0.7, 3.5 mg/ml and 10.5 mg/ml) (Tables S1, S2 and S3). The increase in size of OCP_R27L-Ntag_ and OCP_wt-Ctag_ upon illumination, the dependence of this size to the concentration, and the similarity between OCP^R^_R27L-Ntag_ and OCP^R^_wt-Ctag_ at comparable concentrations can also be deduced from the derived P(r) (Fig. 2*F*), from the I(0)-derived molecular weight and from Vp and Vc (Table S3). Thus, our results not only indicate an overall expansion of the protein upon illumination, but also a change in the OCP^R^ quaternary structure as a function of concentration. Similar results (notwithstanding a missing data point for 3.5 mg/ml concentration) were obtained with OCP^R^_wt-Ntag_ (not shown). For each protein, a light minus dark difference scattering profile ΔI(*q*) was calculated at the highest tested concentration (10.5 mg/ml), offering a signature of the change in scattering signal upon illumination (Fig. 3*D*). The ΔI(*q*) for OCP_wt-Ctag_ is characterized by the presence of a negative peak at ∼ 0.05 Å^-1^ and a positive peak at ∼ 0.12 Å^-1^, whereas for OCP_R27L-Ntag_, the negative peak is observed at ∼ 0.07 Å^-1^ and the positive peak at ∼ 0.14 Å^-1^ (Fig. 3*D*). Since OCP^R^_R27L-Ntag_ and OCP^R^_wt-Ctag_ adopt similar states at 10.5 mg/ml, the displacement of negative and positive peaks can be proposed to emanate solely from the difference between dark-adapted structures, viz. OCP^O^_R27L-Ntag_ and OCP^O^_wt-Ctag_.

We used DAMMIF to compute the low-resolution molecular envelopes of OCP^R^_R27L-Ntag_ and OCP^R^_wt-Ctag_ at the three tested concentrations (0.7, 3.5 mg/ml and 10.5 mg/ml). At 0.7 and 3.5 mg/ml (20 and 100 µM),

We attempted to model the OCP^R^ structures assuming that:

i. OCP^R^ dimers and higher-order oligomers are related, *i.e.* the latter form from the coalescence of two OCP^R^ dimers;
ii. interactions between OCP^R^ monomers in a dimer and between OCP^R^ dimers in a higher-order oligomer should be symmetrical, *i.e.* they should involve the same interface in the two interacting proteins;
iii. functional OCP^R^ dimers should assemble via their CTD so that their two NTD are exposed and free to interact with PBS via R155 (51); consequently, OCP^R^ higher-order oligomers would form from an interaction between two NTD in the OCP^R^ dimer;
iv. the interfaces allowing the assembly of OCP^R^ dimers and higher-order oligomers should not involve R27 (and by extension, the dark dimerization interface contributed by helix αB) since the R27L mutant – unable to form OCP^O^ dimers – is capable of forming OCP^R^ dimers and higher-order oligomers;
v. rather the CTD-CTD and NTD-NTD interfaces involved in OCP^R^ oligomerization should feature residues that are not exposed in the dark OCP^O^ dimer, nor present at symmetrical crystal contacts in the OCP^O^ structure, otherwise OCP^O^ higher oligomers would form in the dark-adapted state. Presumably, residues involved in OCP^R^ oligomerization would be present at the NTD-CTD interface in the dark OCP^O^, echoing the earlier proposal of Moldenhauer et al. 2017 (52) and Muzzopappa et al. (10, 52, 53) that the interface between CTD domains in an OCP^R^ dimer should overlap with the CTD residues that interacts with the NTD in OCP^O^;
vi. the interfaces allowing the assembly of OCP^R^ dimers and higher-order oligomers may already have been observed in available crystal structures of OCP and isolated OCP domains, e.g. in the structure of the isolated holo-NTD of wild-type OCP (19), or in those of NTD and CTD paralogs, respectively coined helical carotenoid proteins (HCP) and CTD-homologues (CTDH) in the literature (21, 40, 41).

Indeed, the study of Harris et al. (21) has shown that the apo-*Anabaena* CTDH – a homologue of the CTD of OCP involved in carotenoid transport and wherein the C-terminal helix is not apposed onto the β-sheet – is capable of forming two types of dimers, associating either through back-to-back (A-type) or head-to-head (F-type) interactions between two CTDHs. In the F-type dimer, wherein CTDH residues equivalent to those involved in carotenoid binding in the CTD of OCP^O^ face one another reconstituting a carotenoid tunnel, were proposed to be involved in carotenoid uptake (21). In absence of the carotenoid, however, the A-type interface can form between CTDH molecules, and it was demonstrated by SAXS that the isolated apo-CTD of OCP^O^ dimerizes by this interface. Golub et al. therefore proposed that the two monomers in a OCP^R^ dimer interact by this interface (10, 24). Like them, we hypothesize that OCP^R^ dimers form through interaction at this interface, which fulfills our above hypotheses for a sound modeling of OCP^R^ oligomers. It was further verified that only the A-type interface can fit the extended molecular envelope derived from our data. Indeed, the F-type interface would result in a “Z”-shaped molecular envelope. In the dark OCP^O^, interaction by the A-type interface is prevented by the presence of helices αN (CTT, corresponding to residues 304-317) and αA (or NTE, corresponding to residues 1-17) which appose aside on the external face of the β-sheet but detach upon photoactivation. These helices were therefore considered as disordered in OCP^R^ and not included in our OCP^R^ dimer model. To further avoid clashes and account for the known exposure, in OCP^R^, of the face of the NTD that interacts with the CTD in OCP^O^, the NTD was rotated by 180° and translated by 10 Å. The OCP^R^ dimer was then used as a starting point for modeling a tetramer, by assembly of two NTD in a fashion reminiscent to that observed in the HCP crystal structures from *Anabaena* Nostoc PCC 7120 and *Fremyella diplosiphon* (pdb ids: 5FCX and 6MCJ, respectively) (40, 41). In these, two molecules are found in the asymmetric unit, forming a dimer assembled through a symmetrical interface that features the residue equivalent to OCP-R155 at its center. Hence, in our model, the two dimers in an OCP^R^ tetramer would interact by the same face known to bind the PBS, leaving two NTD available for binding to the latter. This tetramer model fits the *ab initio* envelopes computed for OCP^R^_R27L-Ntag_ and OCP^R^_wt-Ctag_ at 10.5 mg/ml (300 µM) (Fig. S1) but uncertainties remain as to whether only tetramers or only trimers or a mixture thereof coexists in our experiments.

### Kinetics of thermal recovery after accumulation of OCP^R^ by continuous illumination

The recoveries of the OCP scaffold and the carotenoid structure could be uncoupled. Hence, with aim to split a difference between the structural recovery (informing on the protein structure) and the spectral recovery (informing on the electronic structure and environment of the carotenoid pigment), we monitored the kinetics of thermal OCP^R^ to OCP^O^ recovery using SAXS (OCP_wt-Ctag_) and optical spectroscopy at 467 nm (OCP_R27L-Ntag_, OCP_wt-Ntag_ and OCP_wt-Ctag_). For the structural monitoring, accumulation of the OCP^R^ state was achieved by illuminating a 23 mg/ml (620 µM) solution of OCP_wt-Ctag_ for ∼ 30 minutes at 430 nm with a 500 mW continuous LED source collimated to ∼ 1 cm (*i.e.* the same illumination protocol used in SAXS experiments described in the previous section), providing the starting point of the kinetics, and scattering data were thereafter recorded every ∼ 12 s. It was established in our static SAXS measurements that formation of OCP^R^_wt-Ctag_ state is characterized by the presence, in the ΔI(*q*) difference profile, of a negative peak at ∼ 0.05 Å^-1^ and a positive peak at ∼ 0.12 Å^-1^ (Fig. 3*D*). Hence, we used as a proxy to evaluate the structural recovery of the starting OCP^O^_wt-Ctag_ state the ΔI = I(q)_t_ – I(q)_t0_ difference signal in the 0.05 - 0.5 Å^−1^ range, where I(q)_t_ and I(q)_t0_ are the scattering profiles at a generic time (t) after switching off the 430 nm light and before the 430 nm irradiation procedure, respectively (Fig. S5). These data show that when OCP^R^_wt-Ctag_ is accumulated at high concentrations by prolonged illumination, structural recovery of the dimeric OCP^O^_wt-Ctag_ state occurs on a time scale of tens of minutes.

The spectroscopic recovery of OCP_R27L-Ntag_, OCP_wt-Ntag_ and OCP_wt-Ctag_ was monitored in the 0.1 - 16 mg/ml concentration range, after accumulation of OCP^R^ by the same illumination protocol used in the static SAXS measurements. Just after switching-off the 430 nm light at time *t_0_* (*i.e.* when the concentration of OCP^R^ is at its maximum), the time evolution of the normalized absorption difference [A(*t*) - A(*t_0_*)]/[A(*t_max_*) - A(*t_0_*)] at 467 nm was monitored, where A(*t*) and A(*t_max_*) are the absorption values measured at a generic time *t* during the thermal recovery and at the time *t_max_* = 5000 s, where the initial OCP^O^ state has been 95-99 % recovered (Fig. S5), respectively. The 467 nm wavelength was chosen because it is that at which the difference-absorption signal is maximal (highest contrast) between OCP^O^ and OCP^R^. Normalization was performed to enable direct comparison of data (Fig. S7) and of the fits at the various tested concentrations, viz. 0.1, 1.7, 3 and 16 mg/ml (Fig. S8). Alike the structural recovery, spectroscopic recovery occurs on a time scale of tens of minutes, with recovery slowing down as the protein concentration increases. Regardless of the concentration, however, OCP_R27L-Ntag_ recovers faster than OCP_wt-Ntag_, which itself recovers faster than OCP_wt-Ctag_. As OCP_R27L-Ntag_ and OCP_wt-Ntag_ differ only in their capacity to form OCP^O^ dimers, and OCP_wt-Ntag_ and OCP_wt-Ctag_ only in the location of the his-tag, these observations suggest that (i) the re-association of OCP^O^ monomers into dimers is the limiting step in the recovery of OCP_wt-Ntag_; and (ii) presence of a his-tag at the C-terminus negatively impacts dark-state recovery. Based on the results described in the previous sections we can expect that the thermal recovery of OCP_wt-Ctag_ involves at least the following states: OCP^R^ higher order oligomer → OCP^R^ dimer → OCP^R^ monomer → OCP^O^ monomer → OCP^O^ dimer; while that of OCP_R27L-Ntag_ transitions across OCP^R^ higher-order oligomers → OCP^R^ dimer → OCP^R^ monomer → OCP^O^ monomer. We attempted fitting the recovery data using constrained (same set of lifetimes for the four concentrations) mono-to-quadruple exponential functions (see Materials and Methods), where (i) a**_ci-_**_1_, a**_ci-_**_2_, a**_ci-_**_3_ and a**_ci-_**_4_ are pre-exponential factors depending on the protein variant and the concentration, (ii) τ**_ci-_**_1_, τ**_ci-_**_2_, τ**_ci-_**_3_ and τ**_ci-_**_4_ are lifetimes describing up to four separate molecular steps. A minimum of three exponentials are needed to account for the recovery kinetics of the three OCP variants. The lifetimes derived from the OCP_R27L-Ntag_ data (τ**_ci-_**_1_= 59.8 ± 0.3 s; τ**_ci-_**_2_ = 159.1 ± 0.6 s; τ**_ci-_**_3_ = 641.7 ± 1.1 s, Table S2) are nearly half of those derived from the OCP_wt-Ntag_ data (τ**_ci-_**_1_= 94.5 ± 0.2 s; τ**_ci-_**_2_ = 356.5 ± 1.2 s; τ**_ci-_**_3_ = 1327.5 ± 4.3 s, Table S6), and no better fits are obtained by accounting for a fourth component (similar chi^2^ for either dataset; Tables S6 and S7), precluding extraction of a lifetime for dimerization of OCP_wt-Ntag_. A similar three-exponential fitting procedure applied to the OCP_wt-Ctag_ (Table S3) data yields longer lifetimes (τ**_ci-_** _1_= 76.7 ± 0.2 s; τ**_ci-_**_2_ = 593.1 ± 2.1 s; τ**_ci-_**_3_ = 9517.4 ± 75 s), in line with the observed slower recovery, yet the data are better fitted (∼4 times smaller chi^2^, *i.e.,* a value close to those obtained for OCP_R27L-Ntag_ and OCP_wt-Ntag_ regardless of whether three or four exponential components are considered) by a four-exponential model (τ**_ci-_**_1_= 51.5 ± 0.2 s; τ**_ci-_**_2_ = 209.9 ± 1.3 s; τ**_ci-_**_3_ = 946.0 ± 3.7 s; τ**_ci-_**_4_ = 14057.7 ± 68.8 s; Table S6). A global four exponential-fit of the three data sets, whereby the lifetimes for OCP_R27L-Ntag,_ OCP_wt-Ctag and_ OCP_wt-Ntag_ were constrained to be identical and the simpler kinetics of OCP_R27L-Ntag_ (no dimerization of OCP^O^) accounted for by constraining to zero the population parameter a**_ci-_**_4_, corresponding to the longest lifetime, was unsuccessful. Our data thus point to at least three, and in the case of OCP_wt-Ctag_ four independent molecular steps being involved in the OCP^R^ to OCP^O^ recovery.

### Kinetics of photoactivation and recovery upon formation of OCP^R^ by pulsed illumination

To investigate short-live intermediate states which exist along the OCP^O^ to OCP^R^ and OCP^R^ to OCP^O^ transitions, we used ns-s timescale transient absorption spectroscopy. Briefly, 8 ns pulses from a 470 nm laser (5 mJ pulse energy) were employed to trigger photoactivation, and difference absorbance (ΔA) signals at 440 and 565 nm were monitored, respectively. The 440 nm probe wavelength is located at the blue edge of OCP^O^ absorption spectrum, and therefore informs on the depletion and recovery of this state upon and after excitation by the 470 nm pulse, respectively. Conversely, the 565 nm probe is at the red edge of the OCP^R^ absorption spectrum and thus serves as an indicator for the formation of the red absorbing states, including the first probed photoproduct P_2_ (λ_ΔAmax_ = 565 nm, with occupancy proportional to the ΔA signal), the photoactive OCP^R^ (λ_ΔAmax_ = 550 nm) and all red intermediate states between these (27).

We first investigated OCP_wt-Ctag_, OCP_R27L-Ntag_ and OCP_wt-Ntag_ at 12 µM concentration, *i.e.* ∼ 0.18 mg/ml (Fig. 5*A*), where nearly as many OCP^O^_wt-Ctag_ and OCP^O^_wt-Ntag_ are present in the monomeric form (54%, *i.e.* 6.5 µM) and in the dimeric form (46 %, *i.e.* 2.79 µM (Table S4)), assuming a similar dissociation constant of 14 µM for the two wild-type variants. Given this starting concentration, the pulsed nature of the excitation and the low quantum yield of photoactivation, OCP^R^ cannot accumulate in these experiments, hence the formation of OCP^R^ dimers and higher-order oligomers is extremely unlikely. We examined the temperature dependence (8°-36° C) of the ΔA signals at 440 nm and 565 nm in OCP_wt-Ctag_, with the aim to identify key intermediate states and determine their associated enthalpy and entropy of activation. Experiments revealed that irrespective of temperature, all kinetics recorded at 440 nm start from the same ΔA value, viz. −2 mOD (Fig. S9), indicating that depletion of the initial OCP^O^_wt-Ctag_ state (with ∼ 0.7 % yield, as determined by Ruthenium actinometer; see Materials and Methods) hardly depends on the temperature. Notwithstanding, the initial (∼ 50 ns) ΔA signal at 565 nm decreases with increasing temperature, amounting to 3 mOD at 8° C but only 1.5 mOD at 36° C (Fig. S9*A* and S9*E*).

**Figure 5.**
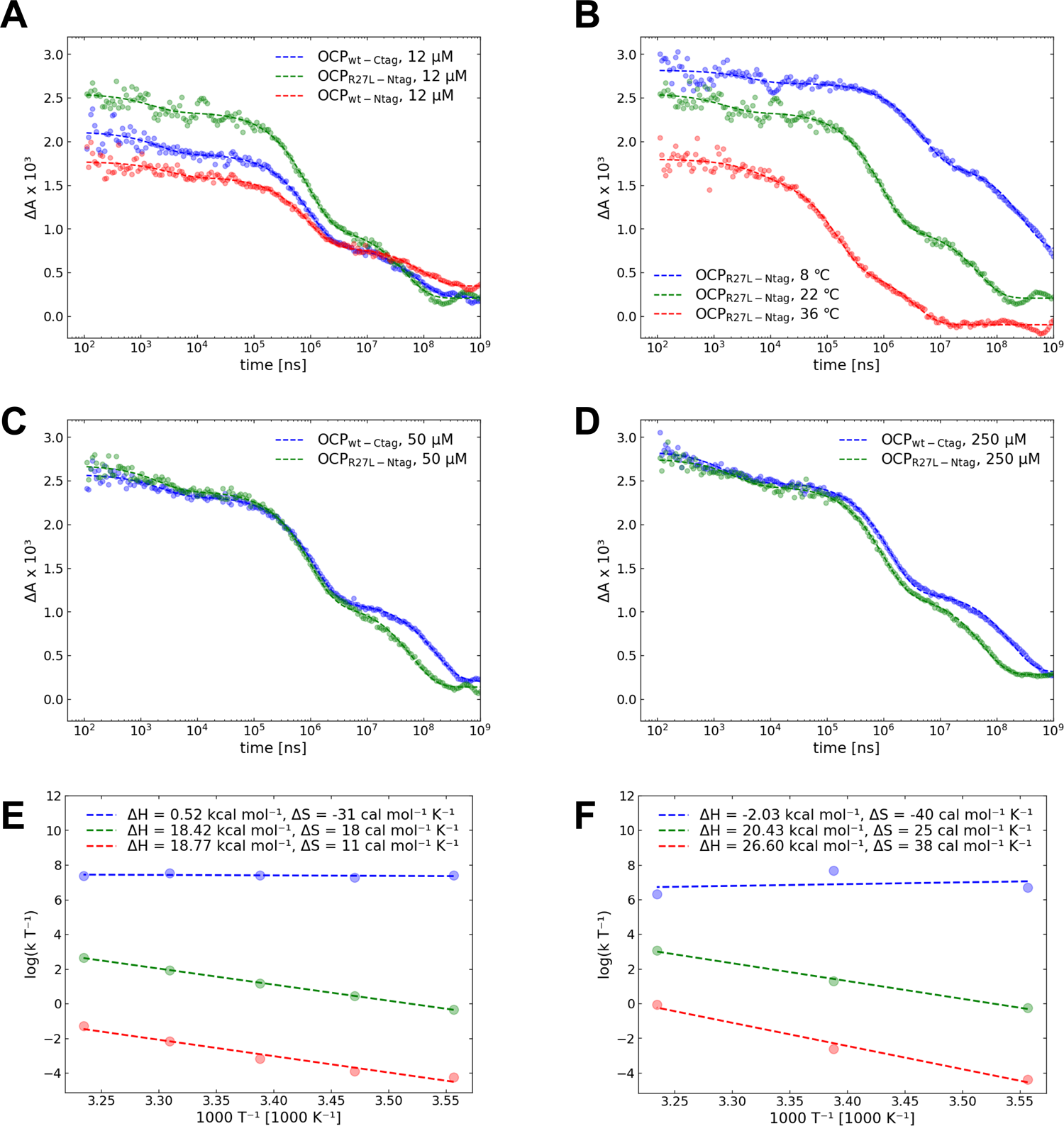
Transient absorption spectroscopy enables identification of three intermediate states along the formation of OCP^R^ and OCP^R^ to OCP^O^ thermal recovery. Difference absorbtion (ΔA) signals were monitored at 565 nm. (*A*) Nanosecond transient absorption data for OCP_wt-Ctag_, OCP_wt-Ntag_ and OCP_R27L-Ntag_ at 12 µM. (*B*) Nanosecond transient absorption data for OCP _R27L-Ntag_ at 12 µM and various temperatures. (*C* and *D*). Comparison of nanosecond transient absorption data between OCP_wt-Ctag_ and OCP_R27L-Ntag_ at 50 µM, and OCP_wt-Ctag_ and OCP_R27L-Ntag_ at 250 µM, respectively. (*E*-*F*) Eyring plots for OCP_wt-Ctag_ (*E*) and OCP_R27L-Ntag_ (*F*). These enable extraction of enthalpy and entropy of activation (κ = 1 is assumed). See Figure S9 for complementary data.

The ΔA signal for OCP_wt-Ctag_ at 565 nm appears triphasic in the 8-36 °C range (Fig. S9*A* – S9*F*), and likewise for the 440 nm ΔA signal in the 8-22 °C range (Fig. S9*C* – S9*F*). Similar lifetimes and activation energies can be derived from the two sets of data (Table S4 and Fig. 5*E*, respectively), confirming that the two sets of difference absorbance signals at 440 and 565 nm probe the same molecular events. Hence, we opted for a global tri-exponential fitting of the 440 and 565 nm data. Note that an additional fourth component intrudes into kinetics after 100 ms at 29° C and 36° C (Fig. S9*D* and S9*E*), which presumably lives longer than our time window at lower temperatures. Lifetimes were extracted for each step (Table S8), and Eyring plots generated from the measurements carried out at different temperatures (Fig. 5*E*). The fastest component (∼ 2 µs) displays an adiabatic behavior and is associated with a negative change in entropy. The two other components, characterized by lifetimes in the 0.2 - 5 and 10 - 250 ms range, respectively, are both characterized by increase of the entropy (11-18 cal.mol^-1^.K^-1^), indicating irreversibility, and display similar enthalpies of activation, viz. 18 kcal/mol (Fig. 5*E*).

The extent of recovery of the ΔA signal at 440 nm is slightly higher at increased temperature, with ≈ 50 % of the initially-depleted OCP^O^ state (0.7 % at 50 ns) having recovered after 1 s at 8 °C, but up to 75 % at 36° C (Fig. S9). Briefly, overall OCP^R^ yields of 0.35, 0.26, 0.25, 0.20 and 0.18 % are found at 8, 15, 22, 29 and 36 °C based on the 440 nm data at 500 ms, respectively. Hence, the recovery of the OCP^O^ state somehow benefits from increased thermal energy, but is not completed after 1 s (*i.e.* the ΔA signal at 440 nm is not null), regardless of temperature (Fig. S9). Hence, the limiting step in the monomeric OCP^R^_wt-Ctag_ to OCP^O^_wt-Ctag_ transition occurs on a time scale longer than 1s (observed in SAXS data, Fig. S3 and Table S2 and S3). The 565 nm kinetics also decay faster at higher temperatures, confirming the drop in yield for the red states (including OCP^R^_wt-Ctag_) as the temperature augments. Surprisingly, however, the ΔA signal at 565 nm is null after 1 s at 29 °C (and negative at 36 °C), which would suggest that no OCP^R^_wt-Ctag_ remains (yields of 0.8, 0.44, 0.18, 0.02 and −0.16 % based on the 565 nm data at 8, 15, 22, 29 and 36 °C, respectively). These diverging results illustrate the usefulness of probing photoactivation and recovery at the two extremes of OCP^O^ and OCP^R^ absorption spectra to obtain meaningful insights into the complex underlying mechanisms.

We repeated the above-described experiments on the constitutively monomeric OCP_R27L-Ntag_ mutant. The 565 nm kinetics, shown in Fig. 5, are overall similar for OCP_wt-Ctag_ and the OCP_R27L-Ntag_ mutant. From the multiexponential fits, we accordingly derive comparable lifetimes, with the main difference between the two proteins being a two-times shorter τ_3_ component for OCP_R27L-Ntag_, as compared to OCP_wt-Ctag_ (Table S8). Thus, only the last component shows a putative influence of the presence of dimers (recall that 46 % of OCP^O^_wt-Ctag_ is present in the form of dimers at 12 µM). Accordingly, the Eyring plots of OCP_R27L-Ntag_ (Fig. 5*F*) reveal analogous trends to those of OCP_wt-Ctag_ for the two first components, while the τ_3_ component is characterized by larger activation enthalpy and entropy. To determine whether or not these differences could arise from the position of the his-tag at the N- or C-terminus of the protein, we further investigated OCP_wt-Ntag_ at 22° C, and compared lifetimes with those characterizing OCP_R27L-Ntag_ and OCP_wt-Ctag_ (Table S8). We again observed similar dynamics, with derived lifetimes and relative contributions for the three steps being almost identical in OCP_wt-Ntag_ and OCP_wt-Ctag_. An interesting difference is, however, that despite a similar P_2_ yield, OCP_wt-Ntag_ displays a nearly two-times increased OCP^R^ yield at 500 ms compared to OCP_wt-Ctag_. This observation suggests that dissociation of the NTE from the CTD, facilitated by adjunction of an his-tag at the N-terminus, is a limiting step in the photoactivation of OCP. This event is yet silent in terms of changes in the absorption of the carotenoid, since none of the components identifiable in the 440 and 565 nm kinetics are affected by the change in tag position in OCP_wt-Ntag_ and OCP_wt-Ctag_.

Transient spectroscopy data presented so far were acquired in conditions where OCP^O^_wt-Ctag_ and OCP^O^_wt-Ntag_ monomers are slightly prevalent (54 %), and differences with the constitutively monomeric OCP^O^_R27L-Ntag_ were seen only for the τ_3_ component, suggesting that the underlying step us sensitive to the dimerization of the protein. To verify this hypothesis, we repeated experiments on OCP_wt-Ctag_ and OCP_R27L-Ntag_ at 50 and 250 µM, where dimers account for 78 and 95 % of the OCP^O^ population, respectively (Fig. 5*C* and 5*D*). Only the ΔA signal at 565 nm was probed. Both the initial P_2_ yield at 50 ns and the overall OCP^R^ yield at 0.1 s increase with concentration for OCP_wt-Ctag_, but not for OCP_R27L-Ntag_, indicating that stabilization of the closed conformation of OCP^O^ through dimerization (54) favors the initial steps of photoactivation (Fig. S10*A* and Table S8). The lifetimes derived for the OCP_R27L-Ntag_ sample are only marginally affected by the increase in concentration, whereas a nearly two-fold increase in τ_3_ lifetime is seen for OCP_wt-Ctag_ at 50 and 250 µM. These experiments confirm that the τ_3_ component, although present also in the monomer, is affected by dimerization. Specifically, a plot of τ_3_ lifetime as a function of OCP dimers concentration, assuming for zero dimer concentration the average τ_3_ lifetime of OCP_R27L-Ntag_, reveals a logarithmic behavior with half-maximum reached at ∼ 10 µM dimer, corresponding to ∼ 25 µM OCP^O^_wt-Ctag_ (Fig. S10*B*). It is interesting to note that regardless of the sample and concentration, the τ_2_ component (∼ 1 ms) has an almost constant value and relative contribution in all kinetics measured at 22 °C. The underlying step is therefore neither affected by tagging nor monomerization.

### Time-resolved X-ray scattering

Time-resolved X-ray scattering measurements were carried out using a pump-probe scheme with view to investigate the large-scale structural changes occurring in OCP_wt_ upon photo-activation, and the associated time scales. We used two variants of OCP_wt,_ viz. OCP_wt-Ntag_ and OCP_wt-Ctag_, enabling to test whether or not presence of a histidine-tag at the N- or C-terminus influences pulsed photoactivation kinetics. Indeed, it has been observed that both OCP^R^ accumulation under prolonged illumination and the subsequent thermal recovery of the OCP^R^ state are faster for OCP_wt-Ntag_ than OCP_wt-Ctag_ (14, 54). From our ns-s transient absorption characterization of photoactivation and recovery in OCP_wt-Ntag_ and OCP_wt-Ctag_ and the derived Eyring plots, it was expected that at 22° C, the two steps involving large scale motions are characterized by lifetimes of ≈ 1 and ≈ 80 ms (Fig. 5). Hence, time delays (Δt) between the pump optical-laser pulse and the probe X-ray pulse ranged from 15 µs to 200 ms. Experiments were performed at 23 mg/ml (620 µM) OCP_wt-Ntag_ and OCP_wt-Ctag_, *i.e.* a concentration at which dimers account for ∼ 98 % of the protein.

The experimental setup designed for this experiment, described in the Experimental Section and illustrated in Fig.6*A*, involves the circulation of a large volume of sample across the X-ray beam to minimize cumulative X-ray damage during the experiment. The circulation speed was set so as to ensure that the sample is kept in the pump–probe intersection area during a pump–probe sequence, but refreshed between two consecutive pump pulses. We verified that with this setup, the same difference scattering signal observed in static SAXS experiments could be obtained upon continuous illumination of the sample reservoir by a 500 mW LED source emitting at 430 nm. The LED illumination was prolonged until the photoinduced change in scattering signal was stable. The resulting difference scattering profile is similar to that produced from our static measurements, *i.e.,* it features a negative peak at ∼0.05 Å^-1^ and a positive peak at ∼0.12 Å^-1^ (Fig. 3*D*). This control indicates that the use of a different continuous illumination setup at a different beamline does not result in shifts in the difference scattering signal (Fig. 6*B*).

**Figure 6.**
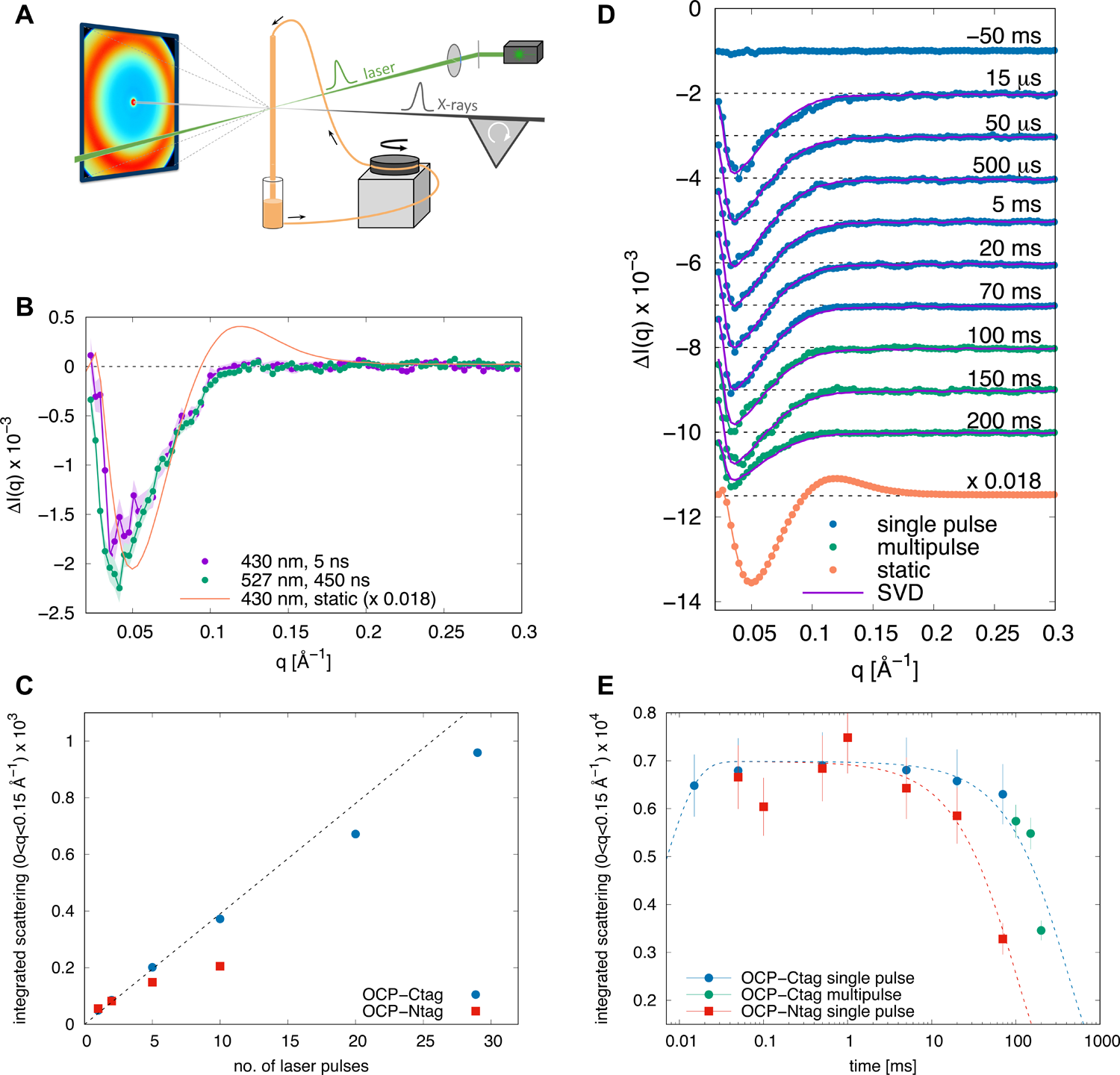
Time resolved X-ray scattering reveals the kinetics of light-induced structural changes in the µs-ms time scale. (*A*) Schematic representation of the experimental setup used for time-resolved SAXS/WAXS experiments at the ID09 beamline of ESRF. The OCP solution (orange) flows through a quartz capillary connected to a peristaltic pump. A nanosecond laser pulse (green) synchronized with a microsecond X-ray pulse train (gray) is used to trigger OCP^O^ to OCP^R^ photoconversion. Protein structural changes give rise to changes in the the X-ray scattering pattern measured on a CCD detector. (*B*) Comparison of the time resolved laser-on - laser-off X-ray difference profile at Δt = 1 ms after excitation by both a 5 ns pulse at 430 nm and a 450 ns pulse at 527 nm with the static OCP^R^ - OCP^O^ X-ray difference profile (scaled by a factor 0.018) calculated by subtracting the profile of the dark-adapted sample from that illuminated at 430 nm. (*C*) Integrated difference scattering in the region 0 < q < 0.15 Å⁻¹ as a function of the number of laser pulses (527 nm) for OCP_wt-Ctag_ (blue circles) and OCP_wt-Ntag_ (red squares). Pulse energy and duration were 2 mJ and 400 ns, respectively, and the pulses were spaced by 1 ms. (*D*) Light-induced time-resolved X-ray scattering difference profiles for OCP_wt-Ctag_. The static OCP^R^ – OCP^O^ X-ray scattering difference pattern (scaled by a factor 0.018) is also shown for comparison. Data are vertically offset for clarity. (*E*) Time evolution of the integrated difference scattering in the region 0 < q < 0.15 Å⁻¹ after laser excitation at 527 nm for OCP_wt-Ctag_ (blue and green circles) and OCP_wt-Ntag_ (red squares). Dashed lines are fitting curves with a double exponential.

With intention to determine the extent to which photoactivation can be increased by successive re-excitations within the pulse, we first measured the ‘light’ (laser-on) minus ‘dark’ (laser-off) difference scattering signal at Δt = 1 ms, using two different laser pulse durations, viz. 5 and 450 ns. This required the use of two different lasers, emitting at 430 and 527 nm, respectively. At these wavelengths, the extinction coefficient of OCP^O^ is similar (≈ 35 000 M^-1^ cm^-1^), suggesting that the OCP^R^ formation quantum yield should be comparable and that an increase in the difference signal at 527 nm would thus stem from successful re-excitation within the 450 ns pulse. Both pulse durations are indeed longer than the time necessary to recover the electronic ground state (∼ 20 ps) (27, 55). A pulse power density of 3 mJ.mm-2 was chosen, affording the delivery of ∼ 75 photons per chromophore across the pulse, corresponding to a photon every 0.06 and 5 ns for the 5 and 450 ns pulse durations, respectively. The ‘light’ (laser-on) minus ‘dark’ (laser-off) difference scattering profiles obtained 1 ms post-excitation by either laser feature a negative peak in the small angle region up to ≈ 0.1 Å^−1^, but the amplitude of the peak does not vary significantly, indicating that stretching the pulse up to 450 ns does not improve the photoactivation extent (Fig. 6*B*). Thus, OCP molecules that have not engaged towards formation of a red state (P_1_) and are back to the ground state (27) cannot be re-excited within the 450 ns pulse. Given the fact that these molecules do not show a difference signal in transient absorption spectroscopy experiments – meaning that the carotenoid is back in the orange state and therefore presumably H-bonded to the protein scaffold – their inability to re-enter a photocycle could be rooted in the fact that some protein residues, which are important for photoactivation but do not influence the spectrum, have not returned yet to their initial conformation after 450 ns. We refer to this *hitherto* undescribed intermediate state, which accounts for 99.3 % of the OCP^O^ population upon pulsed illumination at 22° C, as the OCP “numbed-state”, abbreviated OCP^NS^. To estimate an upper boundary for the lifetime of this intermediate, the repetition rate (1 kHz) of the 527 nm laser, coupled with a synchronized chopper/shutter system, was used to measure time-resolved profiles at a time delay of 100 ms post-excitation by an increasing number (up to 30 pulses) of consecutive pulses spaced by 1 ms. The scattering difference signal (laser-on minus laser-off) was then integrated in the 0-0.15 Å^-1^ range and plotted as a function of the number of pulses (Fig. 6*C*). The signal increases linearly with the number of pulses up to 5 pulses for OCP_wt-Ntag_ and up to 10 pulses for OCP_wt-Ctag_, after which a plateau is observed, presumably due to concomitant recovery of the dark state structure (see below). Thus, the lifetime of the non-re-excitable OCP^NS^ can be estimated to lie between 450 ns and 1 ms. Irrespectively, comparison of the two time-resolved difference signals with the static difference scattering signal shows that the shape of the difference profile generated by pulsed excitation does not coincide with that obtained by stationary illumination. In particular, the position of the negative peak differs, and the positive peak at ∼ 0.12 Å^-1^ is absent (Fig. 6*C*).

A complete time-resolved dataset was measured at time delays between 15 µs and 200 ms. Data for OCP_wt-Ctag_ are shown in Fig. 6*D*, while analogous results are obtained for OCP_wt-Ntag_. Fig. 6*D* demonstrates that substantial changes in the scattering profiles are (i) already visible after 15 µs, (ii) maximized before 1 ms, and (iii) substantially damped by 200 ms. To further investigate the kinetics of the scattering signal changes, the time-resolved scattering dataset was analyzed by the singular value decomposition (SVD) method (see Fig.S12 and Materials and Methods), which revealed that the entire dataset can be described by a single time-independent basis pattern (continuous line in Fig. 6*D*). In other words, no other intermediate state than that responsible for this basis pattern is visible in our data. It must be noted that the difference between the time-resolved and static difference signals (*i.e.* the shift in q-value of the negative peak and the absence of a positive peak) is observed for all recorded time delays (15 µs-200 ms). Notwithstanding, a rough estimation of the photoactivation extent by pulsed ns-excitation can be obtained by scaling the amplitude of the time-resolved difference signal to that measured in static experiments, pointing to a photoactivation efficiency of ∼ 1.8 % (Fig. 6*B*). The difference profiles shown in Fig. 6*D* were integrated in the 0 - 0.15 Å^-1^ range and the results are shown as a function of time for both OCP_wt-Ntag_ and OCP_wt-Ctag_ (Fig. 6*E*). The integrated intensity starts decaying between 1 and 10 ms in OCP_wt-Ntag_, and between 10 and 100 ms in OCP_wt-Ctag_, indicating a recovery of the dark state in these timescales, in full agreement with the results in Fig. 6*C*. Thus, large scale conformational changes start occurring as of ∼ 10 µs, maximize at 1 ms, and revert with apparent lifetimes (*i.e.* time at half-maximum amplitudes) of 69±14 and 280±50 ms for OCP_wt-Ntag_ and OCP_wt-Ctag_, respectively.

It is tantalizing, but difficult, to propose an interpretation for our observations. It has been proposed earlier that the first large-scale conformational change (∼ 1 ms time scale) occurring in OCP post-excitation is the detachment of the NTE from the CTD β-sheet, either concomitant or prior to a repositioning of the CTT, and that thereafter only may befall dissociation of the two domains (27, 29). Given the quantum yield of OCP and the fact that our TR-SAXS experiment was carried out at 23 mg/ml (620 µM), where dimers account for ∼ 98 % of the sample, it is yet extremely unlikely that two monomers in the same dimer would have been photoexcited by the same pulse (probability is the square of the quantum yield, *i.e.* 0.4 x10^-5^) and that our signal would inform on the formation of the photoactive OCP^R^. Furthermore, since the NTE is involved in the dimerization interface, the domain dissociation characteristic of the OCP^R^ state would have to await monomerization. Hence, it is most probable that the conformational change probed in our experiment is neither dimer or domain dissociation, but rather a subtle conformational change occurring in only one of the two monomers in a dimer.

## Discussion

In the present work, we used TR visible spectroscopy, as well as static and TR X-ray scattering to investigate the structural changes – including changes in quaternary structure – accompanying photoactivation of the orange carotenoid protein (OCP). It was repeatedly observed that OCP crystallizes as a dimer, yet it had remained elusive whether or not this interface is functional given the high concentration of the protein in crystals, viz. ≈ 23 mM. Mutation of a single highly-conserved residue at the crystalline dimerization interface results in the loss of the ability of OCP^O^ to dimerize, suggesting that the dark-adapted dimer could be functional (10). Collection and analysis of static X-ray scattering profiles of dark-adapted OCP_wt-Ctag_ as a function of concentration confirm the *in vitro* existence, already suggested in the literature (22, 23, 25, 26, 30, 56), of a dimer which becomes predominant at concentrations ≥ 1 mg/ml (Fig. 3*C* and Fig. S1). Our SAXS titration data (Kd ∼ 14 µM) is in accordance with (i) previous results from native mass spectrometry experiments, reporting the presence of OCP^O^ dimers at concentration as low as 3 µM (22); (ii) the recent assertion based on SAXS and SANS data that OCP^O^ resides in the same state at 1 and 65 mg/ml (*i.e.* 28 µM and 1.8 mM) (23, 24); and (iii) the dissociation constant that can be derived from size exclusion chromatography results presented in Fig. 3 of Muzzopappa et al. (10), *i.e.*, ≈ 0.6 mg/ml (17 µM), provided that we use the same extinction coefficient at 490 nm to estimate protein concentrations.

Analogous concentration-dependent analyses of static X-ray scattering profiles collected after accumulation of OCP^R^_wt-Ctag_ and OCP^R^_R27L-Ntag_ by 30 min irradiation at 430 nm suggest that they are also prone to oligomerization. Indeed, for both proteins, not only is the scattering signal different between the dark-adapted (OCP^O^) and light-adapted (OCP^R^) states, but that difference evolves as a function of concentration. Remarkably, OCP^R^_R27L-Ntag_ and OCP^R^_wt-Ctag_ display similar P(r) (Fig. 2*F*) and Rg (Fig. 3*B*) at the various tested concentrations, indicating that they form the same OCP^R^ state(s). The high-concentration (10.5 mg/ml) difference-scattering profiles between the light- and dark-adapted states, *i.e.* ΔI(q)_R27L-Ntag_ and ΔI(q)_wt-Ctag_, are characterized by the presence of a negative peak (at 0.07 and 0.05 Å^-1^, respectively) and a positive peak (at 0.14 and 0.12 Å^-1^, respectively) (Fig. 3*D*). These peaks respectively signal the loss and gain of interatomic distances in the OCP samples, upon illumination-triggered transition from the OCP^O^ to the OCP^R^ state. Given that OCP^R^_R27L-Ntag_ and OCP^R^_wt-Ctag_ form the same OCP^R^ species, the observation that the negative and positive peaks are found at different q-values in the ΔI(q)_R27L-Ntag_ and ΔI(q)_wt-Ctag_ difference profiles can be rationalized by the fact that their dark-adapted states differ – OCP^O^_R27L-Ntag_ being present as a monomer, whereas OCP^O^_wt-Ctag_ features a dimer.

The molecular envelopes reconstructed from the OCP^R^ data suggest that OCP^R^ dimers form as of 0.7 mg/ml (20 µM), while higher-order oligomers form at 10 mg/ml (∼300 µM) (Fig. 3*A*-*B*, Fig.S1, Fig.S2, Table S4 and Table S5). The observation that the R27L mutant also forms OCP^R^ dimers and higher-order oligomers suggests that the dimerization interfaces of OCP^O^ and OCP^R^ do not overlap. Based on this hypothesis, and on knowledge from the literature, we were able to build tentative models for OCP^R^ dimers and higher-order oligomers. The models notably assume that: i) OCP^R^ higher-order oligomers form from the coalescence of either two OCP^R^ dimers (yielding tetramers) or one OCP^R^ monomer and one OCP^R^ dimer (yielding trimers); ii) interactions between OCP^R^ monomers in a dimer and between OCP^R^ dimers and/or monomers in a higher-order oligomer involve the same interface in the two interacting monomers; iii) residues involved in assembly of OCP^R^ into dimers and higher-order oligomers are not exposed in the dark OCP^O^ dimer (51, 53); and iv) OCP^R^ dimers assemble via their CTD, so that their two NTD are exposed and free to interact with PBS via R155 (51), while the OCP^R^ higher-order oligomers form from an interaction between two NTDs using an interface similar to that observed in all three currently available crystal structures of the homologous HCP homologues (pdb ids: 5FCX and 6MCJ (40, 41)). This interface features the residue equivalent to R155 in OCP, and thus correspond to the face of the NTD that presides to the NTD/CTD interaction in OCP^O^. Our model for the OCP^R^_wt-Ctag_ dimer is in overall agreement with that proposed earlier (23), featuring two monomers associated by their CTD at an interface reminiscent of that observed at the A-type interface in the apo-*Anabaena* CTDH crystal structure. Of note, helices αN (or CTT) and αA (or NTE), which sit aside one another on this side of the β-sheet in the OCP^O^ state but detach upon photoactivation (9), were considered as disordered in OCP^R^ and therefore not included in our OCP^R^ dimer model. To account for the known exposure, in OCP^R^, of the face of the NTD that interacts with the CTD in OCP^O^, each NTD was rotated by 180° and further translated by a few Å to avoid clashes with the CTD. This face of the NTD features R155, which tethers the NTD to the CTD via H-bonding to E244 in OCP^O^ (Figure 1), but is essential for the interaction of OCP^R^ with the PBS. Our data do not allow to firmly assign the precise nature of higher-order oligomers. Indeed, the elongated molecular envelope derived from OCP^R^ samples at high concentrations (> 10 mg/ml) could either stem from the presence of elongated OCP^R^ trimers (formed by the interaction of an OCP^R^ monomers and an OCP^R^ dimer) or more compact OCP^R^ tetramers (formed by the interaction of two OCP^R^ dimers) or a mixture thereof. Supplementary Fig. S1 shows the superimposition of a tentative OCP^R^ tetramer model over the molecular envelopes derived from SAXS data collected on OCP^R^_wt-Ctag_ and OCP^R^_R27L-Ntag_ at ∼10 mg/ml. The molecular envelopes could yet also be explained by elongated OCP^R^ trimers or by a mixture of trimers and tetramers. An important question which remains open is that of the *in vivo* concentration of OCP, as it would determine whether or not the OCP^O^ and OCP^R^ dimers and OCP^R^ higher order oligomers are of functional relevance. The *in vivo* concentration of OCP remains elusive – notably the local concentration in the vicinity of the thylakoid membranes, where OCP localizes – hence we cannot ascertain that the concentrations used in our study are found in the cell. It must be recalled, however, that the overall mass-concentration of proteins in cells is estimated to lie between 200-300 mg/ml (57), corresponding to a molar concentration of ∼5 mM – *i.e.* nearly the concentration of proteins in crystals – assuming an average molecular weight 50 kDa. Hence, we may speculate that in cyanobacterial cells, the propensity of OCP to form dimers would be high even in the case where its overall concentration would be lower than the dissociation constant. Furthermore, OCP is localized in the vicinity of the thylakoid membranes, where its local concentration could be higher. Strongly supporting the relevance of OCP^R^ dimers is, in all cases, the recently determined cryo-electron microscopy structure of the OCP-quenched PBS (58) which reveals binding to the PBS of an OCP^R^ dimer. The overall similarity between our model and that presented in the preprint of the manuscript (coordinates are not accessible as yet) is notable. Given that the dissociation constants of OCP^O^ and OCP^R^ dimers are in the same order, it may be proposed that if the latter is functional, the second must be.

We used a combination of optical spectroscopy and X-ray scattering to investigate OCP^R^ to OCP^O^ thermal recovery and determine whether or not it is influenced by oligomerization processes. First, we monitored the structural recovery of the OCP^O^_wt-Ctag_ state, following accumulation of the OCP^R^_wt-Ctag_ state by prolonged illumination at 430 nm. The difference scattering profile features a positive and a negative peak (Fig. 3*D*), which together sign for the disappearance of the starting OCP^O^_wt-Ctag_ state and the concomitant appearance of the OCP^R^_wt-Ctag_ states. Hence, the time evolution of the difference signal in the 0.05 - 0.5 Å^-1^ q-range was used to follow the overall thermal recovery, which after prolonged illumination occurs on the time scale of tens of minutes (Fig. S4).

Further insights in the OCP^R^ to OCP^O^ thermal recovery were attained by resorting to spectroscopy at various protein concentrations. First, we accumulated OCP^R^ by prolonged illumination at 430 nm, and monitored recovery of OCP^O^ by following absorption at 467 nm. Regardless of the sample, we found that the recovery time markedly increases as a function of concentration, in full agreement with the indication from static X-ray scattering data that both OCP^O^ (OCP_R27L-Ntag_, OCP_wt-Ntag_ and OCP_wt-Ctag_) and OCP^R^ (OCP_R27L-Ntag_, OCP_wt-Ntag_ and OCP_wt-Ctag_) can oligomerize. Indeed, the more oligomeric OCP^R^ states present, the higher the overall energy barrier for a full recovery of the starting dark OCP^O^ to occur, and therefore, the longer it will take. Irrespective of the assayed concentration, the spectroscopic recovery of OCP^O^_wt-Ntag_ was found to be slower than that of OCP^O^_R27L-Ntag_, indicating that the reassembly into dimers of OCP^O^ monomers is the step that limits OCP^O^_wt-Ntag_ recovery. Likewise, the recovery of OCP^O^_wt-Ctag_ was slower than that of OCP^O^_wt-Ntag_, indicating that presence of the his-tag at the C-terminus adversely affect recovery of the dark-adapted state. Recovery data could only be fitted with three- or four-exponential functions, suggesting occurrence of multiple sequential steps. We plotted the pre-exponential factors as a function of concentration to follow the evolution of the associated populations (Fig. 4*A*-*C* and Fig. S8). Focusing first on results from the three-exponential fits, we observe a similar evolution as a function of concentration in the three samples: (i) the amplitude of the fastest component (a**_ci-_**_1_, τ**_ci-_**_1_) decreases; (ii) the amplitude of the intermediate component (a**_ci-_**_2_, τ**_ci-_**_2_) initially increases but then decreases; (iii) the amplitude of the slowest component (a**_ci-_**_3_, τ**_ci-_**_3_) steeply increases. The observation of intersections at 0.5 population between amplitudes of rising and declining components is indicative of a balance in their relative populations and therefore is suggestive of sequentiality between the underlying steps (Fig. 4*A-C*). The populations of the first and second components feature such an intersection at concentrations of ∼ 0.7 and 1.8, in OCP_R27L-Ntag_ (Fig. 4*A*) and OCP_wt-Ntag_ (Fig. 4*C*), and at 6 mg/ml, in OCP_wt-Ctag_ (Fig. S8*C*). Assuming that the first (a**_ci-_**_1_; τ**_ci-_**_1_) and second (a**_ci-_**_2_; τ**_ci-_**_2_) components inform on the contribution to recovery of OCP^R^ monomers and dimers, a possible explanation is that presence of the his-tag at the C-terminus challenges dimerization of OCP^R^ in OCP_wt-Ctag_, elevating the apparent dissociation constant. For OCP_R27L-Ntag_ and OCP_wt-Ntag_, we furthermore see that the population of third component intersects those of the first (at ∼ 1.7 and ∼ 7 mg/ml, respectively) and second components (at ∼ 9.5 and ∼ 14.5 mg/ml, respectively). Assuming that the slowest component informs on the contribution of OCP^R^ higher-order oligomers to the recovery signal, this observation would be suggestive of a facilitated accumulation of these in OCP^R^_R27L-Ntag_, compared to the OCP^R^_wt-Ntag_ – possibly due to elimination of a competition between the dark-adapted dimerization and light-adapted trimerization and/or tetramerization interfaces. Indeed, both are localized in the NTD of OCP_wt-Ntag_. In the OCP_wt-Ctag_ sample, we do not see intersections of the slowest component amplitude with those of others components, when results of the three-exponential fit are considered. However, a much better agreement is obtained when the OCP_wt-Ctag_ data are fit with four-exponentials (Tables S6 and S7), yielding a chi^2^ comparable to that obtained with three exponentials for OCP_R27L-Ntag_ and OCP_wt-Ntag_. Plotting the four amplitudes as a function of concentration, the same observations can be made for the OCP_wt-Ctag_ (Fig. 4*C*), *i.e.,* intersections at 0.5 population can be seen at ∼ 1.7 mg/ml for the first and second component, and at ∼ 10 mg/ml for the second and third component. The population of the additional fourth component, characterized by a lifetime in the order of ∼ 4 hours (14057.7 ± 68.8 s), varies from ∼ 0.04 to 0.1 with increasing concentrations, indicating that the contribution of this rate-limiting step to the recovery signal is small. We see two possible explanations for the presence of this additional component. A first hypothesis is that it informs on the dimerization of OCP^O^_wt-Ctag_ monomers, which due to absence of a tag at the N-terminus would reassemble faster than OCP^O^_wt-Ntag_, explaining that a lifetime can be extracted for the former, but not for the latter, in our experiments (t_max_ = 5000 s). Another hypothesis is that the additional component informs on a step that is dramatically slowed down in OCP_wt-Ctag_, compared to OCP_wt-Ntag_, e.g. the reattachment of the CTT to the F-side of the CTD β-sheet. Indeed, the CTT is longer and presumably more flexible in OCP_wt-Ctag_ compared to OCP_wt-Ntag_ and it was accordingly shown that OCP^O^_wt-Ctag_ recovers faster than OCP^O^_wt-Ntag_ (14, 54). Our data do not allow to favor one hypothesis over the other. However, the doubling of the lifetimes extracted for OCP_wt-Ntag_, compared to OCP_R27L-Ntag_, provides circumstantial evidence that dimerization of the dark-adapted OCP^O^_wt-Ntag_ is the limiting step in its recovery. Therefore, we favor the first hypothesis. We accordingly propose that our recovery data inform on four molecular steps (Fig. 7) involved in the concentration-dependent OCP^R^ to OCP^O^ thermal recovery: (i) the transition from monomeric OCP^R^ to OCP^O^, characterized by a lifetime τ_ci-1_ = 50-100 s; (ii) the dissociation of OCP^R^-dimer into OCP^R^-monomers, characterized by a lifetime τ_ci-2_ = 160-600 s and a dissociation constant in the order of 14-30 µM (recovery data point to values of 30-40 µM but the observation that OCP^R^_R27L-Ntag_ attains dominance at 0.7 mg/ml, where occupancy of the OCP^O^_wt-Ctag_ dimer is partial (0.7 mg/ml), suggests a dissociation constant below 14 µM); (iii) the dissociation of OCP^R^ higher-order oligomers into OCP^R^-dimers and/or monomers, characterized by a lifetime τ_ci-3_ = 640-950 s and a roughly 20-fold higher dissociation constant (∼ 280 µM), since these oligomers are visible as of 10 mg/ml; (iv) the re-association of OCP^O^-monomers into OCP^O^-dimers, characterized by a lifetime τ_ci-4_ > 14 057 s and a dissociation constant in the order of ≈ 0.5 mg/ml (∼ 14 µM). That OCP^O^ dimer formation is slowed down in OCP_wt-Ntag_, compared to OCP_wt-Ctag_, could result from presence of the his-tag at the N-terminus, *i.e.,* in the vicinity of the OCP^O^ dimerization interface.

**Figure 7.**
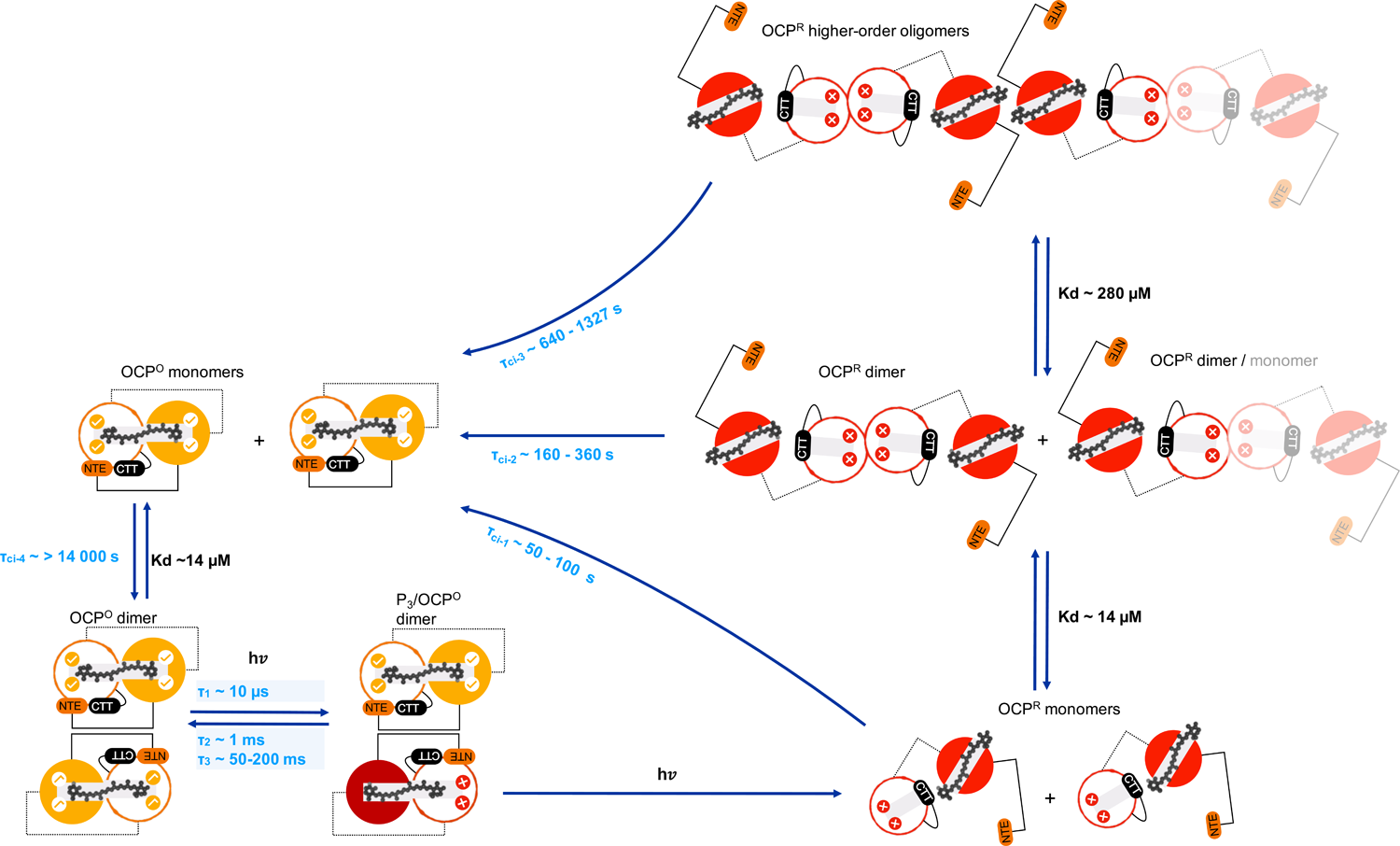
Proposed model for OCP photoactivation and recovery upon continuous illumination. OCP monomers and dimers are represented schematically with their N-terminal (coloured in orange and red for OCP^O^ and OCP^R^, respectively), and C-terminal domains (coloured in white). The ground state keto-carotenoid is shown in grey. Ticks and crosses in the CTD indicate presence and absence of H-bonds between the carotenoid and CTD residues Tyr201 and Trp288. The presence of ticks in the NTD indicate presence of π-stacking interactions between the carotenoid and NTD residues Tyr44 and Trp110. Our data show that dark-adapted OCP^O^_wt_ forms dimers, with a dissociation constant of ∼ 14 µM. The dimerization interface at play is likely that which has been repeatedly observed in OCP crystal structures. In the event where one of the two monomers in a dimer is activated, it may undergo all photoactivation steps until formation of P_3_, but will promptly revert to the dark state unless the second monomer is as well photoactivated, forming a P_3_/OCP^O^ dimer). Indeed, domain dissociation can only occur after dimer disassembly, which itself can only befall after NTE detachment. The latter is in turn dependent on the carotenoid achieving a successful transit across the NTD as required to induce conformational disorder in αC, αB and consequently αA. In practice, this means that reaching the OCP^R^ state in a pulsed illumination experiment probing OCP_wt_ dimer is extremely unlikely (probability = QY^2^ = 0.4 x10^-5^), unless re-excitation can be achieved within the lapse of the excitation pulse. Under continuous illumination, however, the OCP^R^ state can be accumulated, which first coalesces into dimers, with a dissociation constant similar to that displayed by dark-adapted OCP (∼ 14 µM), but may further progresses to the formation of higher-order oligomers at higher concentrations (the dissociation constant is at least 20 times higher, *i.e.,* ∼ 286 µM). Higher stabilization of OCP^R^ in these oligomers elevates the overall energy barrier for a full recovery of the starting dark OCP^O^ to occur, and therefore delay the upturn of the OCP^O^ state. At concentrations where the dark OCP^O^ dimer may form, it is yet the reassociation of dark monomers into dimers (τ_ci-4_) of ∼ 14 000 s, that constitute the limiting step in the recovery. Hence, OCP photoactivation and recovery are limited by oligomerization processes.

To obtain additional insights into molecular events occurring on shorter time scales, we used pulsed ns illumination to generate OCP^R^ (with a yield of ≈ 0.7 %), and monitored the ns-s timescale evolution of the difference absorption signal (ΔA) at the blue and red edges of the OCP^O^ (440 nm) and OCP^R^ (565 nm) absorption spectra, respectively. The use of the two wavelengths was justified by our need to obtain insights into both the recovery of the OCP^O^ state (ΔA440 nm) and the formation and disappearance of OCP^R^ and preceding red intermediates (ΔA565 nm). The wild type OCP_wt-Ntag_ and OCP_wt-Ctag_ were investigated, as well as the constitutively monomeric OCP_R27L-Ntag_ mutant. Regardless of the OCP sample, we could identify three intermediate states with lifetimes of ∼ 2 µs, ∼ 1 ms and ∼ 40-200 ms, respectively, at 22° C. The lifetimes derived from OCP_wt-Ntag_ and OCP_wt-Ctag_ data are nearly identical, indicating that the presence of the his-tag does not influence the dynamics probed at the carotenoid level on the ns-s time scale. Nonetheless, a higher OCP^R^ yield was found for OCP_wt-Ntag_, demonstrating influence of the tag on a step that is not rate-limiting, and whose contribution to photoactivation and recovery is silent from the spectroscopic perspective. In these experiments, carried out at a concentration where the dimeric form accounts for 46% of the wild-type proteins (12 µM), the first and second components were found to not vary as a function of the protein variant, while a two-times reduced lifetime was found for the third component in the constitutively monomeric OCP_R27L-Ntag_. Additional experiments carried out at 50 and 250 µM concentrations of OCP_R27L-Ntag_ and OCP_wt-Ctag_, where the prevalence of OCP_wt-Ctag_ dimers are 78 and 95 %, respectively, revealed a marginal influence of concentration on the lifetimes determined for OCP_R27L-Ntag_, but a two-fold increase in that of the third component τ_ci-3_ for OCP_wt-Ctag_. This result establishes that the presence of dimers influences component τ_3_ in a concentration dependent fashion, but neither τ_1_ or τ_2_. A plot of the τ_3_ lifetime as a function of dimer concentration, using for zero concentration the average value from OCP_R27L-Ntag_ measurements at the 12, 50 and 250 µM (*i.e.* 54.8 ± 7.7 ms), shows a logarithmic shape, with a dimer concentration of ∼ 10 µM (corresponding to an overall OCP_wt-Ctag_ concentration of ∼ 20 µM) at half maximum (Fig. S10*B*). The data also show that the presence of dimers does not benefit the overall OCP^R^_wt-Ctag_ yield, despite the yield of the initial P_2_ intermediate rising with increased OCP_wt-Ctag_ concentration (Fig. S10*A*). To obtain further insights into the enthalpic and entropic contributions to the formation of each of the three states, we furthered our analysis by carrying out experiments at various temperatures in the 8-36 °C range (Fig. 5*B* and Fig. S9). Increased temperatures were associated to a lower yield of the red states, including the final OCP^R^_wt-Ctag_, possibly due to non-adiabatic processes occurring earlier than our first data point (∼ 50 ns) which do not repopulate directly the dark OCP^O^_wt-Ctag_ state (monitored at 440 nm). Another interpretation could be that the molar extinction coefficient of the OCP^R^-bound carotenoid decreases with increasing temperature. Irrespectively, we found that in both OCP_wt-Ctag_ and OCP_R27L-Ntag_, the first intermediate – whose associated lifetime hardly varies among proteins and concentrations – is adiabatically formed, *i.e.* with no exchange of heat (Fig. 5*E* and 5*F*). Thus, all the energy necessary for the formation of this intermediate is funneled into the protein scaffold before 50 ns, *i.e.* the first data point in our ns-s time span. The fact that no enthalpic contribution is associated with this component indicates that no H-bond or salt-bridge forms nor breaks upon accumulation of the corresponding intermediate, while the observation of a negative entropic contribution suggests a reduction in the complexity of the system. Hence, component τ_1_ could underlie the multi-step translocation of the carotenoid from the NTD/CTD interface into the NTD. As to the two other components, τ_2_ and τ_3_, they are associated with similar enthalpies of activation (∼ 18 kcal.mol^-1^) in OCP_wt-Ctag_, whereas a slightly larger value is found for τ_3_ (∼ 26 kcal.mol^-1^) as compared to τ_2_ (∼ 20 kcal.mol^-1^) in OCP_R27L-Ntag_. This difference in enthalpy could be rooted in the absence of dimers in the OCP_R27L-Ntag_ sample, as we have seen that the τ_3_ component – but not the τ_2_ component – is affected by the presence of these.

Time-resolved SAXS/WAXS on the 15 µs – 200 ms timescale allowed us to shed partial light on the structural changes associated with the OCP photocycle. Pulsed illumination results in a negative peak at ∼ 0.03 Å^-1^ in the ‘light’-minus-‘dark’ ΔI(q) profile, with this peak being already present at 15 µs, maximizing before 1 ms, and starting to decrease as of ∼10 ms and 100 ms in OCP_wt-Ntag_ (lifetime ∼70 ms) and OCP_wt-Ctag_ (lifetime ∼280 ms), respectively. Hence, our data demonstrate that the first large scale conformational changes associated with OCP photoactivation take place on the tens of µs time scale (15 µs), and that the localization of the his-tag influences the structural recovery rate. Both transient absorption spectroscopy and TR-SAXS required to work at comparatively high OCP concentration, with more than 98 % dimers at the highest concentration tested in spectroscopy experiments or used in TR-SAXS/WAXS experiments, and a minimum of ∼ 46 % dimers at the lowest concentration tested in spectroscopy experiments. Hence, we probed, with the two techniques, the effect on OCP dimers of ns pulses illumination, from the structural and the spectral perspectives. The TR-SAXS/WAXS data is indeed sensitive to structural changes regardless of where they occur in the protein, but is too low-resolution to afford information on the carotenoid position and on non-predominant species. This is at variance with our spectroscopy data which may inform on the formation of all red-shifted intermediates – productive (OCP^R^, offset at 0.5 s) and unproductive alike (Table S8) – but is blind to structural changes that do not affect the local environment of the carotenoid (e.g., detachment of NTE and CTT) (14, 54). As structural and spectroscopic transitions may be asynchronous, they should be considered independently.

Given that the probability that two monomers in a dimer can be activated by the same pulse is the square of the quantum yield, i.e. < 0.4 x10^-5^, our TR-SAXS could not probe the formation of OCP^R^, but only that of the predominant non-productive intermediate state. With keeping this in mind, we can propose the following sequence of events for the OCP photocycle based on ours as well as others results (Fig. 8). Upon absorption of the actinic photon, a S_2_ state forms which decays within ps into at least three states: the S_1_ and ICT excited states, and S*, which could either be an excited state (27) or a vibrationally-hot ground state (59). Within 20 ps, 99.3 % of molecules are back to the ground state and do not progress further towards OCP^R^. Our TR-SAXS/WAXS experiment reveals the existence of a ‘numbed’ OCP intermediate, OCP^NS^, which is formed upon such non-productive laser excitation and remains up to the µs time scale in a non-re-excitable structural state (Fig. 6*C* and 8A). Evidence for the existence of this state was obtained serendipitously, as we attempted to increase the extent of photoactivation by distribution of actinic photons in a longer pulse (5 vs 450 ns pulses). This attempt was infructuous, indicating that the ground state OCP^O^ structure formed after excitation remains non-photoactivatable for at least 0.5 µs, in line with the previous report that increasing the power of ns pulses does not result in a higher photoactivation yield (13). By use of a multi-pulse approach, whereby re-excitation was triggered every 1 ms for up to 30 pulses, we were able to estimate the lifetime of OCP^NS^ to lie between 450 ns and 1 ms (Fig. 6B, *C*).

**Figure 8.**
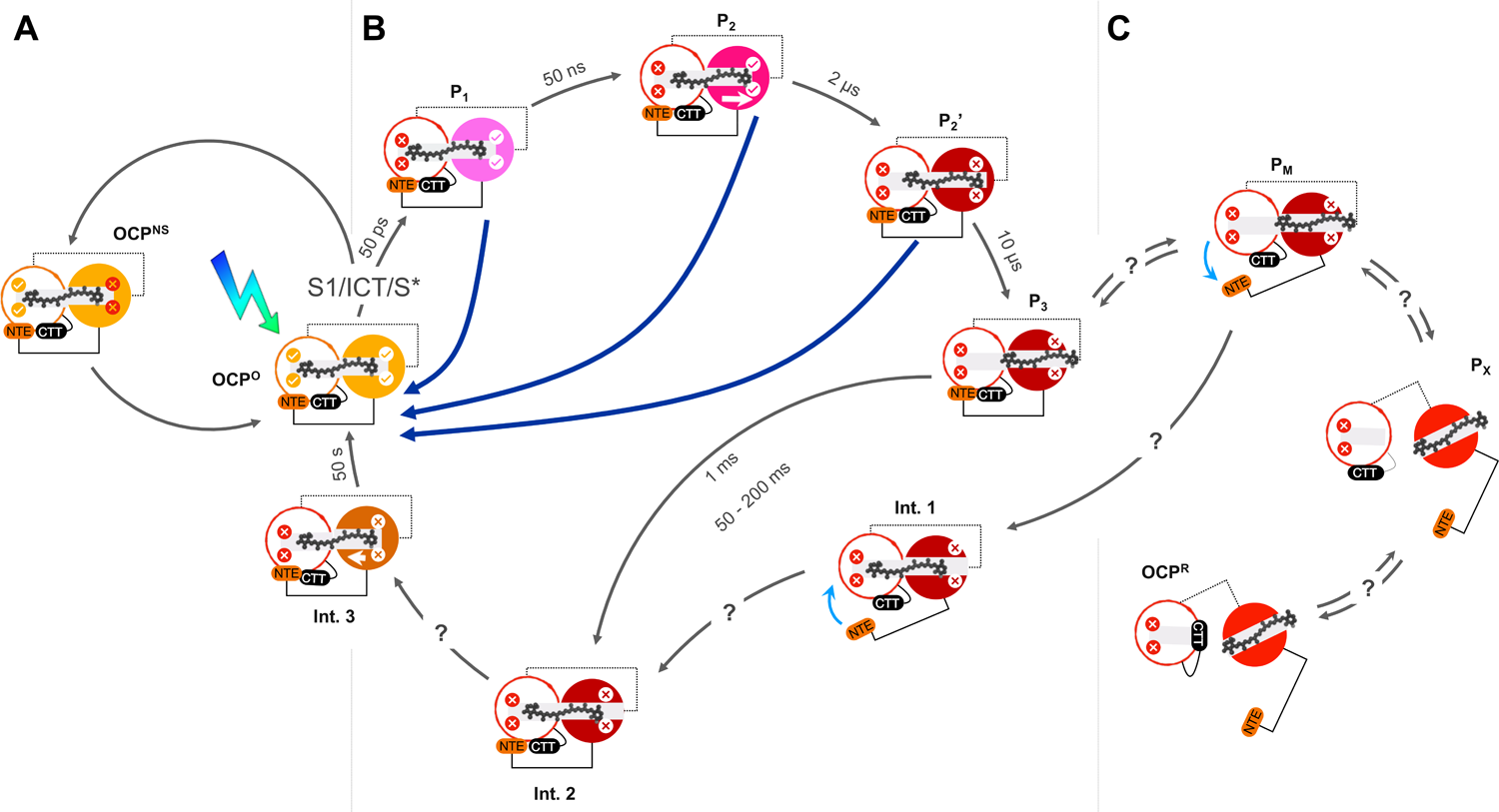
Proposed model for OCP photoactivation and recovery upon pulsed excitation. OCP is schematically represented with its N-terminal (coloured in orange for OCP^O^, pink for P_1_, magenta for P_2_, and red for P_2’_, P_3_, OCP^R^, Int.1, Int.2 and Int.3) and C-terminal domains (coloured in white). The ground state keto-carotenoid is shown in grey. Ticks and crosses in the CTD indicate presence and absence of H-bonds between the carotenoid and CTD residues Tyr201 and Trp288. Ticks and crosses in the NTD indicate presence and absence of π-stacking interactions between the carotenoid and NTD residues Tyr44 and Trp110. In absence of the former, the carotenoid may partially see the bulk solvent, since Tyr44 is a gating residue for a side channel to the carotenoid tunnel (ref. BBA). Complete migration of the carotenoid across the NTD tunnel requires the unsettling of interactions with Tyr44 and Trp110 and passage of a bottleneck contributed by NTD residues Leu37, Met83 and Met117. *(A)* Non-productive photoexcitation does not allow recovery of a photo-excitable state; instead a numbed-state forms, OCP^NS^, which remains in a non-excitable structural-state for at least 0.5 µs. *(B)* Productive photoexcitation yields P_1_, after rupture of H-bonds to the carotenoid. Past 50 ns, the carotenoid repositions at the NTD/CTD interface (P_2_), before bridging a first bottleneck at 2-3 µs (P_2’_; _;_ 0.5-1 and 0.4 µs, according to (27) and (54), respectively). Conformational changes occur in the NTD domain, enabling the carotenoid to pass across the NTD wall and its β2 ionone ring to become exposed to the solvent (P_3_; _;_ 10 and 15 µs, according to (27) and our TR-SAXS/WAXS data, respectively). At all steps before P_3_, back-migration of the carotenoid to its dark-state position can occur with rapid recovery of the orange state, as the tunnel remains shielded from the bulk (Fig. 5 and Fig. S9). After P_3_, recovery yet follows another pathway that does not enable rapid recovery of the orange state spectrum. (C) We presume that in the fraction of P_3_ that does not recover and further advances towards the OCP^R^ state, the NTE will detach from the CTD β-sheet, forming P_M_. Dissociation of the two domains thereafter becomes possible, yielding P_X_. The CTT may next reposition itself on the CTD, enabling to shield residues contributing to the carotenoid tunnel in OCP^O^ from the bulk, and thereby forming the minute-lived, photoactive OCP^R^. Recovery of OCP^O^ from OCP^R^ will have to await unbinding of the CTT from the carotenoid tunnel residues and reassociation of the two domains. It yet remains unclear whether the next step will be back migration of the carotenoid into the tunnel or reattachment to the CTD of the NTE. In the latter case, P_3_ would serve as an intermediate along the OCP^R^ to OCP^O^ recovery. In the former case, another pathway would exist whereby the carotenoid first back-migrates into the carotenoid tunnel (Int. 1) and then only does the NTE reattach to the CTD (Int. 2). The latter intermediate Int. 2 would exist in the two pathways. It seems likely that another intermediate would exist, which we refer to as Int. 3, given that even after 500 ms, the P_3_/OCP^O^ dimer has not reverted to the starting OCP^O^ state. This state would share all structural features of the OCP^O^ state albeit H-bonding of the carotenoid to CTD residues Y201 and W288.

Specific to the photo-productive pathway (Fig. 8B), it has been proposed that S* is the precursor of the first red photoproduct P_1_, characterized by rupture of the H-bonds between the carotenoid and the protein scaffold and a lifetime of ≈ 50 ns. The carotenoid therefrom debuts its migration from the NTD/CTD interface into the NTD, first forming photoproduct P_2_, after repositioning at the NTD/CTD interface, and then P_2_’, after further migration across the NTD. From the structural perspective, and as noted already by Konold et al. (27), this migration of the carotenoid must be accompanied by changes in the relative orientation of helices αC, αE and αG, which harbor the carotenoid-tunnel bottleneck-residues L37, M83 and M117. Eventually, these changes will propagate to the αG-αH loop and to E34, at the kink between αB and αC, resulting in an opening of the carotenoid tunnel across the NTD wall and complete migration of the carotenoid pigment, forming P_3_, wherein the β2 ionone ring of the carotenoid is exposed to the solvent – possibly as in the structure of the isolated NTD with canthaxanthin (19). We suppose that the first steps identified in our transient absorption spectroscopy (τ_1_ ∼2-3 µs) and TR-SAXS/WAXS(< 15 µs) experiments correspond to the P_2_/P_2_’ to P_3_ transition, in line with Konold et al. (0.5-1 and 10 µs for P_2_ to P_2_’ and P_2_’ to P_3_ transitions, respectively) (27). In the TR-SAXS/WAXS experiments, however, it is clear that this transition can only occur within OCP^O^ dimers. Hence, a partly-activated dimer would form, featuring a P_3_ monomer and a dark-adapted monomer (*i.e.* P_3_/OCP^O^ dimer). Failure to fully migrate across the carotenoid tunnel and form P_3_ is presumably sanctioned by rapid back-migration and re-binding to H-bonding partners in the CTD (component τ_1_), *i.e.* partial recovery of OCP^O^ from P_2_/P_2_’, as supported by the fact that the decrease in ΔA565 nm is mirrored by an increase in ΔA440 nm (Fig. S9). The next spectroscopically-visible steps (τ_2_, τ_3_) occur on the ∼1 ms and ∼50-200 ms time scales, respectively, and they are characterized by a large decrease in ΔA565 nm that is not accompanied by a similarly-extended increase in ΔA440 nm. Hence, only limited spectral recovery of the OCP^O^ state occurs on this time-scale. Remarkably, the τ_2_ and τ_3_ components display the same activation energy, suggesting that they would inform a similar molecular event, but only the latter is influenced by concentration. TR-SAXS data indicate that no structural changes other than reversion to the OCP^O^ state occurs up to 200 ms, excluding that the τ_2_ and τ_3_ components would inform on further progression of the P_3_/OCP^O^ dimers towards the OCP^R^ state – e.g. detachment of NTE from the CTD, which would be a prerequisite to dimer dissociation that itself would precede separation of the two domains. Hence, we propose that the τ_2_ and τ_3_ components both sign for the back migration of the carotenoid into the carotenoid tunnel of the P_3_ monomer in the P_3_/OCP^O^ dimer, but that each component informs on this process in a structurally-different partly-activated dimer. It has indeed been shown that two types of OCP^O^ exist, *i.e.* a red-shifted and a blue-shifted OCP^O^ (13). It is reasonable to propose that OCP dimerization would influence the equilibrium between these two OCP^O^ states, with the less flexible state being favored in the dimer – explaining the longer lifetime before back-migration occurs. The τ_2_ and τ_3_ components could thus underlie back-migration of the carotenoid in two such OCP^O^.

Taken altogether, our steady-state and time-resolved optical-spectroscopy and X-ray scattering data point to the conclusion that formation of P_3_ must occur in each of the two monomers, and the P_3_/P_3_ dimer dissociate into its monomeric counterparts, for OCP^O^ dimers to yield (monomeric) OCP^R^ (Fig 8*C*). Presumably, the next step in the photoactivation mechanism is the detachment from the CTD of αA and αN (associated with loss of 794 and 394 Å^2^ of buried surface area in the OCP^O^ monomer, respectively), which will rapidly be followed by dissociation of the two domains, given that the carotenoid has already fully migrated into the NTD (loss of 345 Å^2^ of buried surface area in the CTD) and does not contribute anymore to the stabilization of the closed-conformation (only 674 Å^2^ of buried surface area is left at the NTD/CTD interface). The lifetime of monomeric OCP^R^ is 50-100 s, but it may further associate into dimers or higher order oligomers upon increase in concentration, and these changes in quaternary structure will extend its lifetime to tens of minutes (lifetimes of 160-360 and 640 - 1327 s for OCP^R^ dimers or higher order oligomers, respectively) (Fig. 7). It is unclear if recovery of the OCP^O^ state from the OCP^R^ monomer channels back through the P_3_ intermediate (*i.e.* the NTD and CTD domains reassociate and αA and αN reattach to the CTD before back migration of the carotenoid into the tunnel) or follow another pathway, whereby after reassociation of the NTD and the CTD, back migration of the carotenoid into the tunnel occurs before re-attachment of αA and αN to the CTD (Fig. 8*C*). Reformation of an OCP^O^ dimer may ensue depending on concentration (Fig. 7). Given the low quantum yield of photoactivation, the existence of OCP^NS^, the back-migration of the carotenoid on the timescales of 1 and 50-200 ms in partly-activated P_3_/OCP^O^ dimers, and our finding that P_3_ starts recovering the structural OCP^O^ state on the time scale of ∼70 and ∼280 ms in N-tagged and C-tagged partly-activated dimers, respectively, the time window for absorption of an actinic photon by the second monomer in a partly-activated dimer is 0.5 µs – 200 ms. Any photon that arrives before 0.5 µs is set to be unproductive, while that which arrives after 200 ms will hit a dimer wherein both monomers are structurally back in the starting state (Figs. 7 and 8). Hence, productive photoactivation of dimers (meaning, accumulation of photoactive OCP^R^) is achievable only in experiments where one photon is delivered per chromophore every few ms. This is for example the case in our continuous illumination experiments where we estimate that 1 photon is delivered per chromophore every 10 - 30 ms upon illumination of a 23 mg/ml (620 µM) OCP solution with a 500 mW LED emitting at 430 nm. In hindsight, it must be acknowledged that a TR-SAXS experiment conducted on the constitutively-monomeric OCP_R27L Ntag_ mutant would have been more informative on functionally-relevant structural dynamics, as progression towards the OCP^R^ state would not have been impeded by presence of OCP^O^ dimers. Next experiments will focus on this variant and may enable tracking of the photoproductive OCP^R^ state.

## Conclusion

The emerging picture from our study suggests that oligomerization partakes in the regulation of the OCP photocycle, both at the level of OCP^O^ and OCP^R^. Whether or not these findings have implications regarding the biological function depends on the local concentration of OCP in the vicinity of cyanobacterial thylakoid membranes; yet the observation that the mutation of a single conserved amino-acid at the OCP^O^ dimerization interface results in monomerization of the protein is a strong argument in favor of a functional role for OCP^O^ dimers. The dissociation constant of OCP^O^ was estimated to be 14 µM, and that of OCP^R^ dimers suggested to be similar. Hence, OCP^R^ dimers could be functional. The dissociation constant of OCP^R^ higher-order oligomers is likely to fall in the 100-280 µM range, raising doubts as to their relevance for the physiological context. We also found that the first large-scale conformational changes occurring in photoactivated OCP take place on the µs time scale, likely corresponding to the P_2’_-P_3_ transition. We note that ours is the first study to offer time-resolved structural insights into the OCP photocycle with a non-resonant structural technique, and as well the first TR structural study to be carried out on a protein with such a low quantum yield. We hope our work attracts the attention of structural biologists on the possibilities offered by time-resolved SAXS/(WAXS).

## Author Contributions

J.-P.C. coordinated the project. E.A.A., G.S. and A.T. performed static X-ray scattering. E.A.A. analyzed static X-ray scattering data and performed thermal recovery assays. S.N., M.S. and G.B. performed time-resolved spectroscopy. S.N. analyzed time-resolved spectroscopy data. A.W. and D.K. performed protein expression and purification; E.A.A., N.Z. and R.M. prepared protein samples. E.D.Z. performed fitting analysis of thermal recovery data. J.-P.C., G.S. and M.L. designed the TR-SAXS experiment. E.A.A., S.N., M.L., F.M., G.B., M.S., G.S., J.-P.C. performed the TR-SAXS experiment. G.S. analyzed time-resolved X-ray scattering data; E.A.A. and J.-P.C. wrote the paper with input from other coauthors.

## Acknowledgments

We are grateful to Ilme Schlichting and Martin Weik for continued support of the project. We thank the SOLEIL and ESRF synchrotron radiation facilities for repeated allocation of beamtime on the SWING and ID09 beamlines, where the static and time-resolved SAXS experiments were performed, respectively. IBS acknowledges integration into the Interdisciplinary Research Institute of Grenoble (IRIG, CEA). This work was supported by the Agence Nationale de la Recherche (grants ANR-17-CE11-0018-01 and ANR-2018-CE11-0005-02 to J.-P.C.) and the Polish National Science Centre (NCN project 2018/31/N/ST4/03983), and used the platforms of the Grenoble Instruct-ERIC center (ISBG; UMS 3518 CNRS-CEA-UGA-EMBL) within the Grenoble Partnership for Structural Biology (PSB). Platform access was supported by FRISBI (ANR-10-INBS-05-02) and GRAL, a project of the University Grenoble Alpes graduate school (Ecoles Universitaires de Recherche) CBH-EUR-GS (ANR-17-EURE-0003). The access and travel fees of G.B. and S.N. to ESRF were financed by the Polish Ministry of Science and High Education - decision number: DIR/WK/2016/19. R.M. is supported by a GRAL PhD fellowship (7C047GRAL).

## Competing interests

Authors declare no competing interests.

## Data and materials availability

All data are available in the main text or the supplementary materials. Raw SAXS data have been deposited in the SASDB under accession codes can be made available upon request. SASDNY9, SASDNZ9, SASDP22, SASDP32, SASDP42, SASDP52, SASDP62, SASDP72, SASDP82, SASDP92, SASDPA2, SASDPB2, SASDPC2, SASDPD2. See Table S5 for details on the corresponding datasets.

## Supplementary Figures captions

**Figure S1.** Molecular envelopes and proposed OCP^R^ models for dark-adapted and light-adapted OCP_wt-Ctag_ and OCP_R27L-Ntag_ at different concentrations. The molecular envelopes, determined *ab initio* based on experimental static SAXS data are shown for OCP_R27L-Ntag_ and OCP_wt-Ctag_ in their dark-adapted and light-adapted states at three concentrations. The proposed models are overlayed on the molecular envelopes. For the dark-adapted OCP^O^_R27L-Ntag_, the molecular envelopes are consistent with the presence of an OCP^O^ monomer, whereas for the dark-adapted OCP^O^_wt-Ctag_, the molecular envelopes are better fitted by the crystallographic OCP^O^ dimer (PDB id: 3MG1). At each concentration, the molecular envelopes determined for light-adapted OCP^R^_R27L-Ntag_ and OCP^R^_wt-Ctag_ are similar, and consistent with the presence of an OCP^R^ dimer, at 0.7 and 3.5 mg/ml, and a higher-order OCP^R^ oligomers at 10.5 mg/ml, e.g., a compact tetramer or a more extended trimer, wherein one monomer associated by the NTD interface would be more opened than those associated via the CTD.

**Figure S2. SREFLEX modeling of light-adapted OCP.** The red circles show the X-ray scattering profiles of OCP^R^_wt-Ctag_ and OCP^R^_R27L-Ntag_ at 3.5 and 10.5 mg/ml. Continuous curves (coloured in green) represent calculated profiles using SREFLEX normal mode analysis with different OCP light-adapted oligomeric forms. The resulting chi-squared of SREFLEX analyses are indicated.

**FIGURE S3. Dimensionless Kratky plots for dark-adapted and light-adapted OCP variants.** (*A, B)* Normalized Kratky profiles of dark-adapted OCP^O^_wt-Ctag_ (*A*) and OCP^O^_R27L-Ntag_ (*B*) at various concentrations. (*C, D*) Normalized Kratky profiles of light-adapted OCP^R^_wt-Ctag_ (*C*) and OCP^R^_R27L-Ntag_ (*D*) at various concentrations. Note that the Kratky plot for OCP^O^_wt-Ctag_ at 0.7 mg/ml does not show the expected profile for a folded protein likely due to the poor signal to noise ratio.

**Figure S4. Photoactivation of OCP at 30 mg/ml requires tens of minutes of illumination by a 430 nm LED operating at 500 mW.** The figure shows X-ray scattering difference signals collected on OCP_wt-Ctag_ at 30 mg/ml during continuous illumination by a 430 nm collimated light-source in a 2 mm thick capillary. The LED nominal power was first set at 250 mW, switched to 500 mW after ∼750 s and kept on during 750 s after which saturation of the signal change was reached. Data was collected at the ID09 beamline (ESRF, Grenoble). Accumulation of the red state required using a nominal LED power of 500 mW and an illumination time of at least 700 sec. In these conditions, it is estimated that 1 to 3 photons are delivered every 30 ms per chromophore.

**Figure S5. Structural recovery of OCP^O^_wt-Ctag_, following accumulation of OCP^R^_wt-Ctag_.** The plot shows the time evolution of the integrated intensity in the 0.05-0.5 Å^-1^ region of the X-ray difference profile ΔI(q) = I(q)_t_ – I(q)_t0_, with I(q)_t_ and I(q)_t0_ the scattering intensities at a generic time t after switching off the 430 nm light and before the 430 nm irradiation procedure, respectively. The data establish that after accumulation of the OCP^R^ state at 23 mg/ml, recovery of the starting structural state of the OCP_wt-Ctag_ sample spans a time-scale of at least 2.5 hours.

**Figure S6.** The illumination procedure used to investigate OCP photoactivation and recovery does not significantly damage the protein sample. Absorption spectra recorded before (black) and after (green) 30 minutes illumination plus 16 hours of thermal recovery for (*A)* OCP_wt-Ntag_ at 3 mg/ml, (*B*) OCP_R27L-Ntag_ at 1.7 mg/ml and (*D*) OCP_wt-Ctag_ at 1.7 mg/ml. The OCP concentration and optical pathlength were 3 mg/ml and 0.1 cm, respectively.

**Figure S7. Global fitting of the recovery data recorder for the different OCP variants.** (*A*, *B*) Thermal recovery was monitored at four concentrations of OCP_R27L-Ntag_ (*A*, *B*), OCP_wt-Ntag_ (*C*, *D*) and OCP_wt-Ctag_ (*E*, *F*), and data were globally fitted for each variant using a model accounting either for three (*A, C, E*) or four (*B, D, F*) exponential components. Fitting parameters are reported in Table S6 and S7.

**Figure S8.** Kinetic analysis of OCP thermal recovery data at various concentrations suggests that oligomerized OCP^R^ reverts more slowly to the dark-adapted state. Plots of the pre-exponential factors retrieved from fits of thermal recovery kinetics (see also Fig. S7, Table S6 and Table S7) when four exponential components are used to account for the change in absorbance at 467 nm of OCP_R27L-Ntag_ (*A*) and OCP_wt-Ntag_ (*B*) data, and three exponential components for that of OCP_wt-Ctag_ (*C*). The apparent lifetime of OCP^R^ state is increased in dimers and higher-order oligomers, suggesting that oligomerization stabilizes the light-adapted state.

**Figure S9. Nanosecond transient absorption kinetics recorded on OCP_wt-Ctag_ at different temperatures.** (*A*-*E*) Difference absorption was monitored at both 440 and 565 nm, enabling to probe photoactivation and recovery at the blue and red edges of OCP^O^ and OCP^R^ absorption spectra, respectively. (*F*) Nanosecond transient absorption data recorded at 565 nm for OCP_wt-Ctag_ at various temperatures, from 8 to 36 °C. See Table S4 for complementary data.

**Figure S10.** OCP dimerization increases the yields at shorter time delays and delays the occurrence of the last recovery step in the ns-s spectral evolution. (*A*) Estimated yields of OCP_wt-Ctag_ red-states at various time delays and concentrations. The plot indicates that dimerization benefits the P_1_ and P_3_ yields. The OCP^R^ yield, however, only improves slightly. (*B*) The lifetime for the third component used to fit the ns-s spectral evolution of OCP_wt-Ctag_ increases as a function of OCP-dimer concentration. The average lifetime extracted from fits of the OCP_R27L-Ntag_ data at various concentrations was used for zero dimer concentration. (*C, D*) Predicted fraction of OCP^O^ dimers as a function of concentration (*C*) or optical density (OD) at 490 nm (*D*) based on a dissociation constant (K_d_) of 14 µM for OCP^O^ dimers.

**Figure S11. Size exclusion chromatography assesses the purity and dispersity of our OCP samples.** (*A*) At all tested concentration, the OCP^O^_R27L-Ntag_ sample is monodisperse and features a monomer. (*B*) Contrastingly, the OCP^O^_wt-Ctag_ sample features monomers and dimers.

**Figure S12. SVD analysis.** Python-based script for singular value decomposition (SVD) analysis of time-resolved (TR) difference profiles measured at ID09 (ESRF, Grenoble) in the µs-ms time-scale (See Fig. 6D).

## Supplementary Tables

**Table S1:** Data collection and processing parameters for static SAXS measurements recorded at the BioSAXS SWING beamline (Synchrotron SOLEIL, France)

**Table S2:** Structural parameters extracted using ATSAS from static SAXS analysis on dark-adapted (OCP^O^) OCP_wt-Ctag_ and OCP_R27L-Ntag_ at various concentrations.

**Table S3:** Structural parameters extracted using ATSAS from static SAXS analysis on light-adapted (OCP^R^) OCP_wt-Ctag_ and OCP_R27L-Ntag_ at various concentrations.

**Table S4:** Modeling parameters retrieved from static SAXS analysis on dark-adapted (OCP^O^) and light-adapted (OCP^R^) OCP_wt-Ctag_ and OCP_R27L-Ntag_ at various concentrations.

**Table S5:** Fitting parameters retrieved from static SAXS analysis on dark-adapted (OCP^O^) and light-adapted (OCP^R^) OCP_wt-Ctag_ and OCP_R27L-Ntag_ at various concentrations.

**Table S6:** Retained fitting parameters for the spectral (absorbance at 467 nm) recovery of OCP_wt-Ctag_, OCP_wt-Ntag_ and OCP_R27L-Ntag_, following accumulation of their OCP^R^ counterparts.

**Table S7:** Alternative Fitting parameters for the spectral (absorbance at 467 nm) recovery of OCP_wt-Ctag_, OCP_wt-Ntag_ and OCP_R27L-Ntag_, following accumulation of their OCP^R^ counterparts.

**Table S8: Fitted parameters extracted from nanosecond transient absorption data.** Percentage contributions in the table are calculated as a given pre-exponential factor divided by a_1_+ a_2_+ a_3_+ a_4_. Measurement of C-tag at 22 °C was repeated twice to confirm repeatability (note replicated row in the table).

---

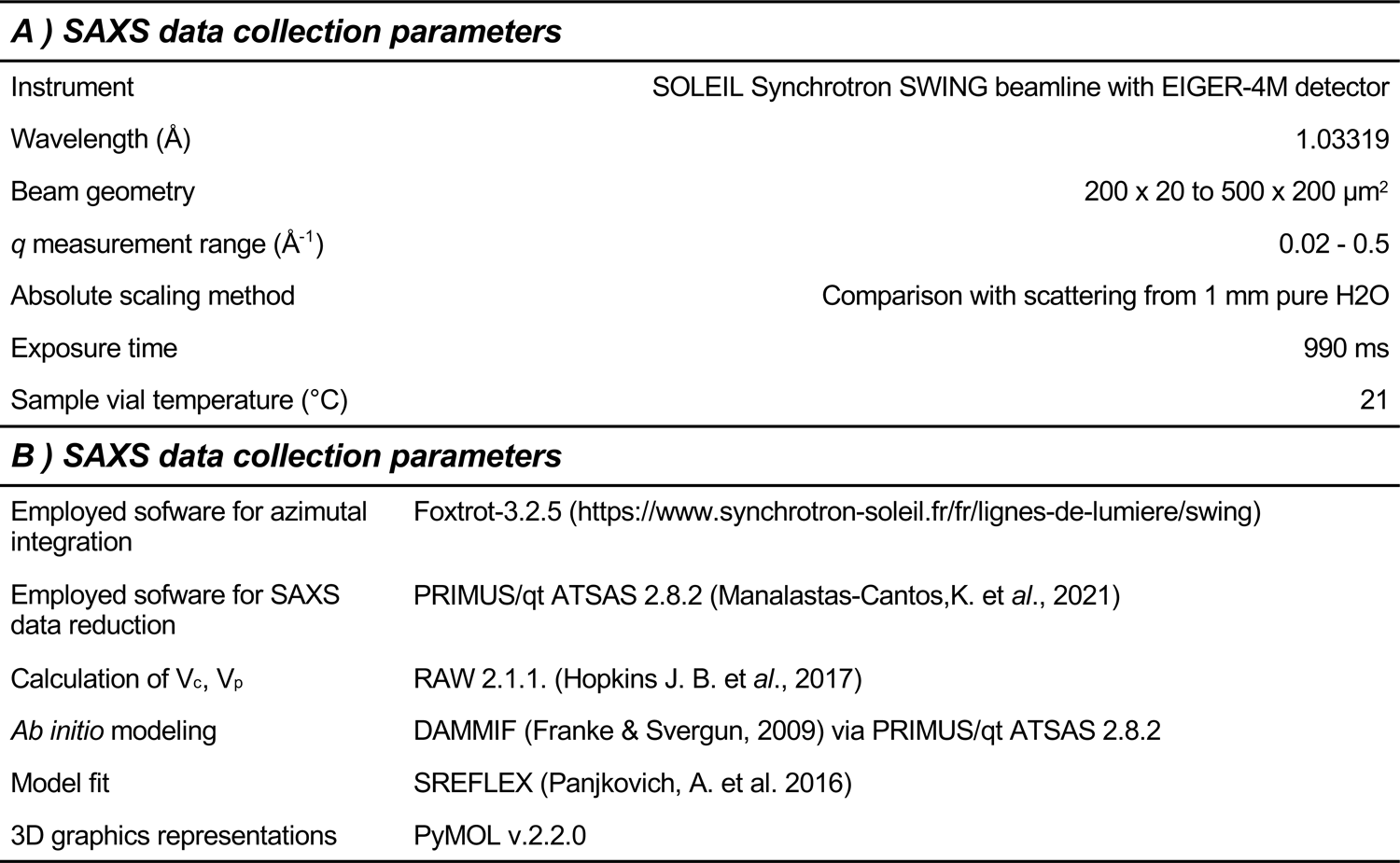

|  |  |
| --- | --- |
| Instrument | SOLEIL Synchrotron SWING beamline with EIGER-4M detector |
| Wavelength (Å) | 1.03319 |
| Beam geometry | 200 x 20 to 500 x 200 $\mu\text{m}^2$ |
| $q$ measurement range ($\text{\AA}^{-1}$) | 0.02 - 0.5 |
| Absolute scaling method | Comparison with scattering from 1 mm pure H<sub>2</sub>O |
| Exposure time | 990 ms |
| Sample vial temperature (°C) | 21 |
| **B ) SAXS data collection parameters** |  |
| Employed software for azimuthal integration | Foxtrot-3.2.5 (https://www.synchrotron-soleil.fr/fr/lignes-de-lumiere/swing) |
| Employed software for SAXS data reduction | PRIMUS/qt ATSAS 2.8.2 (Manalastas-Cantos, K. et al., 2021) |
| Calculation of $V_c$, $V_p$ | RAW 2.1.1. (Hopkins J. B. et al., 2017) |
| *Ab initio* modeling | DAMMIF (Franke & Svergun, 2009) via PRIMUS/qt ATSAS 2.8.2 |
| Model fit | SREFLEX (Panjkovich, A. et al. 2016) |
| 3D graphics representations | PyMOL v.2.2.0 |

| **Structural parameters of dark-adapted OCP (OCP<sup>0</sup>)** |  |  |  |  |  |  |  |  |
| --- | --- | --- | --- | --- | --- | --- | --- | --- |
| Protein variant | **OCP<sub>wt-Ctag</sub>** |  |  |  | **OCP<sub>R27L-Ntag</sub>** |  |  |  |
| Conc. (mg/ml) | 0.7 | 3.5 | 10.5 | 23 | 0.7 | 3.5 | 10.5 | 14 |
| **Guinier analysis** |  |  |  |  |  |  |  |  |
| $I(0)$ (cm<sup>-1</sup>) | 0.00824 ± 0.0001 | 0.0533 ± 0.00008 | 0.1771 ± 0.0002 | 0.3552 ± 0.0004 | 0.00727 ± 0.00002 | 0.0487 ± 0.00003 | 0.1183 ± 0.00009 | 0.1444 ± 0.00007 |
| $R_g$ (Å) | 24.35 ± 0.77 | 28.27 ± 0.16 | 28.02 ± 0.05 | 28.13 ± 0.01 | 22.24 ± 0.19 | 24.2 ± 0.04 | 22.93 ± 0.08 | 23.48 ± 0.02 |
| $q$ min (Å<sup>-1</sup>) | 0.02 | 0.02 | 0.02 | 0.02 | 0.02 | 0.02 | 0.02 | 0.02 |
| $q$ Rg max | 1.2931 | 1.2912 | 1.2954 | 1.2984 | 1.2950 | 1.2915 | 1.2943 | 1.2918 |
| $R^2$ | 0.8483 | 0.9979 | 0.9986 | 0.9995 | 0.9928 | 0.9995 | 0.9994 | 0.9994 |
| **P (r) analysis** |  |  |  |  |  |  |  |  |
| $I(0)$ (cm<sup>-1</sup>) | 0.00789 ± 0.00006 | 0.0532 ± 0.00007 | 0.1765 ± 0.0001 | 0.3536 ± 0.00006 | 0.0073 ± 0.00002 | 0.0484 ± 0.00002 | 0.1177 ± 0.00008 | 0.1436 ± 0.00007 |
| $R_g$ (Å) | 24.68 ± 0.17 | 27.9 ± 0.05 | 28.48 ± 0.03 | 28.17 ± 0.005 | 22.58 ± 0.06 | 24.19 ± 0.01 | 22.76 ± 0.02 | 23.29 ± 0.01 |
| $d_{max}$ | 71 | 90.2 | 93 | 90 | 68 | 76 | 75.2 | 76 |
| $q$ range | 0.02 – 0.5 | 0.02 – 0.5 | 0.02 – 0.5 | 0.02 – 0.5 | 0.02 – 0.5 | 0.02 – 0.5 | 0.02 – 0.5 | 0.02 – 0.5 |
| MW (kDa) from $I(0)$ | n/a | 45.6 | 58 | 25 | 35.1 | 36.4 | 36.8 | 36.5 |
| MW (kDa) from Vp | 37.1 | 56.2 | 63.4 | 63.1 | 34.9 | 40.8 | 38.3 | 39 |
| MW (kDa) from Vc | 37 | 52.9 | 59.6 | 57.8 | 37.1 | 37 | 36.5 | 37 |
| Calculated $R_g$ (Å) monomer | | | | 21.72 | | | | |
| Calculated $R_g$ (Å) dimer | | | | 27.16 | | | | |

| **Structural parameters of light-adapted (OCP<sup>R</sup>)** |  |  |  |  |  |  |
| --- | --- | --- | --- | --- | --- | --- |
| Protein variant | OCP<sub>wt-Ctag</sub> |  |  | OCP<sub>R27L-Ntag</sub> |  |  |
| Conc. (mg/ml) | 0.7 | 3.5 | 10.5 | 0.7 | 3.5 | 10.5 |
| **Guinier analysis** |  |  |  |  |  |  |
| *I*(0) (cm<sup>-1</sup>) | 0.00811 ± 0.000063 | 0.00549 ± 0.00022 | 0.4975 ± 0.00204 | 0.00921 ± 0.00007 | 0.0605 ± 0.00018 | 0.2817 ± 0.00100 |
| R<sub>g</sub> (Å) | 29.11 ± 0.35 | 34 ± 0.2 | 43.8 ± 0.04 | 32.5 ± 0.23 | 34.15 ± 0.09 | 48 ± 0.2 |
| *q* min (Å<sup>-1</sup>) | 0.02 | 0.02 | 0.02 | 0.02 | 0.02 | 0.02 |
| *q* R<sub>g</sub> max | 1.2808 | 1.2936 | 1.2870 | 1.2915 | 1.2869 | 1.2918 |
| R<sup>2</sup> | 0.9674 | 0.9924 | 0.9975 | 0.9949 | 0.9969 | 0.9971 |
| **P (r) analysis** |  |  |  |  |  |  |
| *I*(0) (cm<sup>-1</sup>) | 0.00835 ± 0.000063 | 0.0056 ± 0.0021 | 0.5571 ± 0.00216 | 0.00943 ± 0.00005 | 0.0628 ± 0.0014 | 0.3192 ± 0.0031 |
| R<sub>g</sub> (Å) | 31.59 ± 0.6 | 36.05 ± 0.2 | 59.48 ± 0.41 | 35.13 ± 0.3 | 34.70 ± 0.1 | 61.56 ± 0.8 |
| d<sub>max</sub> | 127 | 126 | 260 | 126 | 130 | 260 |
| *q* range | 0.02 – 0.5 | 0.02 – 0.5 | 0.02 – 0.5 | 0.02 – 0.5 | 0.02 – 0.5 | 0.02 – 0.5 |
| MW (kDa) from *I*(0) | 62 | 62.7 | 169.4 | 60 | 62.5 | 165 |
| MW from V<sub>p</sub> | 54.2 | 64.8 | 99.3 | 58.2 | 54.3 | 76.1 |
| MW from V<sub>c</sub> | 52.1 | 56.4 | 84.2 | 47.4 | 46.1 | 92.8 |

A) *Ab initio* modeling of dark-adapted OCP (OCP<sup>0</sup>)
| Protein variant | OCP<sub>wt-Ctag</sub> |  |  |  | OCP<sub>R27L-Ntag</sub> |  |  |  |
| --- | --- | --- | --- | --- | --- | --- | --- | --- |
| Conc. (mg/ml) | 0.7 | 3.5 | 10.5 | 23 | 0.7 | 3.5 | 10.5 | 14 |
| **DAMMIF (default parameters, 20 repetitions)** |  |  |  |  |  |  |  |  |
| *q* range (Å<sup>-1</sup>) | 0.02-0.156 | 0.02-0.135 | 0.02-0.227 | 0.02-0.279 | 0.02-0.357 | 0.02-0.357 | 0.02-0.357 | 0.02-0.357 |
| Symmetry | *P1* | *P1* | *P1* | *P1* | *P1* | *P1* | *P1* | *P1* |
| Mean value of NSD | 1.108 | 1.062 | 0.650 | 0.630 | 0.719 | 0.776 | 0.715 | 0.699 |
| χ<sup>2</sup> | 1.1 | 1.057 | 1.127 | 1.517 | 1.029 | 1.173 | 0.956 | 1.036 |
| constant adjustment | none | none | 5.676E-04 | 1.592E-03 | 5.336E-05 | 2.268E-04 | 3.020E-04 | 5.026E-04 |

B) *Ab initio* modeling of light-adapted (OCP<sup>R</sup>)
| Protein variant | OCP<sub>wt-Ctag</sub> |  |  | OCP<sub>R27L-Ntag</sub> |  |  |
| --- | --- | --- | --- | --- | --- | --- |
| Conc. (mg/ml) | 0.7 | 3.5 | 10.5 | 0.7 | 3.5 | 10.5 |
| **DAMMIF (default parameters, 20 repetitions)** |  |  |  |  |  |  |
| *q* range (Å<sup>-1</sup>) | 0.02-0.293 | 0.02-0.293 | 0.02-0.293 | 0.02-0.293 | 0.02-0.293 | 0.02-0.293 |
| Symmetry | *P1* | *P1* | *P1* | *P1* | *P1* | *P1* |
| Mean value of NSD | 1.324 | 0.849 | 0.718 | 0.573 | 0.927 | 0.765 |
| χ<sup>2</sup> | 0.995 | 1.022 | 1.124 | 1.035 | 1.135 | 1.024 |
| Constant adjustment | 4.125E-05 | 1.270E-04 | 1.303E-03 | none | 2.303E-04 | 5.046E-04 |

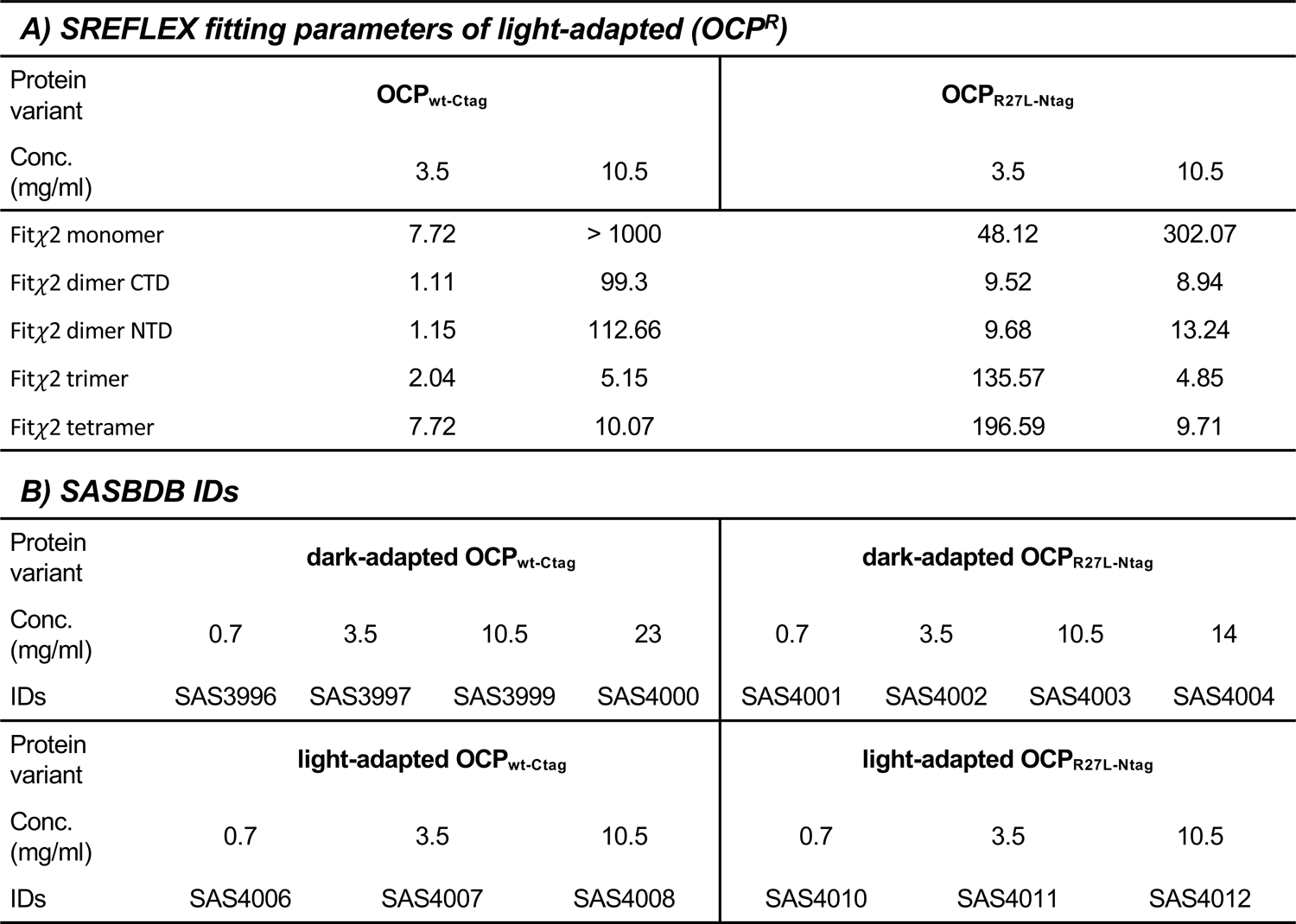

A) SREFLEX fitting parameters of light-adapted (OCP<sup>R</sup>)
| Protein variant | OCP<sub>wt-Ctag</sub> |  | OCP<sub>R27L-Ntag</sub> |  |
| --- | --- | --- | --- | --- |
|  | Conc. (mg/ml) |  | Conc. (mg/ml) |  |
|  | 3.5 | 10.5 | 3.5 | 10.5 |
| Fitχ2 monomer | 7.72 | > 1000 | 48.12 | 302.07 |
| Fitχ2 dimer CTD | 1.11 | 99.3 | 9.52 | 8.94 |
| Fitχ2 dimer NTD | 1.15 | 112.66 | 9.68 | 13.24 |
| Fitχ2 trimer | 2.04 | 5.15 | 135.57 | 4.85 |
| Fitχ2 tetramer | 7.72 | 10.07 | 196.59 | 9.71 |

B) SASBDB IDs
| Protein variant | dark-adapted OCP<sub>wt-Ctag</sub> |  |  |  | dark-adapted OCP<sub>R27L-Ntag</sub> |  |  |  |
| --- | --- | --- | --- | --- | --- | --- | --- | --- |
|  | Conc. (mg/ml) |  |  |  | Conc. (mg/ml) |  |  |  |
|  | 0.7 | 3.5 | 10.5 | 23 | 0.7 | 3.5 | 10.5 | 14 |
| IDs | SAS3996 | SAS3997 | SAS3999 | SAS4000 | SAS4001 | SAS4002 | SAS4003 | SAS4004 |
| Protein variant | light-adapted OCP<sub>wt-Ctag</sub> |  |  |  | light-adapted OCP<sub>R27L-Ntag</sub> |  |  |  |
|  | Conc. (mg/ml) |  |  |  | Conc. (mg/ml) |  |  |  |
|  | 0.7 | 3.5 | 10.5 |  | 0.7 | 3.5 | 10.5 |  |
| IDs | SAS4006 | SAS4007 | SAS4008 |  | SAS4010 | SAS4011 | SAS4012 |  |

| Spectroscopic recovery | fitting function: |  |  |
| --- | --- | --- | --- |
| | $a_{ci-1} \times [1 - \exp(-t/\tau_{ci-1})] + a_{ci-2} \times [1 - \exp(-t/\tau_{ci-2})] + a_{ci-3} \times [1 - \exp(-t/\tau_{ci-3})]$ | | |
| | $\tau_{ci-1}$ [s] | $\tau_{ci-2}$ [s] | $\tau_{ci-3}$ [s] |
|  | 59.825 ± 0.253 | 159.132 ± 0.594 | 641.708 ± 1.147 |
| OCP<sub>R27L-Ctag</sub> | $a_{ci-1}$ | $a_{ci-2}$ | $a_{ci-3}$ |
| 0.1 mg/ml | 0.899 ± 0.004 | 0.013 ± 0.005 | 0.084 ± 0.001 |
| 1.7 mg/ml | 0.023 ± 0.003 | 0.810 ± 0.003 | 0.156 ± 0.001 |
| 3.0 mg/ml | 0.000 ± 0.035 | 0.805 ± 0.003 | 0.190 ± 0.037 |
| 16.0 mg/ml | 0.000 ± 0.022 | 0.180 ± 0.003 | 0.820 ± 0.003 |

<sup>§</sup> $a_{ci-4}$ was constrained to $a_{ci-3} = (1 - a_{ci-1} - a_{ci-2} - a_{ci-3})$ $\chi^2 = 0.157$ ; reduced $\chi^2 = 2.242e-05$

| Spectroscopic recovery | fitting function: |  |  |
| --- | --- | --- | --- |
| | $a_{ci-1} \times [1 - \exp(-t/\tau_{ci-1})] + a_{ci-2} \times [1 - \exp(-t/\tau_{ci-2})] + a_{ci-3} \times [1 - \exp(-t/\tau_{ci-3})]$ | | |
| | $\tau_{ci-1}$ [s] | $\tau_{ci-2}$ [s] | $\tau_{ci-3}$ [s] |
|  | 94.513 ± 0.247 | 356.523 ± 1.246 | 1327.486 ± 4.338 |
| OCP<sub>wt-Ntag</sub> | $a_{ci-1}$ | $a_{ci-2}$ | $a_{ci-3}$ |
| 0.1 mg/ml | 0.909 ± 0.001 | 0.088 ± 0.002 | 0.000 ± 0.000 |
| 3.0 mg/ml | 0.194 ± 0.002 | 0.722 ± 0.002 | 0.083 ± 0.001 |
| 10.0 mg/ml | 0.093 ± 0.002 | 0.740 ± 0.002 | 0.167 ± 0.001 |
| 16.0 mg/ml | 0.000 ± 0.004 | 0.411 ± 0.003 | 0.584 ± 0.003 |

<sup>§</sup> $a_{ci-3}$ was constrained to $a_{ci-3} = (\text{maximum} - a_{ci-1} - a_{ci-2})$ $\chi^2 = 0.305$ ; reduced $\chi^2 = 3.261e-05$

| Spectroscopic recovery | fitting function: |  |  |  |
| --- | --- | --- | --- | --- |
| | $a_{ci-1} \times [1 - \exp(-t/\tau_{ci-1})] + a_{ci-2} \times [1 - \exp(-t/\tau_{ci-2})] + a_{ci-3} \times [1 - \exp(-t/\tau_{ci-3})] + a_{ci-4} \times [1 - \exp(-t/\tau_{ci-4})]$ | | | |
| | $\tau_{ci-1}$ [s] | $\tau_{ci-2}$ [s] | $\tau_{ci-3}$ [s] | $\tau_{ci-4}$ [s] |
|  | 51.491 ± 0.218 | 209.893 ± 1.288 | 945.989 ± 3.703 | 14057.656 ± 68.830 |
| OCP<sub>wt-Ctag</sub> | $a_{ci-1}$<sup>§</sup> | $a_{ci-2}$ | $a_{ci-3}$ | $a_{ci-4}$ |
| 0.1 mg/ml | 0.688 ± 0.002 | 0.234 ± 0.003 | 0.038 ± 0.001 | 0.039 ± 0.000 |
| 1.7 mg/ml | 0.382 ± 0.002 | 0.431 ± 0.002 | 0.140 ± 0.001 | 0.047 ± 0.000 |
| 3.0 mg/ml | 0.291 ± 0.002 | 0.462 ± 0.002 | 0.193 ± 0.001 | 0.053 ± 0.000 |
| 16.0 mg/ml | 0.063 ± 0.001 | 0.297 ± 0.002 | 0.533 ± 0.002 | 0.107 ± 0.000 |

<sup>§</sup> $a_{ci-4}$ was constrained to $a_{ci-4} = (\text{maximum} - a_{ci-1} - a_{ci-2} - a_{ci-3})$ $\chi^2 = 0.281$ ; reduced $\chi^2 = 1.93e-05$

| Spectroscopic recovery | fitting function: |  |  |  |
| --- | --- | --- | --- | --- |
| | $a_{ci-1} \times [1 - \exp(-t/\tau_{ci-1})] + a_{ci-2} \times [1 - \exp(-t/\tau_{ci-2})] + a_{ci-3} \times [1 - \exp(-t/\tau_{ci-3})] + a_{ci-4} \times [1 - \exp(-t/\tau_{ci-4})]$ | | | |
| | $\tau_{ci-1}$ [s] | $\tau_{ci-2}$ [s] | $\tau_{ci-3}$ [s] | $\tau_{ci-4}$ [s] |
|  | 9.402 ± 1.551 | 76.390 ± 1.121 | 257.289 ± 3.199 | 760.380 ± 8.141 |
| OCP<sub>R27L-Ctag</sub> | $a_{ci-1}^{\S}$ | $a_{ci-2}$ | $a_{ci-3}$ | $a_{ci-4}$ |
| 0.1 mg/ml | 0.081 ± 0.008 | 0.864 ± 0.007 | 0.000 ± 0.009 | 0.050 ± 0.002 |
| 1.7 mg/ml | 0.000 ± 0.003 | 0.343 ± 0.009 | 0.595 ± 0.007 | 0.052 ± 0.004 |
| 3.0 mg/ml | 0.000 ± 0.003 | 0.281 ± 0.009 | 0.651 ± 0.006 | 0.063 ± 0.004 |
| 16.0 mg/ml | 0.000 ± 0.001 | 0.000 ± 0.004 | 0.374 ± 0.011 | 0.627 ± 0.009 |

<sup>§</sup> $a_{ci-4}$ was constrained to $a_{ci-4} = (\text{maximum-}a_{ci-1}-a_{ci-2}-a_{ci-3})$ $\chi^2 = 0.824$ ; reduced $\chi^2 = 1.180\text{e-}04$

| Spectroscopic recovery | fitting function: |  |  |  |
| --- | --- | --- | --- | --- |
| | $a_{ci-1} \times [1 - \exp(-t/\tau_{ci-1})] + a_{ci-2} \times [1 - \exp(-t/\tau_{ci-2})] + a_{ci-3} \times [1 - \exp(-t/\tau_{ci-3})] + a_{ci-4} \times [1 - \exp(-t/\tau_{ci-4})]$ | | | |
| | $\tau_{ci-1}$ [s] | $\tau_{ci-2}$ [s] | $\tau_{ci-3}$ [s] | $\tau_{ci-4}$ [s] |
|  | 94.786 ± 0.336 | 225.655 ± 298.575 | 360.181 ± 10.521 | 1325.753 ± 7.249 |
| OCP<sub>wt-Ntag</sub> | $a_{ci-1}^{\S}$ | $a_{ci-2}$ | $a_{ci-3}$ | $a_{ci-4}$ |
| 0.1 mg/ml | 0.913 ± 0.003 | 0.000 ± 0.000 | 0.080 ± 0.003 | 0.004 ± 0.001 |
| 3.0 mg/ml | 0.188 ± 0.005 | 0.009 ± 0.068 | 0.729 ± 0.064 | 0.074 ± 0.002 |
| 10.0 mg/ml | 0.096 ± 0.010 | 0.026 ± 0.090 | 0.703 ± 0.097 | 0.175 ± 0.002 |
| 16.0 mg/ml | 0.005 ± 0.004 | 0.000 ± 0.028 | 0.407 ± 0.023 | 0.583 ± 0.005 |

<sup>§</sup> $a_{ci-4}$ was constrained to $a_{ci-4} = (\text{maximum-}a_{ci-1}-a_{ci-2}-a_{ci-3})$ $\chi^2 = 0.323$ ; reduced $\chi^2 = 3.45\text{e-}05$

| Spectroscopic recovery | fitting function: |  |  |
| --- | --- | --- | --- |
| | $a_{ci-1} \times [1 - \exp(-t/\tau_{ci-1})] + a_{ci-2} \times [1 - \exp(-t/\tau_{ci-2})] + a_{ci-3} \times [1 - \exp(-t/\tau_{ci-3})]$ | | |
| | $\tau_{ci-1}$ [s] | $\tau_{ci-2}$ [s] | $\tau_{ci-3}$ [s] |
|  | 76.669 ± 0.202 | 593.111 ± 2.088 | 9517.368 ± 75.252 |
| OCP<sub>wt-Ctag</sub> | $a_{ci-1}^{\S}$ | $a_{ci-2}$ | $a_{ci-3}$ |
| 0.1 mg/ml | 0.880 ± 0.001 | 0.072 ± 0.001 | 0.047 ± 0.000 |
| 1.7 mg/ml | 0.646 ± 0.001 | 0.300 ± 0.001 | 0.053 ± 0.000 |
| 3.0 mg/ml | 0.555 ± 0.001 | 0.382 ± 0.001 | 0.063 ± 0.000 |
| 16.0 mg/ml | 0.152 ± 0.001 | 0.688 ± 0.001 | 0.159 ± 0.001 |

<sup>§</sup> $a_{ci-3}$ was constrained to $a_{ci-3} = (\text{maximum-}a_{ci-1}-a_{ci-2})$ $\chi^2 = 1.170$ ; reduced $\chi^2 = 8.031\text{e-}05$

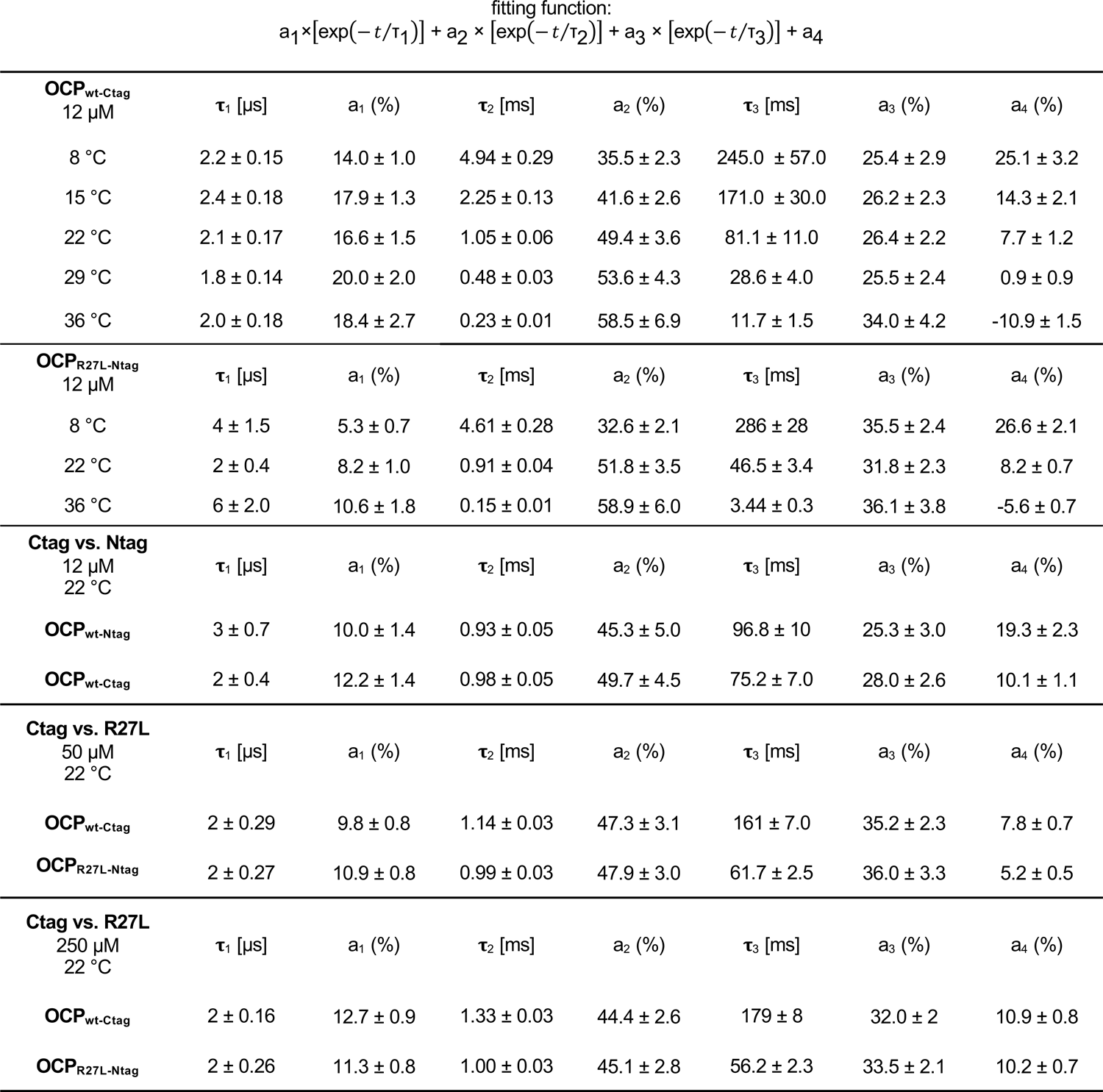

|  |  |  |  |  |  |  |  |
| --- | --- | --- | --- | --- | --- | --- | --- |
| **OCP<sub>wt-Ctag</sub>**<br>12 μM | **τ<sub>1</sub> [μs]** | **a<sub>1</sub> (%)** | **τ<sub>2</sub> [ms]** | **a<sub>2</sub> (%)** | **τ<sub>3</sub> [ms]** | **a<sub>3</sub> (%)** | **a<sub>4</sub> (%)** |
| 8 °C | 2.2 ± 0.15 | 14.0 ± 1.0 | 4.94 ± 0.29 | 35.5 ± 2.3 | 245.0 ± 57.0 | 25.4 ± 2.9 | 25.1 ± 3.2 |
| 15 °C | 2.4 ± 0.18 | 17.9 ± 1.3 | 2.25 ± 0.13 | 41.6 ± 2.6 | 171.0 ± 30.0 | 26.2 ± 2.3 | 14.3 ± 2.1 |
| 22 °C | 2.1 ± 0.17 | 16.6 ± 1.5 | 1.05 ± 0.06 | 49.4 ± 3.6 | 81.1 ± 11.0 | 26.4 ± 2.2 | 7.7 ± 1.2 |
| 29 °C | 1.8 ± 0.14 | 20.0 ± 2.0 | 0.48 ± 0.03 | 53.6 ± 4.3 | 28.6 ± 4.0 | 25.5 ± 2.4 | 0.9 ± 0.9 |
| 36 °C | 2.0 ± 0.18 | 18.4 ± 2.7 | 0.23 ± 0.01 | 58.5 ± 6.9 | 11.7 ± 1.5 | 34.0 ± 4.2 | -10.9 ± 1.5 |
| **OCP<sub>R27L-Ntag</sub>**<br>12 μM | **τ<sub>1</sub> [μs]** | **a<sub>1</sub> (%)** | **τ<sub>2</sub> [ms]** | **a<sub>2</sub> (%)** | **τ<sub>3</sub> [ms]** | **a<sub>3</sub> (%)** | **a<sub>4</sub> (%)** |
| 8 °C | 4 ± 1.5 | 5.3 ± 0.7 | 4.61 ± 0.28 | 32.6 ± 2.1 | 286 ± 28 | 35.5 ± 2.4 | 26.6 ± 2.1 |
| 22 °C | 2 ± 0.4 | 8.2 ± 1.0 | 0.91 ± 0.04 | 51.8 ± 3.5 | 46.5 ± 3.4 | 31.8 ± 2.3 | 8.2 ± 0.7 |
| 36 °C | 6 ± 2.0 | 10.6 ± 1.8 | 0.15 ± 0.01 | 58.9 ± 6.0 | 3.44 ± 0.3 | 36.1 ± 3.8 | -5.6 ± 0.7 |
| **Ctag vs. Ntag**<br>12 μM<br>22 °C | **τ<sub>1</sub> [μs]** | **a<sub>1</sub> (%)** | **τ<sub>2</sub> [ms]** | **a<sub>2</sub> (%)** | **τ<sub>3</sub> [ms]** | **a<sub>3</sub> (%)** | **a<sub>4</sub> (%)** |
| **OCP<sub>wt-Ntag</sub>** | 3 ± 0.7 | 10.0 ± 1.4 | 0.93 ± 0.05 | 45.3 ± 5.0 | 96.8 ± 10 | 25.3 ± 3.0 | 19.3 ± 2.3 |
| **OCP<sub>wt-Ctag</sub>** | 2 ± 0.4 | 12.2 ± 1.4 | 0.98 ± 0.05 | 49.7 ± 4.5 | 75.2 ± 7.0 | 28.0 ± 2.6 | 10.1 ± 1.1 |
| **Ctag vs. R27L**<br>50 μM<br>22 °C | **τ<sub>1</sub> [μs]** | **a<sub>1</sub> (%)** | **τ<sub>2</sub> [ms]** | **a<sub>2</sub> (%)** | **τ<sub>3</sub> [ms]** | **a<sub>3</sub> (%)** | **a<sub>4</sub> (%)** |
| **OCP<sub>wt-Ctag</sub>** | 2 ± 0.29 | 9.8 ± 0.8 | 1.14 ± 0.03 | 47.3 ± 3.1 | 161 ± 7.0 | 35.2 ± 2.3 | 7.8 ± 0.7 |
| **OCP<sub>R27L-Ntag</sub>** | 2 ± 0.27 | 10.9 ± 0.8 | 0.99 ± 0.03 | 47.9 ± 3.0 | 61.7 ± 2.5 | 36.0 ± 3.3 | 5.2 ± 0.5 |
| **Ctag vs. R27L**<br>250 μM<br>22 °C | **τ<sub>1</sub> [μs]** | **a<sub>1</sub> (%)** | **τ<sub>2</sub> [ms]** | **a<sub>2</sub> (%)** | **τ<sub>3</sub> [ms]** | **a<sub>3</sub> (%)** | **a<sub>4</sub> (%)** |
| **OCP<sub>wt-Ctag</sub>** | 2 ± 0.16 | 12.7 ± 0.9 | 1.33 ± 0.03 | 44.4 ± 2.6 | 179 ± 8 | 32.0 ± 2 | 10.9 ± 0.8 |
| **OCP<sub>R27L-Ntag</sub>** | 2 ± 0.26 | 11.3 ± 0.8 | 1.00 ± 0.03 | 45.1 ± 2.8 | 56.2 ± 2.3 | 33.5 ± 2.1 | 10.2 ± 0.7 |

## Notes

### Competing Interest Statement

The authors have declared no competing interest.

### Summary of Updates

Revised version after talking into account reviewers remarks.

