## Supplementary Figures for "Oligomerization processes limit photoactivation and recovery of the Orange Carotenoid Protein"

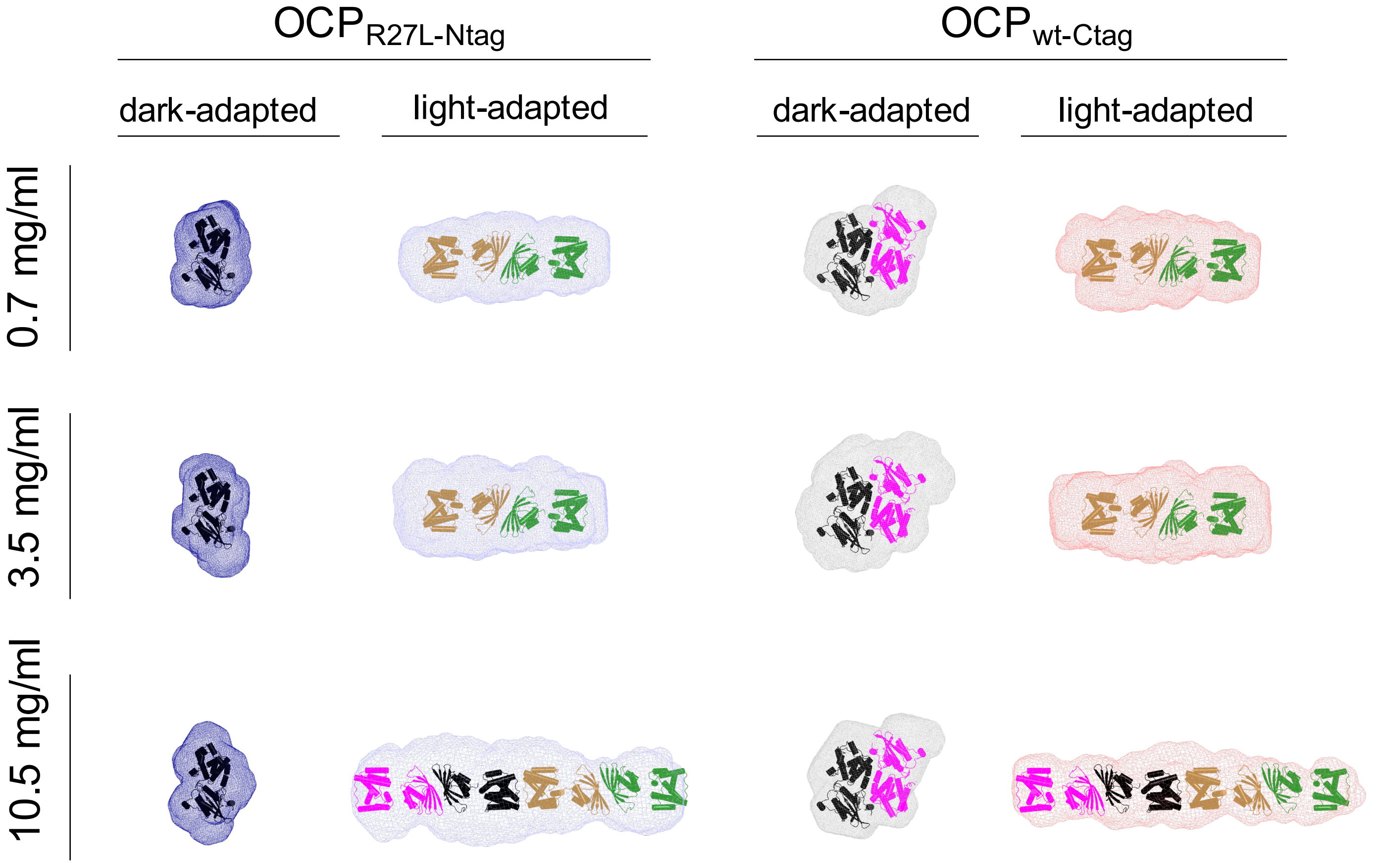
OCP$_{R27L-Ntag}$
OCP$_{wt-Ctag}$
dark-adapted
light-adapted
dark-adapted
light-adapted
0.7 mg/ml
3.5 mg/ml
10.5 mg/ml

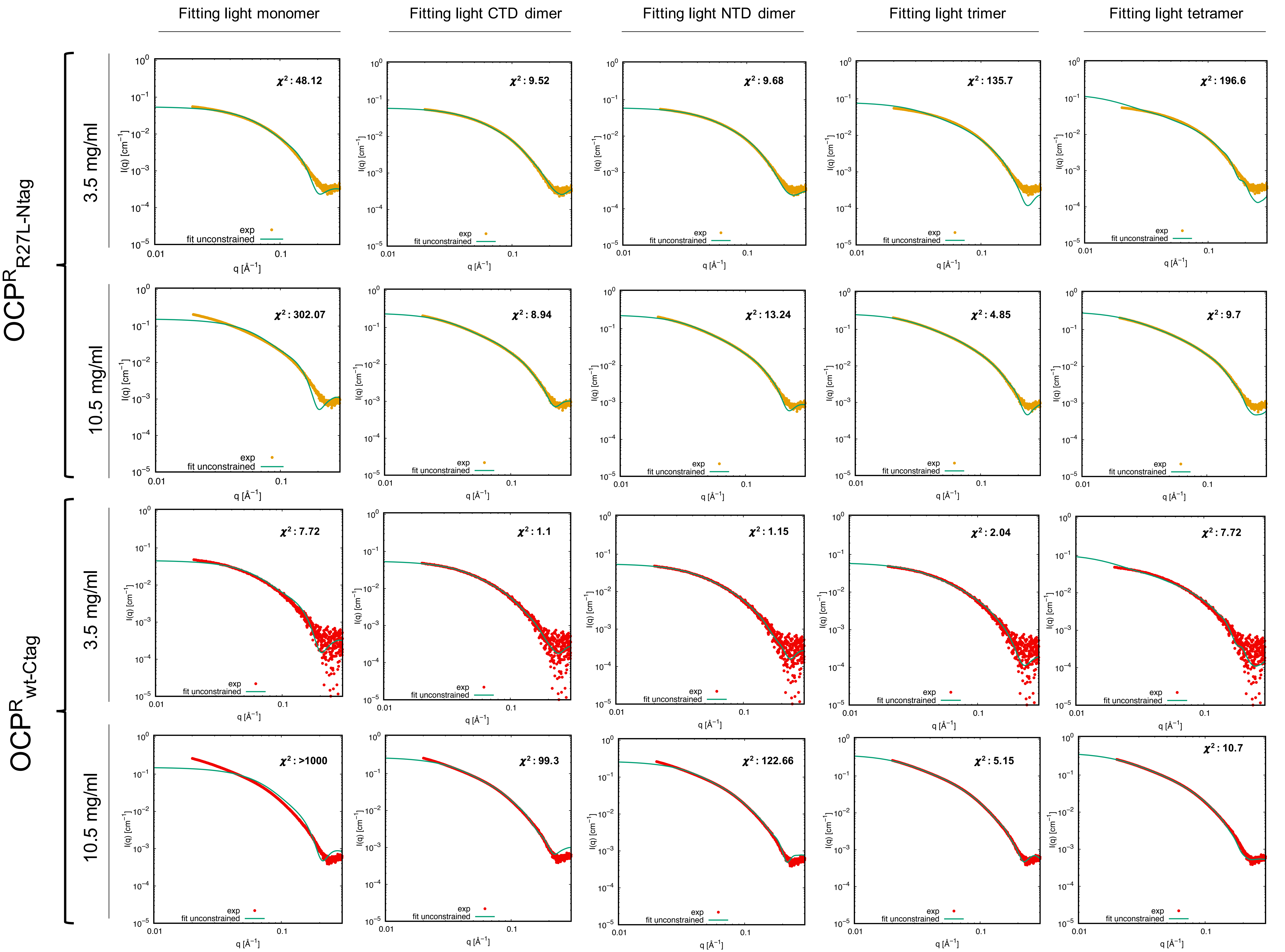

Fitting light monomer
Fitting light CTD dimer
Fitting light NTD dimer
Fitting light trimer
Fitting light tetramer
$OCP^R_{R27L\text{-}Ntag}$
$OCP^R_{wt\text{-}Ctag}$
3.5 mg/ml
10.5 mg/ml
3.5 mg/ml
10.5 mg/ml
$\chi^2$ : 48.12
$\chi^2$ : 9.52
$\chi^2$ : 9.68
$\chi^2$ : 135.7
$\chi^2$ : 196.6
$\chi^2$ : 302.07
$\chi^2$ : 8.94
$\chi^2$ : 13.24
$\chi^2$ : 4.85
$\chi^2$ : 9.7
$\chi^2$ : 7.72
$\chi^2$ : 1.1
$\chi^2$ : 1.15
$\chi^2$ : 2.04
$\chi^2$ : 7.72
$\chi^2$ : >1000
$\chi^2$ : 99.3
$\chi^2$ : 122.66
$\chi^2$ : 5.15
$\chi^2$ : 10.7
exp
fit unconstrained
I(q) [cm$^{-1}$]
q [Å$^{-1}$]

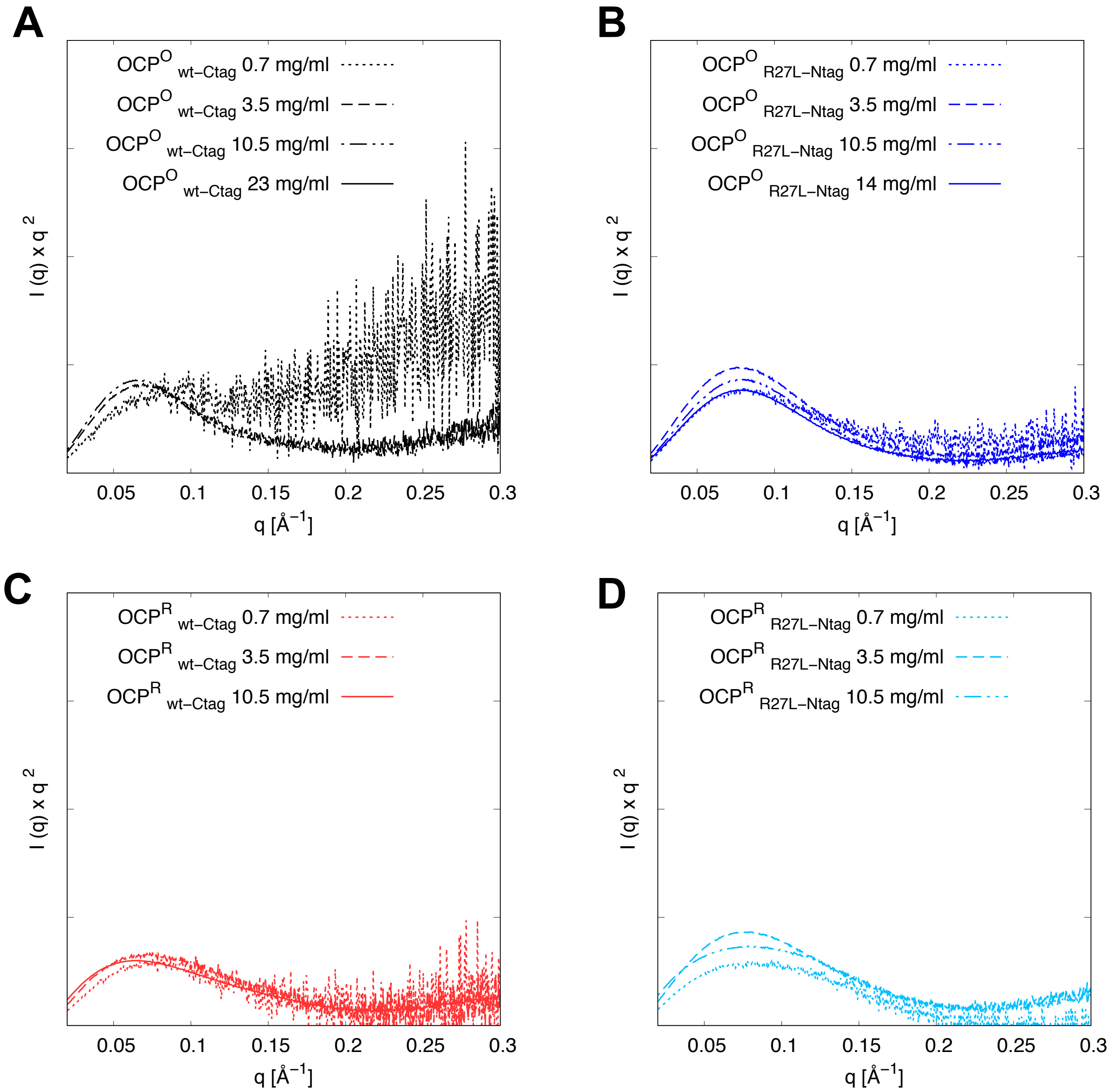
A
OCP^O wt–Ctag 0.7 mg/ml
OCP^O wt–Ctag 3.5 mg/ml
OCP^O wt–Ctag 10.5 mg/ml
OCP^O wt–Ctag 23 mg/ml
I (q) x q 2
0.05
0.1
0.15
0.2
0.25
0.3
q [Å−1]
B
OCP^O R27L–Ntag 0.7 mg/ml
OCP^O R27L–Ntag 3.5 mg/ml
OCP^O R27L–Ntag 10.5 mg/ml
OCP^O R27L–Ntag 14 mg/ml
I (q) x q 2
q [Å−1]
C
OCP^R wt–Ctag 0.7 mg/ml
OCP^R wt–Ctag 3.5 mg/ml
OCP^R wt–Ctag 10.5 mg/ml
I (q) x q 2
q [Å−1]
D
OCP^R R27L–Ntag 0.7 mg/ml
OCP^R R27L–Ntag 3.5 mg/ml
OCP^R R27L–Ntag 10.5 mg/ml
I (q) x q 2
q [Å−1]

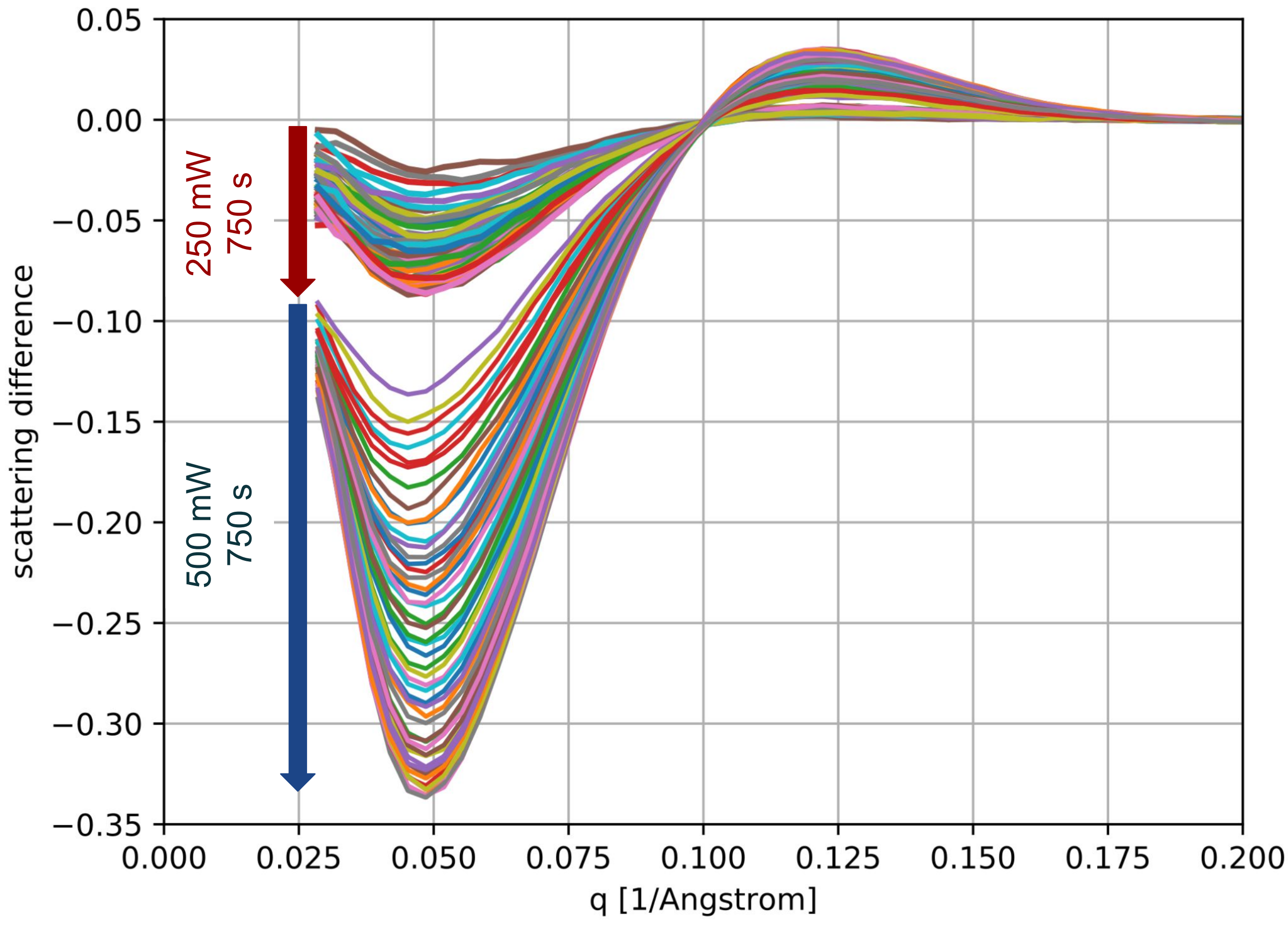

0.05
0.00
−0.05
−0.10
−0.15
−0.20
−0.25
−0.30
−0.35
0.000
0.025
0.050
0.075
0.100
0.125
0.150
0.175
0.200
q [1/Angstrom]
scattering difference
250 mW
750 s
500 mW
750 s

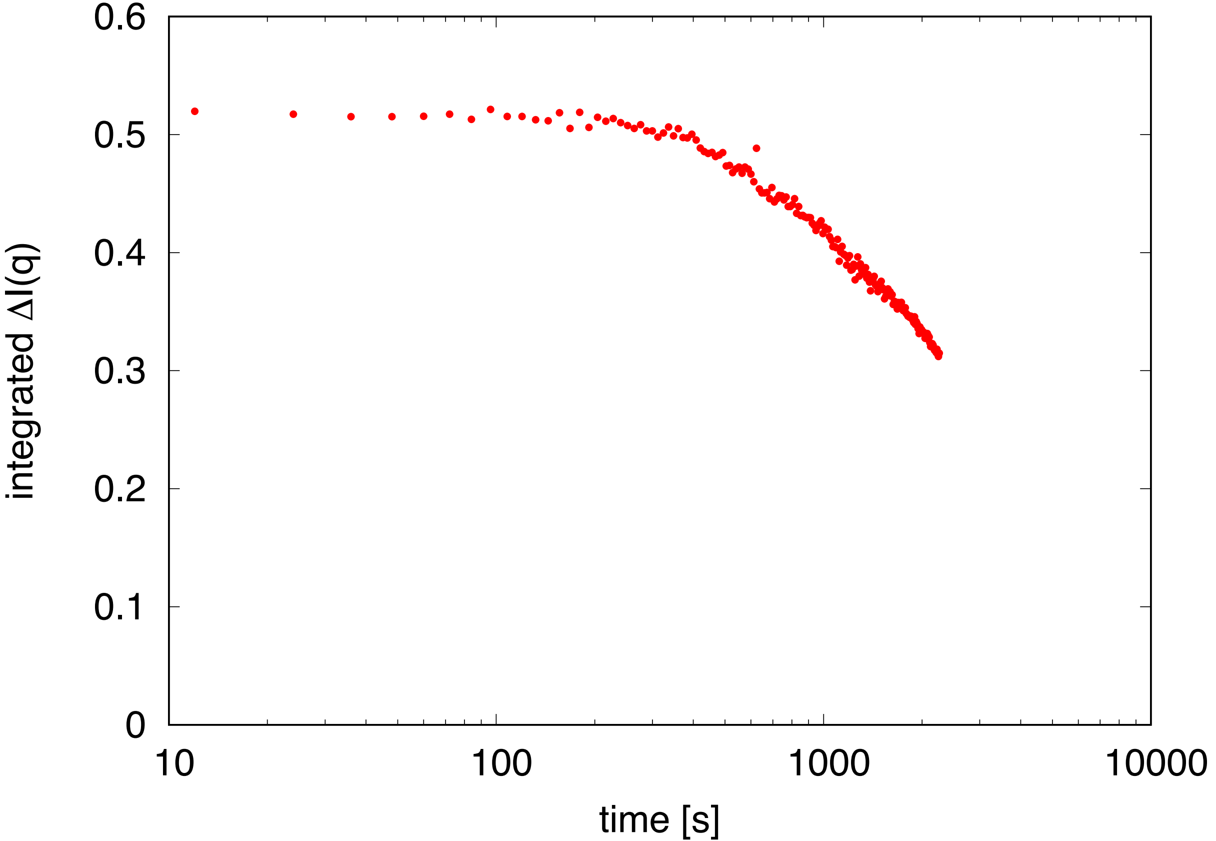
0.6
0.5
0.4
0.3
0.2
0.1
0
integrated ΔI(q)
10
100
1000
10000
time [s]

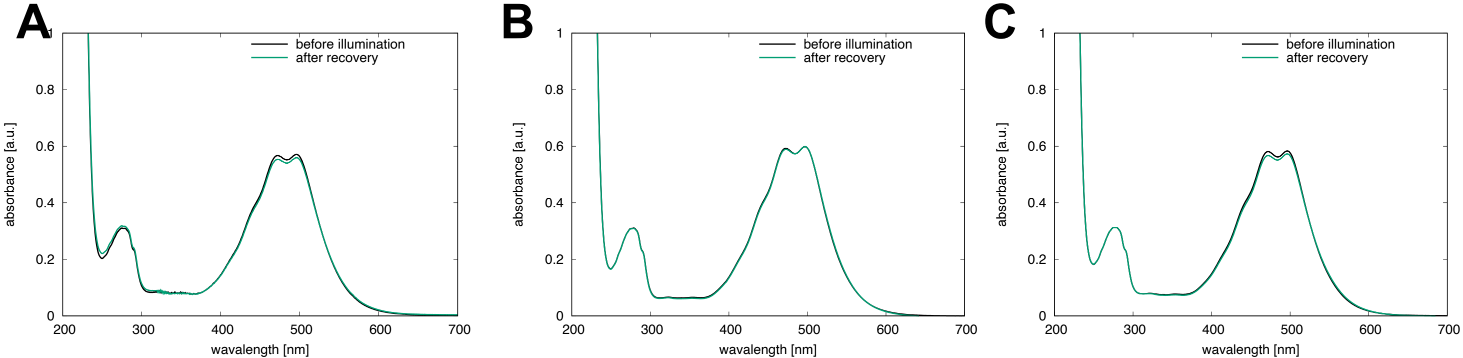

A
B
C
before illumination
after recovery
absorbance [a.u.]
wavalength [nm]
0
0.2
0.4
0.6
0.8
1
200
300
400
500
600
700

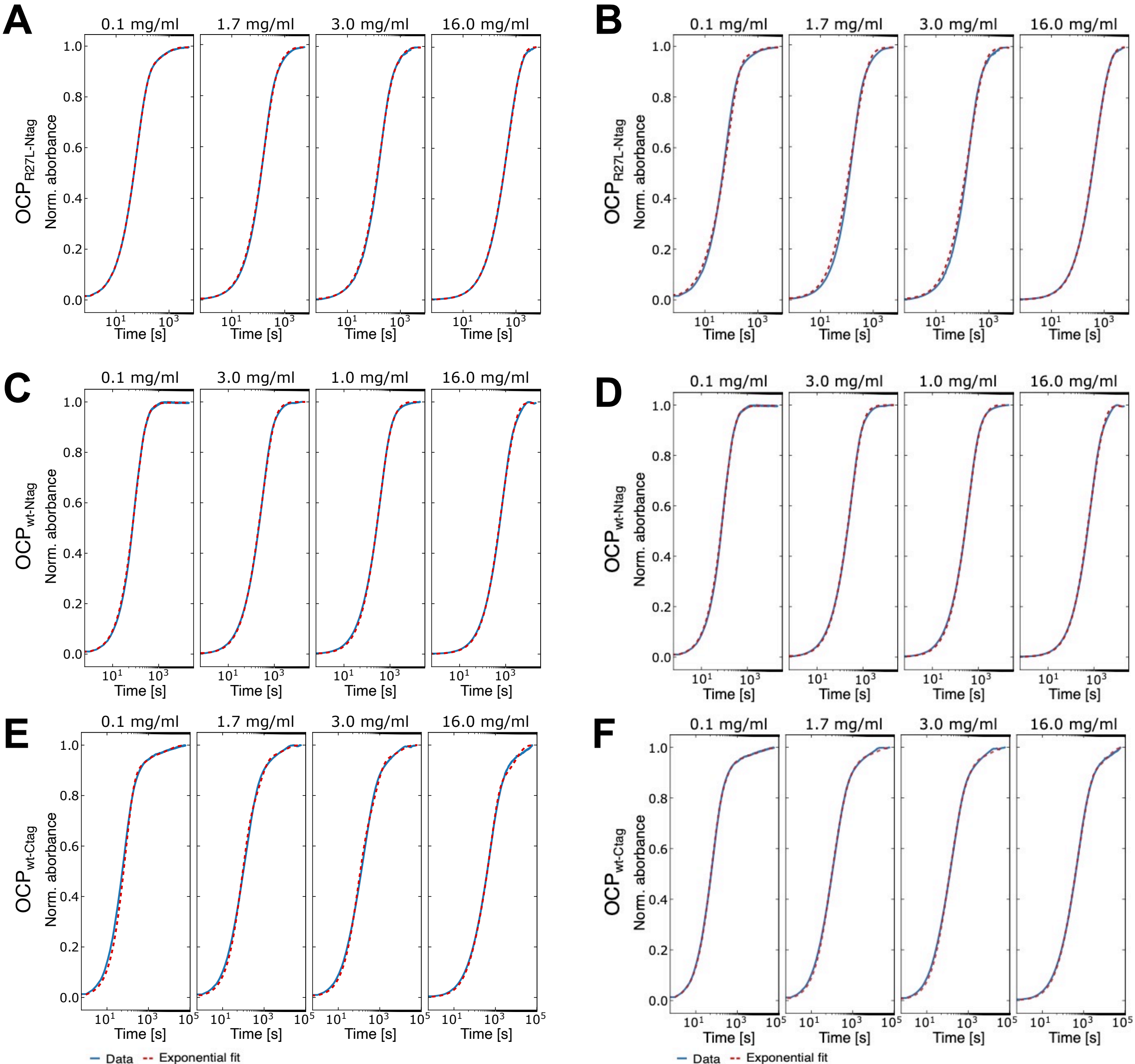
A
B
C
D
E
F
0.1 mg/ml
1.7 mg/ml
3.0 mg/ml
1.0 mg/ml
16.0 mg/ml
OCP$_{R27L-Ntag}$
OCP$_{wt-Ntag}$
OCP$_{wt-Ctag}$
Norm. aborbance
Time [s]
Data
Exponential fit

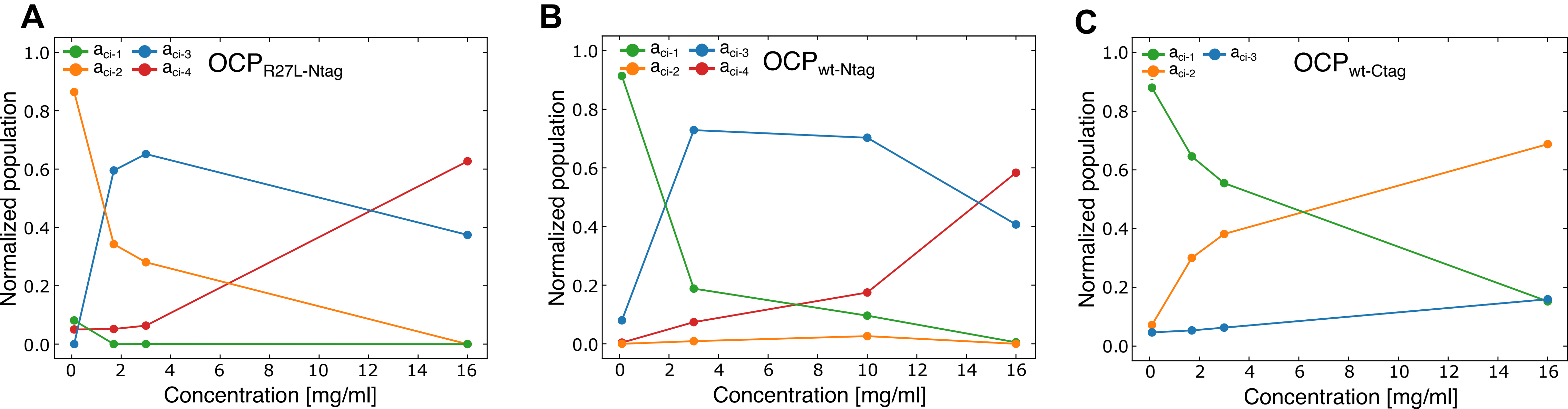

A
B
C
OCP_R27L-Ntag
OCP_wt-Ntag
OCP_wt-Ctag
a_ci-1
a_ci-2
a_ci-3
a_ci-4
Normalized population
Concentration [mg/ml]

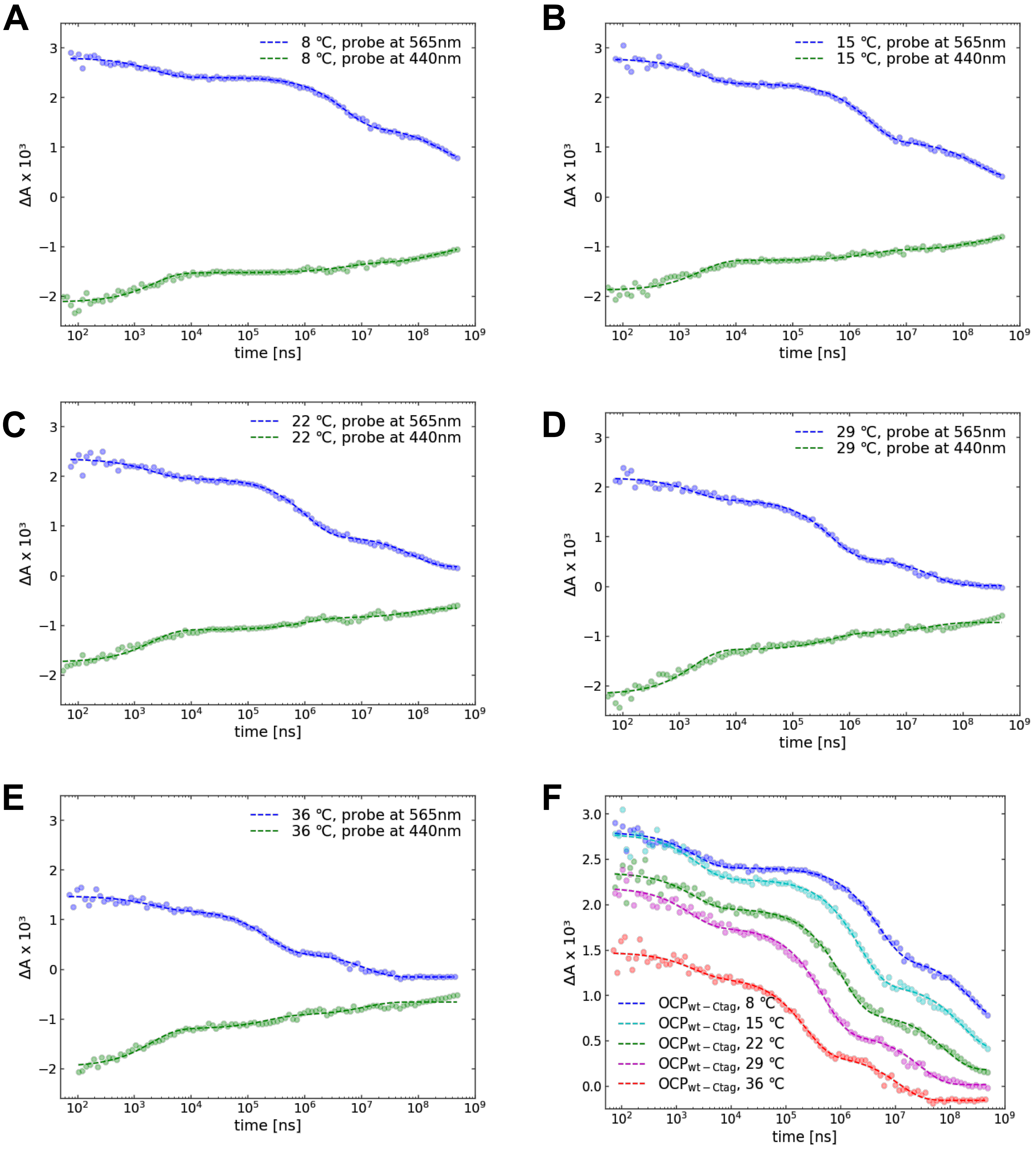
A
8 ℃, probe at 565nm
8 ℃, probe at 440nm
ΔA x 10³
time [ns]
B
15 ℃, probe at 565nm
15 ℃, probe at 440nm
ΔA x 10³
time [ns]
C
22 ℃, probe at 565nm
22 ℃, probe at 440nm
ΔA x 10³
time [ns]
D
29 ℃, probe at 565nm
29 ℃, probe at 440nm
ΔA x 10³
time [ns]
E
36 ℃, probe at 565nm
36 ℃, probe at 440nm
ΔA x 10³
time [ns]
F
OCP$_{wt-Ctag}$, 8 ℃
OCP$_{wt-Ctag}$, 15 ℃
OCP$_{wt-Ctag}$, 22 ℃
OCP$_{wt-Ctag}$, 29 ℃
OCP$_{wt-Ctag}$, 36 ℃
ΔA x 10³
time [ns]

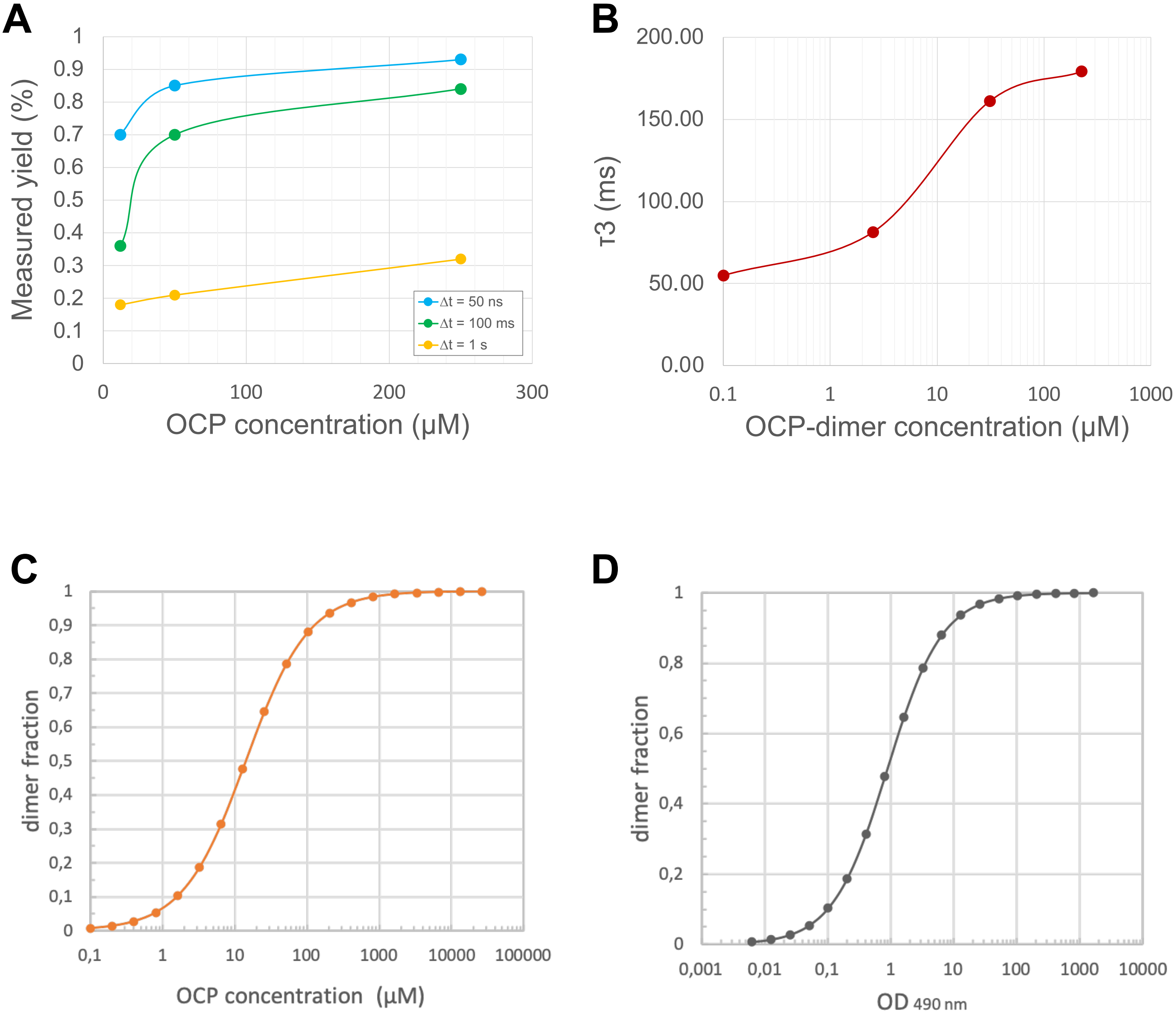

A
Measured yield (%)
OCP concentration (μM)
Δt = 50 ns
Δt = 100 ms
Δt = 1 s
B
τ3 (ms)
OCP-dimer concentration (μM)
C
dimer fraction
OCP concentration (μM)
D
dimer fraction
OD 490 nm

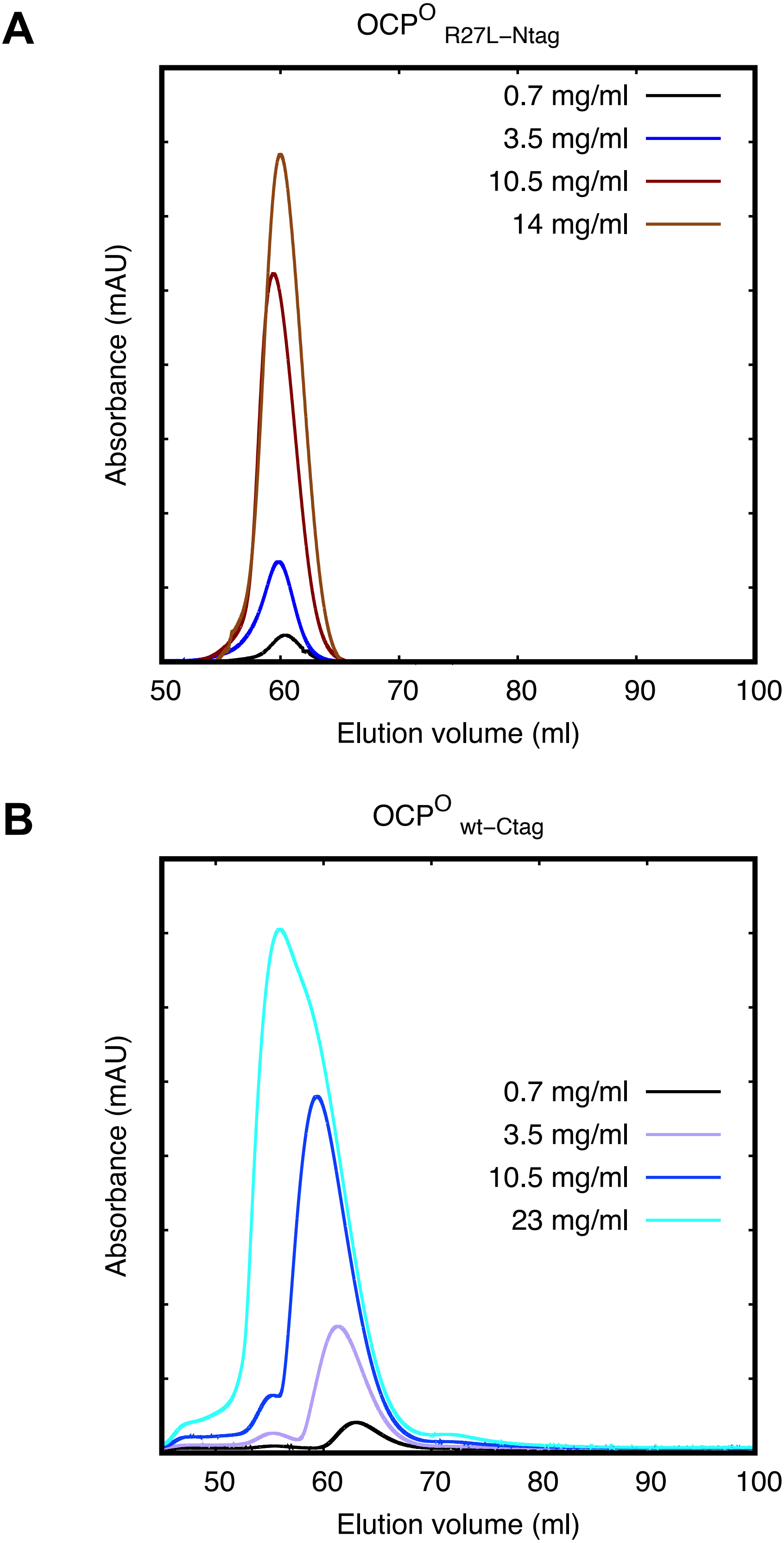
A
OCP^O R27L−Ntag
0.7 mg/ml
3.5 mg/ml
10.5 mg/ml
14 mg/ml
Absorbance (mAU)
50
60
70
80
90
100
Elution volume (ml)
B
OCP^O wt−Ctag
0.7 mg/ml
3.5 mg/ml
10.5 mg/ml
23 mg/ml
Absorbance (mAU)
50
60
70
80
90
100
Elution volume (ml)

```
import numpy as np
import pylab as plt
from scipy import optimize
import scipy.optimize
from sys import exit

np.set_printoptions(threshold=np.inf)
datafile = ' '
f = open(datafile,"r")
data = f.readlines()
f.close()
time = data[0].split()[1:]
# ----------- read data and calculate SVD
matrix_plot = np.loadtxt(datafile)
q = matrix_plot.T[0]
matrix = np.delete(matrix_plot,[0],1)
rank_full = len(matrix[0])
U,s,V = np.linalg.svd(matrix,0)
S = np.diag(s)
br =  # basis rank to be plotted
# ----------- plot basis patterns
fig1 = plt.figure()
fig1.suptitle("basis patterns amplitude vectors")
for (ind,u) in zip(range(br),np.dot(U,S).T):
 plt.subplot(br,2,2*ind+1)
 plt.plot(q,u)
 plt.grid()
plt.xlabel("q / A^-1")
plt.show(block=False)
open(' ', "w").close()
q_array = np.array([q])
q_basis = np.concatenate((q_array.T,np.dot(U,S)),axis=1)
np.savetxt(' ',q_basis,delimiter="\t",newline="\n")
# ----------- plot amplitude vectors
for (ind,v) in zip(range(br),V):
 plt.subplot(br,2,2*ind+2)
 plt.plot(time,v,"o-")
 plt.xscale('log')
 plt.grid()
plt.xlabel("time / sec")
plt.show(block=False)
# ----------- plot singular values
fig2 = plt.figure()
fig2.suptitle("singular values")
plt.plot(np.arange(1,rank_full+1),s,"o-")
plt.axis([0,20,0,0.1])
plt.grid()
plt.show(block=False)
plt.xlabel("matrix index")
plt.show(block=False)
A1 = np.dot(U,S)
A2 = np.dot(A1,V)
# ----------- calculate and plot U and V autocorrelations
fig3 = plt.figure()
fig3.suptitle("U and V autocorrelations")
u_corr = [np.sum(U.T[i][0:len(U)-1]*U.T[i][1:len(U)]) for i in range(len(U.T))]
plt.subplot(211)
plt.plot(np.arange(1,rank_full+1),u_corr,"o-")
plt.grid()
v_corr = [np.sum(V[i][0:len(V)-1]*V[i][1:len(V)]) for i in range(len(V))]
plt.subplot(212)
plt.plot(np.arange(1,rank_full+1),v_corr,"o-")
plt.xlabel("matrix index")
plt.grid()
plt.show(block=False)
# ----------- reconstruct data with n. #rank basis patterns
data1 = np.loadtxt(datafile)
rank =   #rank
U3 = U[:,0:rank]
S3 = S[0:rank,0:rank]
V3 = V[0:rank,:]
A3a = np.dot(U3,S3)
A3b = np.dot(A3a,V3)
fig4 = plt.figure()
fig4.suptitle("reduced rank representation")
for j in np.arange(rank_full):
 plt.plot(data1.transpose()[0],data1.transpose()[j+1]*data1.transpose()[0] - 0.0001*j,"bo", ms = 5)
 plt.plot(data1.transpose()[0],A3b.transpose()[j]*data1.transpose()[0] - 0.0001*j,"r-", linewidth = 3)
plt.axis([0.02,0.3,-0.0005*rank_full-0.0002,0.0002])
plt.yticks(np.arange(-0.0005*rank_full,0.0002,0.0005),([''] + time[::-1] + ['seconds']))
plt.grid()
plt.xlabel("q / $\AA^{-1}$")
plt.ylabel("$\Delta$S x q / $\AA^{-1}$")
plt.show(block=False)
open(' ', "w").close()
svd_reconstr = np.concatenate((q_array.T,A3b),axis=1)
np.savetxt(' ',svd_reconstr,delimiter="\t",newline="\n")
```
